# Sexual reproduction impacts vertebrate organ aging

**DOI:** 10.64898/2026.10.08.757363

**Authors:** Jingxun Chen, Aleksandra Tsenter, Emma K. Costa, Doudou Yu, Xiangping Lin, Kenneth L. Chen, Veronica A. Bot, Madeline J. Housh, Léo-Paul Héraud, S.V. Perry, Rishad C. Khondker, Michael P. Snyder, Ashley E. Webb, Tony Wyss-Coray, Anne Brunet

## Abstract

Sexual reproduction is a central component of life. Yet, an unanswered question remains: how does sexual reproduction shape animal physiology and health across an individual’s lifetime? Unlike other external factors such as diet and exercise, the impact of sexual reproduction on organ health during aging is far less studied, particularly in vertebrates. Here, we develop the short-lived killifish as a vertebrate model to study how lifelong mating affects organ aging. We discover that mating regulates liver aging in a cell-type and sex-specific manner. In old female livers, mating dampened hepatocyte inflammation and reduced myeloid cell levels, and machine-learning models predicted that mating led to a ‘younger’ transcriptomic age in several liver cell types. Functionally, mating reduced hepatic accumulation of several lipid classes and was associated with enhanced female resilience against the liver toxin acetaminophen. More broadly, mating induced anti-inflammatory and metabolic rewiring transcriptomic signatures in the brain and gut, indicating that mating may affect vertebrate organ aging more globally. Using publicly available human datasets and plasma proteomics aging clocks, we observed an association between sexual experience and a ‘younger’ plasma proteomic age of several cell types, including hepatocytes. Our work provides evidence that lifelong mating reshapes the physiological context during aging in a vertebrate, highlighting an understudied connection between vertebrate sexual history, health, and aging.

## Introduction

Sexual reproduction is essential for the production of progeny in many species. A fundamental question in biology is how sexual reproduction shapes animal physiology and health over an individual’s lifetime. Studies in invertebrate model species, such as *C. elegans* and *Drosophila*, have provided foundational understanding of this question, revealing that mating can alter organ homeostasis (*1–5*), metabolism (*2, 5–12*), infection susceptibility (*13–16*), and even lifespan (*17–27*). But the impact of sexual reproduction on organ health is less extensively studied in vertebrates. Importantly, vertebrates have evolved specialized organs and systems (*28–30*), including adaptive immunity and specialized immune cell types, such as macrophages and neutrophils (*31, 32*). Therefore, sexual reproduction may affect vertebrate health and aging in a manner distinct from what is observed in invertebrate models.

In vertebrates, mounting evidence indicates that sexual reproduction also influences physiology and lifespan. For example, sexually active male rats have elevated adrenal and gonadal hormone levels, increased neurogenesis in the hippocampus, and reduced anxiety-like behaviors compared to unmated rats (*33–38*). Female mice with different reproductive histories exhibit differences in lifespans (*39*). Emerging evidence in humans suggests that sexual activity can be associated with a range of health outcomes, including both beneficial and adverse effects (*40–45*). Understanding the mating-aging interactions in a vertebrate model is critical to establishing a vertebrate paradigm that informs our understanding of how human sexual history may relate to health, aging, and disease. Yet, a challenge in studying mating-aging interactions in vertebrates is the long lifespan of traditional vertebrate model systems, such as mice, which typically live 2-3 years.

The African turquoise killifish (*Nothobranchius furzeri*) is a promising model for examining how sexual reproduction impacts organ aging in vertebrates. The killifish is a naturally short-lived vertebrate, with a median lifespan of 4-6 months (about a fifth of the mouse median lifespan and a seventh of the zebrafish median lifespan) (*30, 46–53*). Recent developments in genetic and ‘omic’ resources for the killifish (*54–63*) have revealed conserved aging mechanisms and interventions (*64–73*). Importantly, it has vertebrate-specific organs and systems (e.g., adaptive immunity) that are also present in humans but absent in invertebrates (*30, 47, 49, 50, 53*). The killifish reproduces sexually via external fertilization, and embryos also develop externally without parental care. While these reproductive features differ from those of mammals, they enable us to focus on specific aspects of mating (e.g., the physical act of mating, gamete production, and pheromone sensing), which are difficult to disentangle from pregnancy and parenting in mammalian models.

In this study, we developed a lifelong mating paradigm in killifish to uncover the effects of mating on organ aging. Our work revealed that mating affected liver aging in a cell-type and sex-specific manner. In old females, mating dampened inflammatory signatures in the liver and led to a ‘younger’ transcriptome in several liver cell types. Functionally, mating altered liver lipid profiles and was associated with enhanced detoxification resilience in old females. The mating effect on immune cell regulation in female livers was diminished in a mutant that produced infertile eggs. More generally, mating induced anti-inflammatory and metabolic rewiring transcriptomic signatures in the female brain and gut, suggesting possible organism-level effects.

Using publicly available human plasma proteomics datasets and plasma proteomics aging clocks, we found an association between sexual experience in women with ‘younger’ plasma proteomic age specific to several cell types.

## Results

### Mating ‘rejuvenates’ liver transcriptomes in a sex- and cell-type-specific manner

To assess the long-term effects of mating on killifish aging, we developed a lifelong mating paradigm for the killifish. In this paradigm, diet and mating duration, timing, and frequency were carefully controlled throughout an individual’s entire life (**Fig. 1A**, see Methods for details). Fish underwent four-hour mating sessions twice per week, from the onset of sexual maturity until death. During the mating session, the male killifish courts the female multiple times and presses against her belly to stimulate the release of eggs onto a sandy surface, where the eggs are externally fertilized by his sperm (**fig. S1A**). The fertilized eggs then develop externally without parental care. In our experiments, a mate pair consisted of a single pair of male and female killifish (**Fig. 1A**, ‘Mated’). As a control, the age-matched unmated fish were also subjected to the same fish-handling and environmental changes but remained lifelong virgins (**Fig. 1A**, ‘Unmated’). Under this paradigm, unmated females appeared to be healthy overall and did not exhibit a pathologic ‘egg-bound’ phenotype sometimes reported in unmated females of fish species (*74, 75*) (**fig. S1B**). Mated killifish females in this paradigm were fertile (**Fig. 1B**), and they also exhibited a decline in the number of eggs laid with age (**Fig. 1B**), consistent with a previous report using a different mating paradigm (*76*).

**Fig. 1.**
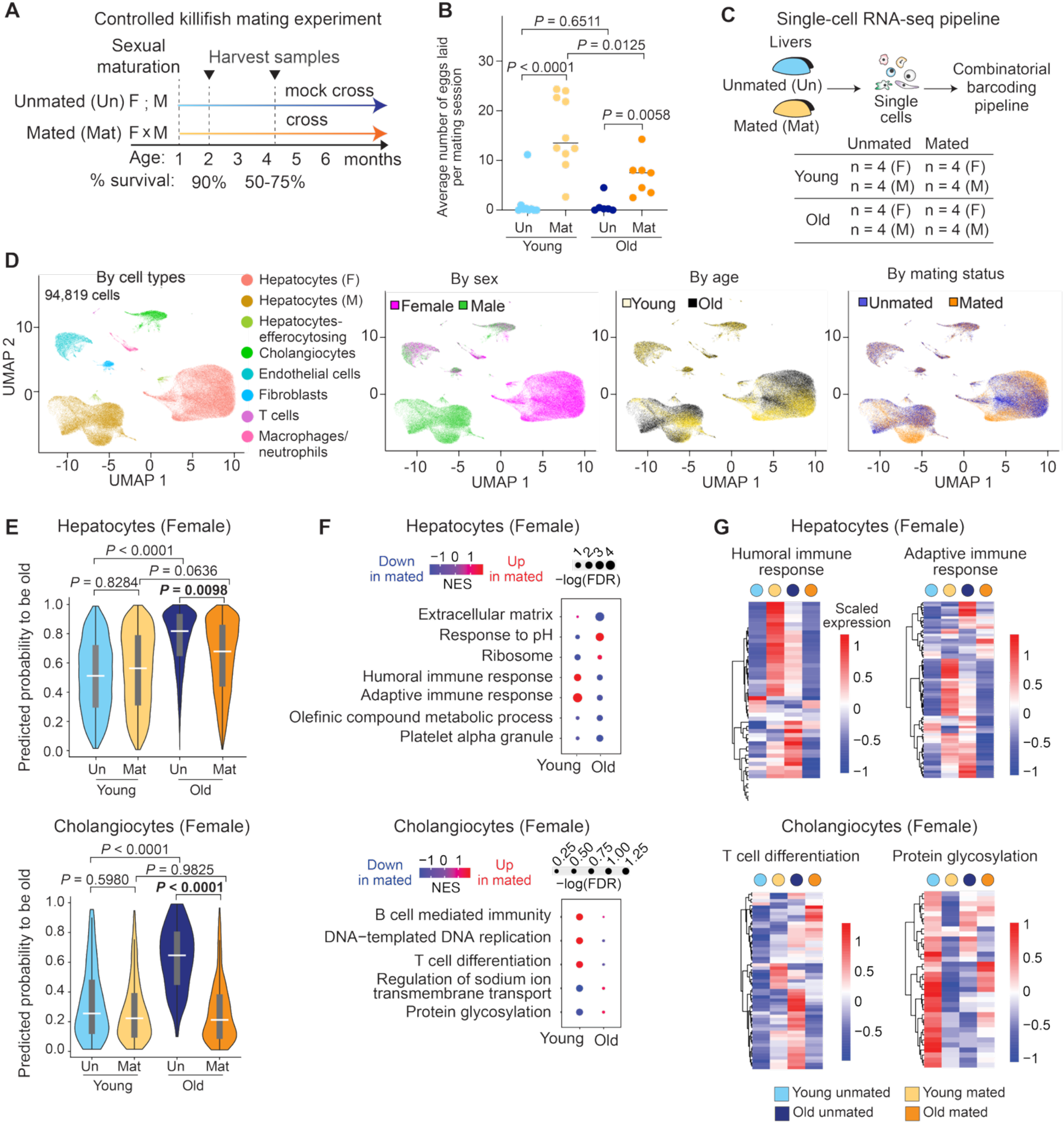
Mating ‘rejuvenates’ liver transcriptomes in a sex- and cell-type-specific manner. (A) Schematic for the lifelong killifish mating experiment. Male and female killifish were paired at sexual maturation and mated regularly throughout life. Unmated killifish were subjected to the same fish-handling and environmental change as the mated fish (‘mock cross’). Samples were harvested at 24 hours after mating and at ∼2 (young) and 4.5 (old) months of age, corresponding to ∼90% and 50-75% survival of the cohorts, respectively. F, female; M, male. (B) Average number of eggs laid per mating session for mated or unmated females at young or old age. Each dot represents an individual fish (young unmated: 9 fish; young mated: 10 fish; old unmated: 6 fish; old mated: 7 fish), and the bar is the median of all fish. The *P* values were determined using the Mann-Whitney *U* test for pairwise comparisons. Un, unmated; Mat, mated. (C) Schematic for the single-cell RNA-seq pipeline. The livers from unmated (‘Un’) or mated (‘Mat’) animals were dissociated into single-cell suspensions on ice, fixed, and processed together using a multiplex split-pool combinatorial barcoding method. Four animals were used for each condition (animals were collected from 2-3 independent cohorts), and eight conditions were studied, including all combinations of sex (male or female), age (young or old), and mating status (unmated or mated). F, female; M, male. (D) UMAP colored by various factors, including cell types (left), sex (middle left), age (middle right), and mating status (right). (E) Distribution of each cell’s CellBiAge predicted probability of being ‘old,’ plotted for selected female cell types—hepatocytes (top) and cholangiocytes (bottom). Bar represents the median of all cells (grouping all fish of each condition), and the quantiles 1 (Q1) and 3 (Q3) are the 25^th^ and 75^th^ percentiles, respectively. The *P* values were determined using a linear mixed-effects model for all pairwise comparisons, except the comparison between unmated young vs. unmated old, where a Welch’s t-test was used. Un, unmated; Mat, mated. (F) Gene set enrichment analysis (GSEA) results for selected female cell types (hepatocytes, top; cholangiocytes, bottom), identifying the pathways significantly enriched for the genes upregulated or downregulated by mating in young or old females. NES, normalized enrichment score. Dot size, –log10 of the false discovery rate (FDR). (G) Heatmaps of the genes that drive each selected gene ontology (GO) term. Each gene’s pseudobulked and DESeq2-normalized expression by cell type (hepatocytes, top; cholangiocytes, bottom) was averaged across four animals per condition and z-scaled.

To understand how lifelong mating affects organ aging, particularly vertebrate-specific organs, we first examined the liver. It is a vertebrate-specific organ (*28–30*) with versatile functions, including nutrient sensing, metabolism, endocrine signaling, and detoxification (*77–80*). Liver health is tightly linked to reproduction in vertebrates, including humans. For example, liver diseases (e.g., Metabolic Dysfunction-Associated Steatotic Liver Disease, previously known as Non-Alcoholic Fatty Liver Disease) are associated with reduced fertility (*81–84*); reciprocally, reproductive disorders (e.g., Polyendocrine Metabolic Ovarian Syndrome, previously known as Polycystic Ovary Syndrome) are linked to liver dysfunction (*85–89*). These features prompted us first to analyze how liver aging is affected by lifelong mating.

We systematically characterized how lifelong mating affected liver aging at a single-cell resolution. To this end, we used a cohort of mated or unmated males and females at two different ages (young: 2 months, corresponding to 90% survival of the cohort; and old: 4-5 months, corresponding to 50-75% survival) (**Fig. 1C**). Using an optimized, multiplexed single-cell RNA-sequencing (scRNA-seq) approach (**Fig. 1C**; see Methods), we generated a high-quality dataset comprising four biological replicates per condition (n = 4, each condition corresponding to a different combination of sex, age, and mating status), with >85% of cells containing <5% mitochondrial reads and ∼95,000 high-quality cells overall. Uniform Manifold Approximation and Projection (UMAP) of the dataset revealed distinct cell clusters corresponding to key liver cell types, including hepatocytes, cholangiocytes, endothelial cells, fibroblasts, and immune cells, such as T cells and macrophages/neutrophils (**Fig. 1D**). We found that hepatocytes were the most sex-dimorphic among the liver cell types (**Fig. 1D**, ‘By sex’), consistent with previous reports in killifish and mice (*61, 90, 91*).

Given the strong sexual dimorphism in hepatocyte transcriptomes, we kept the male and female hepatocytes as distinct cell clusters in subsequent analyses. After pseudobulking by cell type and condition to increase the robustness of the analysis (*92, 93*), variance partitioning analysis revealed that, for both sexes, age had the strongest impact on hepatocyte transcriptomes among all cell types (**fig. S1C**). In females, mating also had the strongest impact on hepatocyte transcriptomes among all cell types (**fig. S1C**). This result was supported by UMAP, where cells were separated by age and mating status within hepatocyte clusters in both sexes (**Fig. 1D**, ‘By age’ and ‘By mating status’) but less obviously in other cell types (e.g., endothelial cells). Therefore, hepatocyte transcriptomes not only differed between the sexes but were also particularly affected by aging and mating.

Does mating impact the transcriptomes of male and female hepatocytes to the same degree? Principal Component Analysis (PCA) revealed that for female hepatocytes, PC2 (15% variance explained) separated the samples by age, and PC1 (33% variance explained) separated the samples by mating status within each age group (**fig. S1D**, ‘Hepatocytes (Female)’). In contrast, for male hepatocytes, there was clear separation by age in PC2 (23% variance explained) but not by mating status in PC1 (30% variance explained) (**fig. S1D**, ‘Hepatocytes (Male)’). Consistently, compared to male hepatocytes, female hepatocytes had more genes differentially expressed by mating status at both young and old ages (**fig. S1E**). These analyses suggested that, at least at the transcriptome level, mating impacted female hepatocytes more strongly than male hepatocytes.

What are the main effects of mating on the ‘transcriptomic age’ of liver cell types – does mating exacerbate or rejuvenate aging signatures? We applied the CellBiAge machine learning model (*94*) to unbiasedly evaluate the transcriptomic age signatures of liver cell types. We trained the model on the unmated young and old samples, built a classifier to determine the probability of a cell being ‘old,’ and applied this classifier to individual cells in the mated samples (cells that were not used to build the model) (**fig. S2A**; see Methods for details and Supplementary Text for quality-control results of the classifier). We observed a significantly greater proportion of ‘younger’ hepatocytes, cholangiocytes, and endothelial cells in mated old females than in unmated females, suggesting that mating leads to ‘transcriptomic rejuvenation’ in several liver cell types (**Fig. 1E** and **fig. S2C**). Independently, we correlated the hepatocyte gene expression changes due to aging (**fig. S2B**, ‘Average Log_2_FC of old unmated vs. young unmated’) against those due to mating at old age (**fig. S2B**, ‘Average Log_2_FC of old mated vs. old unmated’). This analysis showed that most age-dependent changes were reversed in direction for mated old females (**fig. S2B**, ‘Hepatocytes (Female)’, Q2 and Q4), corroborating that mating ‘reverses’ transcriptional aging signatures in female hepatocytes.

For males, neither CellBiAge nor correlation analysis clearly revealed a transcriptomic ‘rejuvenating’ effect of mating for any liver cell type (**fig. S2B**, ‘Hepatocytes (Male),’ and **fig. S2C**; see Supplementary Text for extended analysis). These findings are consistent with a weaker effect of mating on male transcriptomes, as shown by variance analysis and PCA (**fig. S1C** and **S1D**).

We next identified the main pathways that may explain this transcriptomic ‘rejuvenation,’ particularly in female hepatocytes at old age. Gene Set Enrichment Analysis (GSEA) revealed that immune-related pathways were downregulated by mating at old age, including ‘humoral immune response’ and ‘adaptive immune response’ (**Fig. 1F** and **1G**). Consistently, for several age-altered pathways, mated females had reverse directions of change compared to unmated females, including those related to immune regulation, such as ‘immune effector process,’ ‘regulation of macrophage chemotaxis,’ and ‘C-C chemokine receptor activity’ (**fig. S2D**). These data suggest that mating reduces several immune-related signatures at old age, raising the possibility that it might dampen liver inflammation at old age (see below for independent validation).

Together, these findings revealed that lifelong mating affects killifish livers in ways specific to biological sex and cell types and may induce transcriptomic ‘rejuvenation’ in female liver cell types, such as hepatocytes.

### Mating dampens innate immune cell levels and inflammation in old female livers

Next, we independently validated whether lifelong mating dampens age-dependent inflammation characteristics in female livers. Age-related increases in immune cell levels have been observed in killifish and mammalian organs (*66, 95–99*). Thus, we performed RNA *in situ* hybridization to quantify immune cell abundance and pro-inflammatory marker expression in liver tissue sections. We generated independent mating cohorts to compare unmated and mated males and females at young (2-2.5 months old) and old (4.5 months old) ages (**Fig. 2A**). First, we quantified the proportion of immune cells, particularly myeloid immune cells, including macrophages and neutrophils. For both unmated and mated female livers, the percentage of cells that were *CD45+* leukocytes per section significantly increased with age (**Fig. 2B**, *P* = 0.0262 for unmated and *P* = 0.0041 for mated), consistent with increased immune cell abundance with age in mammalian livers (*97–99*). Interestingly, lifelong mating led to fewer leukocytes (*P* = 0.0146), macrophages (*P* = 0.0379), and neutrophils (*P* = 0.0128) in female livers at old age (**Fig. 2B** and **fig. S3A**), a trend also observed by scRNA-seq (**fig. S2E**). A trending decrease in *CD45*+ leukocytes and macrophages in the liver was also observed in mated young females (*P* = 0.1096 and *P* = 0.0221, respectively) (**Fig. 2B**). At old age, lifelong mating also led to a significantly reduced percentage of *CD45+* leukocytes expressing the pro-inflammatory marker *TNFα* (**Fig. 2C** and **fig. S3B**, *P* = 0.0436), which marks pro-inflammatory M1 macrophages in zebrafish and in mice (*100–103*). The reduction in immune cell levels, particularly leukocytes with pro-inflammatory markers, suggests that lifelong mating dampens immune cell levels and their inflammation status in old female livers.

**Fig. 2.**
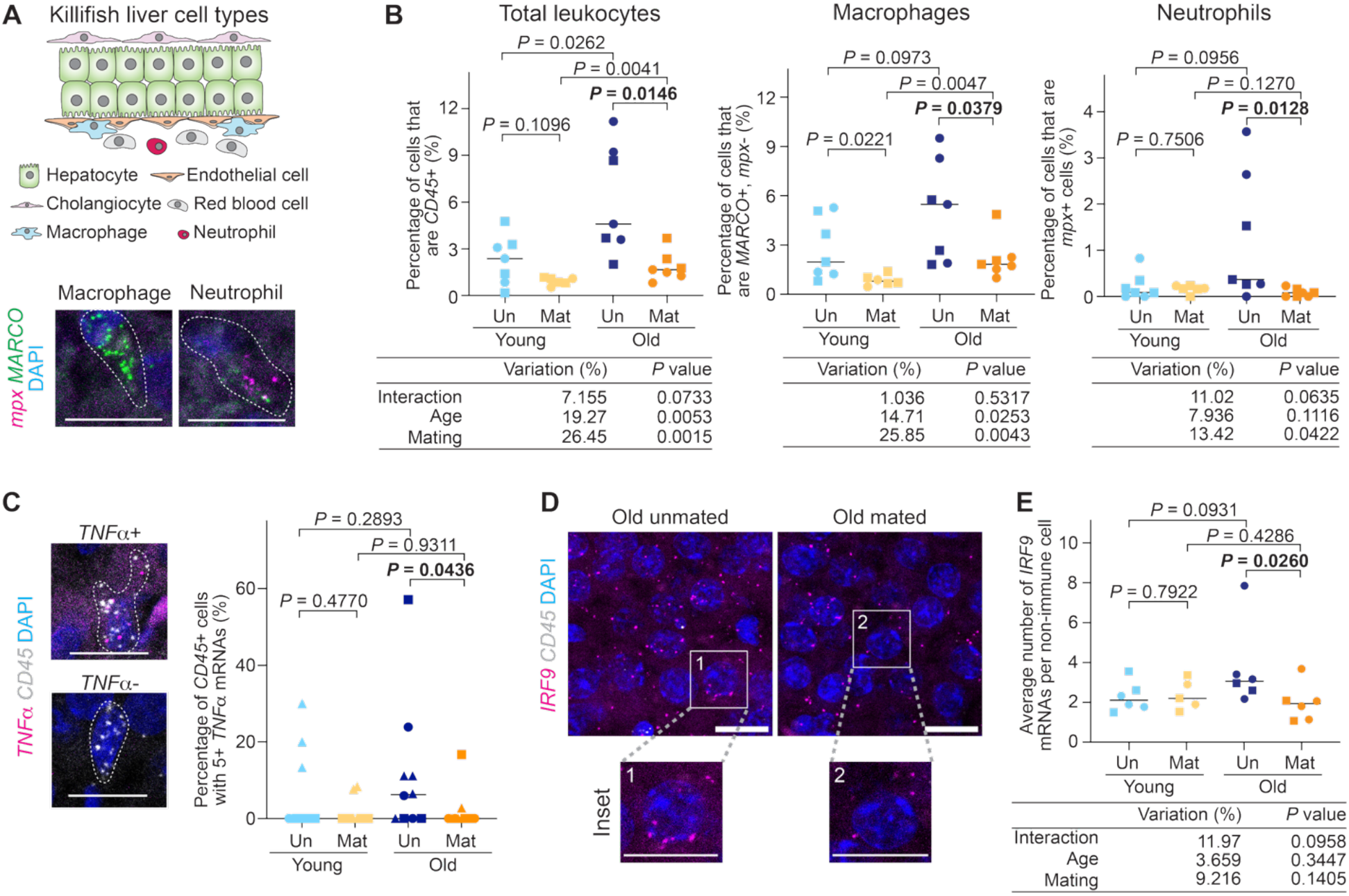
Mating dampens immune cell levels and inflammation in old female livers. (A) Schematic for liver cell types in the killifish and example RNA *in situ* images for a macrophage (*MARCO*+, *mpx*-) and a neutrophil (*mpx+*), after staining liver tissue sections for *mpx* (magenta) and *MARCO* (green) mRNAs and nuclei by DAPI (blue). Dashed lines outline cells. Scale bar, 10 *µm*. Note: *mpx* is the zebrafish ortholog name (killifish name: *LOC107372911*). (B) Proportion of different immune cells in female livers, including total leukocytes (*CD45*+), macrophages (*MARCO*+, *mpx*-), and neutrophils (*mpx*+). Each dot represents an individual fish (young unmated, n = 7 fish; young mated, n = 6 fish; old unmated, n = 7 fish; old mated, n = 7 fish), and the bar is the median of all fish per condition. Symbol shape, cohort number (animals were raised in 2 cohorts independent of the scRNA-seq cohorts). Hereinafter, in this figure, young, 2-2.5 months old; old, 4.5 months old. The *P* values were determined using the Mann-Whitney *U* test for pairwise comparisons and a two-way ANOVA to assess the effects of age, mating status, and age-mating interaction. Un, unmated; Mat, mated. (C) Left, example RNA *in situ* images for a *TNFα*+ or a *TNFα*-immune cell (*CD45*+), after staining liver tissue sections for *TNFα* (magenta) and *CD45* (gray) mRNAs and nuclei by DAPI (blue). Dashed lines outline cells. Scale bar, 10 *µm*. Right, quantification of the proportion of *TNFα*+ immune cells in female livers (young unmated, 11 fish; young mated, 10 fish; old unmated, n = 10 fish; old mated, n = 11 fish), and the bar is the median of all fish. Symbol shape, cohort number (the plot includes animals pooled from the cohorts used for scRNA-seq and two additional independent cohorts used in panel B). The graph was plotted, and statistics for pairwise comparisons were performed as in panel B. Note that two-way ANOVA was not performed due to non-normal sample distributions. (D) Example RNA *in situ* images for female liver sections (young unmated, n = 6 fish; young mated, n = 5 fish; old unmated, n = 6 fish; old mated, n = 6 fish), staining for *IRF9* (magenta) and *CD45* (gray) mRNAs and nuclei by DAPI (blue). Scale bar, 10 *µm*. (E) Quantification of panel D, plotting average *IRF9* expression in *CD45*-negative (non-immune) cells of female livers (young unmated, n = 6 fish; young mated, n = 5 fish; old unmated, n = 6 fish; old mated, n = 6 fish), and the bar is the median of all fish per condition. Symbol shape, cohort number (these cohorts were raised independently of the scRNA-seq cohorts, same as those used in panel B). The graph was plotted, and statistics were performed as in panel B.

In contrast, mating had different effects on immune cell balance in male livers. At baseline, male livers had fewer macrophages than female livers (**fig. S3C**), a trend also observed by scRNA-seq (**fig. S2E**). Mating appeared to increase macrophage percentage per section in male livers when both age groups were considered, with a trending increase at each age (**fig. S3C**, ‘Macrophage’, *P* = 0.0337 by two-way ANOVA). This trend was opposite to that in the female livers (**Fig. 2B**, ‘Macrophage’, *P* = 0.0043 by two-way ANOVA), suggesting a sex-dimorphic impact of mating on liver macrophage abundance. Thus, mating distinctly affects liver aging in males and females with respect to immune cell levels.

How does mating affect non-immune cells in the female liver? In *CD45*-negative liver cells, predominantly hepatocytes, we quantified the expression of a pro-inflammatory marker *IRF9*, which has a mammalian ortholog with pro-inflammatory activity (**fig. S4A**) (*104–107*). *IRF9* showed a trending upregulation during liver aging (Spearman’s rank correlation with age: 0.4429; *P* = 0.164) (**fig. S4B,** see Methods for expanded discussion of *IRF9*) and was upregulated upon liver damage in females (see **Fig. 4 below**). We found that at old age, mated females had lower average *IRF9* expression in the liver than unmated females (**Fig. 2D** and **2E**, *P* = 0.0260). This mating-dependent decrease in *IRF9* expression was not seen in young females (**Fig. 2D** and **2E**) (and immune-related pathways, including *IRF9*, were upregulated by mating in young females in our scRNA-seq dataset, possibly due to technical or cohort differences (see **Fig. 1**)). Collectively, our findings supported the idea that, particularly in the livers of old females, lifelong mating dampens inflammation, reducing the levels of pro-inflammatory immune cells and inflammatory markers in non-immune liver cells.

### Mating modulates liver lipid profiles in old males and females

The liver performs several key functions, including metabolism, detoxification, nutrient sensing, and endocrine signaling (*77–80*). Increased liver inflammation is observed in liver aging and pathological conditions in humans, such as Metabolic Dysfunction-Associated Steatotic Liver Disease (*108, 109*) and drug-induced liver injury (*110*). The inflammation dampening by lifelong mating in old female livers raises the question of whether mating is beneficial or detrimental to these physiological functions of the liver.

To directly assess how lifelong mating impacts the liver lipidome, we raised independent mating cohorts and performed untargeted lipidomics on killifish livers along with 23 lipid standards. We profiled mating-associated changes in males and females at young (2-2.5 months old) and old (4.5 months old) ages (**Fig. 3A** and **fig. S5A**). PCA revealed that the quality-control technical replicates were tightly grouped (**fig. S5B, ‘QC’**), and all samples exhibited similar detection depth (**fig. S5C**), indicating high-quality data. We identified 313 uniquely annotated lipids. The lipid profiles of the killifish livers were highly sex dimorphic (PC1, ∼33% variance explained) (**fig. S5B**), with phosphatidylcholines and triglycerides being the most abundant lipid classes for the female and male livers, respectively (**Fig. 3B**). Given this sexual dimorphism, we analyzed each sex separately to understand how mating and aging affected lipid profiles. Within each sex, samples were separated primarily by age in PCA and, to a lesser extent, by mating status, indicating that both aging and mating contributed to global shifts in liver lipid profiles (**Fig. 3C**).

**Fig. 3.**
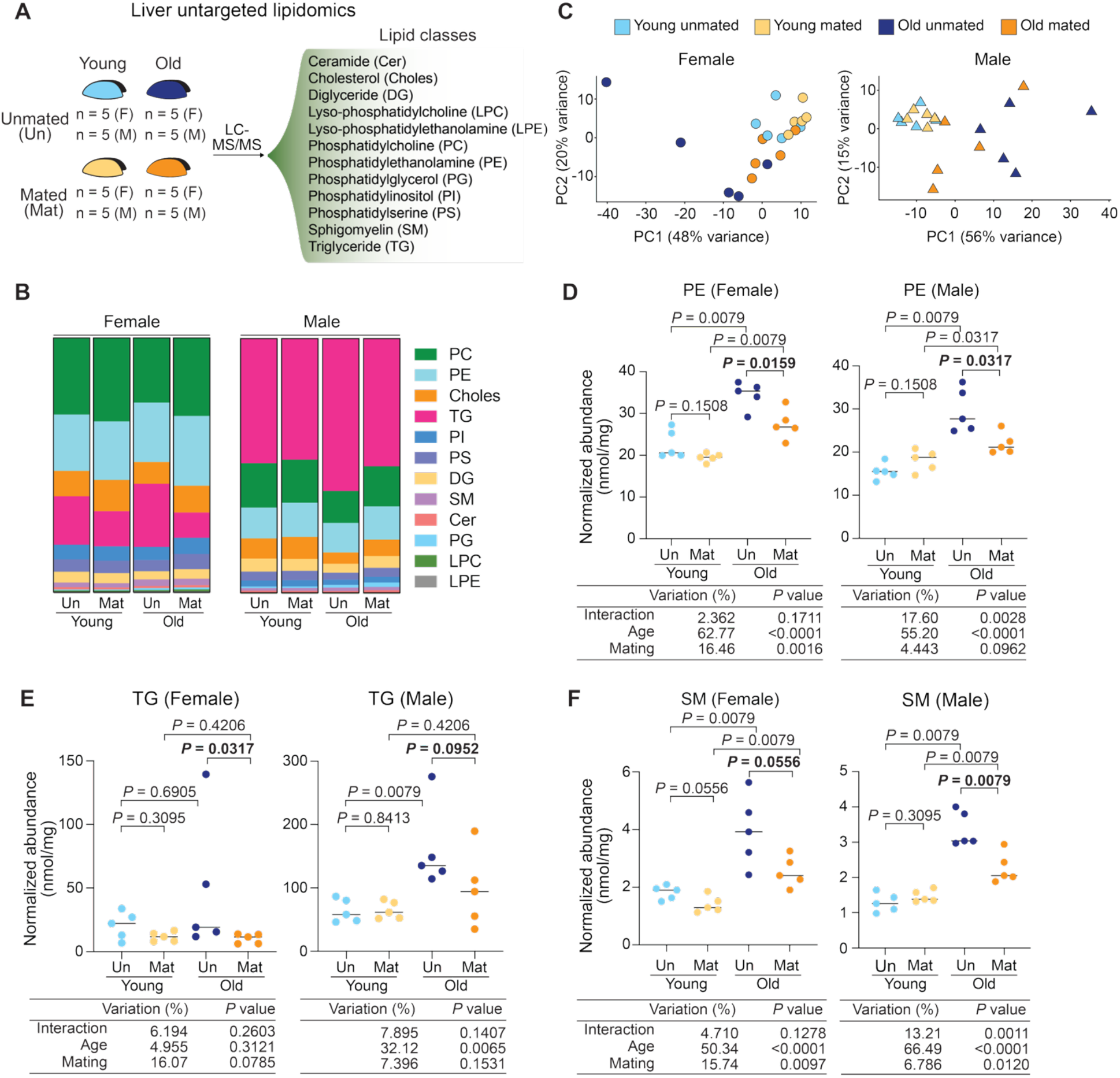
Mating impacts specific lipid class levels in old livers of both sexes. (A) Schematic for liver untargeted lipidomics. The livers from unmated (‘Un’) or mated (‘Mat’) animals were dissected at young (2-2.5 months old) and old (4.5 months old) ages. Five animals were used for each condition (animals were collected from 3 independent cohorts), and eight conditions were used, including all combinations of sex (male or female), age (young or old), and mating status (unmated or mated). F, female; M, male. (B) Proportion of each lipid class for the mated (‘Mat’) and unmated (‘Un’) females and males at each age. The percentage of each lipid class in total lipids, averaged from five fish, was stacked in a column. Each lipid class is color-coded. (C) Principal Component Analysis (PCA) for the estimated concentrations of the 313 uniquely annotated lipids, separately for males and females. Each dot is an animal (n = 5 fish per condition). Circle, female; triangle, male. (D-F) The abundance of the phosphatidylethanolamine (PE, panel D), triglyceride (TG, panel E), and sphingomyelin (SM, panel F) lipid classes in male and female livers. Each dot is a fish (n = 5 per condition), and the bar is the median of the 5 fish in each condition, plotting the normalized abundance summed across all detected lipid species belonging to a given lipid class. Un, unmated; Mat, mated. The *P* values were determined using the Mann-Whitney *U* test for pairwise comparisons and a two-way ANOVA to assess the effects of age, mating status, and the age-mating interaction.

We performed sex-specific differential expression analysis to identify lipid species or lipid classes altered by lifelong mating at either age. Although few individual lipid species were significantly altered by mating at the false discovery rate (FDR) of 0.05 in either sex (**fig. S6A**), we found that mating affected lipid classes as a group. Particularly, mating significantly reduced several lipid classes in old female livers, including phosphatidylethanolamines (‘PE (Female),’ *P* = 0.0159) (**Fig. 3D** and **fig. S6B**) and triglycerides (‘TG (Female),’ *P* = 0.0317) (**Fig. 3E** and **fig. S6B**), as well as two less abundant classes, including phosphatidylinositols (‘PI (Female),’ *P* = 0.0159) (**fig. S6C**) and phosphatidylglycerols (‘PG (Female),’ *P* = 0.0317) (**fig. S6C**). For old male livers, mating also lowered the abundance of phosphatidylethanolamines (PE (Male),’ *P* = 0.0317) (**Fig. 3D** and **fig. S6B)** and phosphatidylinositol (‘PI (Male),’ *P* = 0.0317) (**fig. S6C**), as well as sphingomyelin (‘SM (Male),’ *P* = 0.0079) (**Fig. 3F** and **fig. S6B**). Emerging evidence suggests that elevated levels of some of these lipid classes, such as PE and TG, have been associated with hepatic steatosis and inflammation in mice and with Metabolic Dysfunction-Associated Steatotic Liver Disease in humans (*111–117*). Thus, these shifts in lipid class abundances could be consistent with a mating-related reduction of liver inflammation in old females and reflect a lipid profile often characteristic of a healthy liver, potentially indicating a protective effect of lifelong mating in females.

### Mating is associated with enhanced resilience against drug-induced lethality in old females

In addition to lipid metabolism, the liver has another central function: drug detoxification. The liver acts as a critical filtration system, converting drugs and toxins into inactive or less toxic forms for removal. The most common cause of acute liver failure is drug-induced liver injury (primarily due to overdose of acetaminophen, also known as Tylenol), which accounts for 20-40% acute liver failure cases in the United States and Europe (*118*). To assess how lifelong mating impacts liver resilience against toxins, we developed a killifish liver injury model using acetaminophen overdose. In zebrafish and in mice, acetaminophen-induced lethality and liver injury are dose-dependent (*119–122*). We found that acetaminophen treatment in middle-aged killifish (3 months old) also led to dose-dependent lethality in both sexes (**Fig. 4A**). Sub-lethal acetaminophen treatment increased several pro-inflammatory characteristics in the female liver, such as the percentage of *IL1β*+ cells (*P* = 0.0173) and (albeit to a lesser extent) *TNF*α+ immune cells (*P* = 0.0844) (**fig. S7A**). Sub-lethal acetaminophen treatment also induced the expression levels of two inflammation-related markers in non-immune cells, *IRF9* (*P* = 0.0087) and *TLR5* (*P* = 0.0584) (**Fig. 4B** and **4C**), both of which were also upregulated in mammals exposed to acetaminophen (*110*). These features supported that we successfully established a liver drug-induced injury model in killifish.

**Fig. 4.**
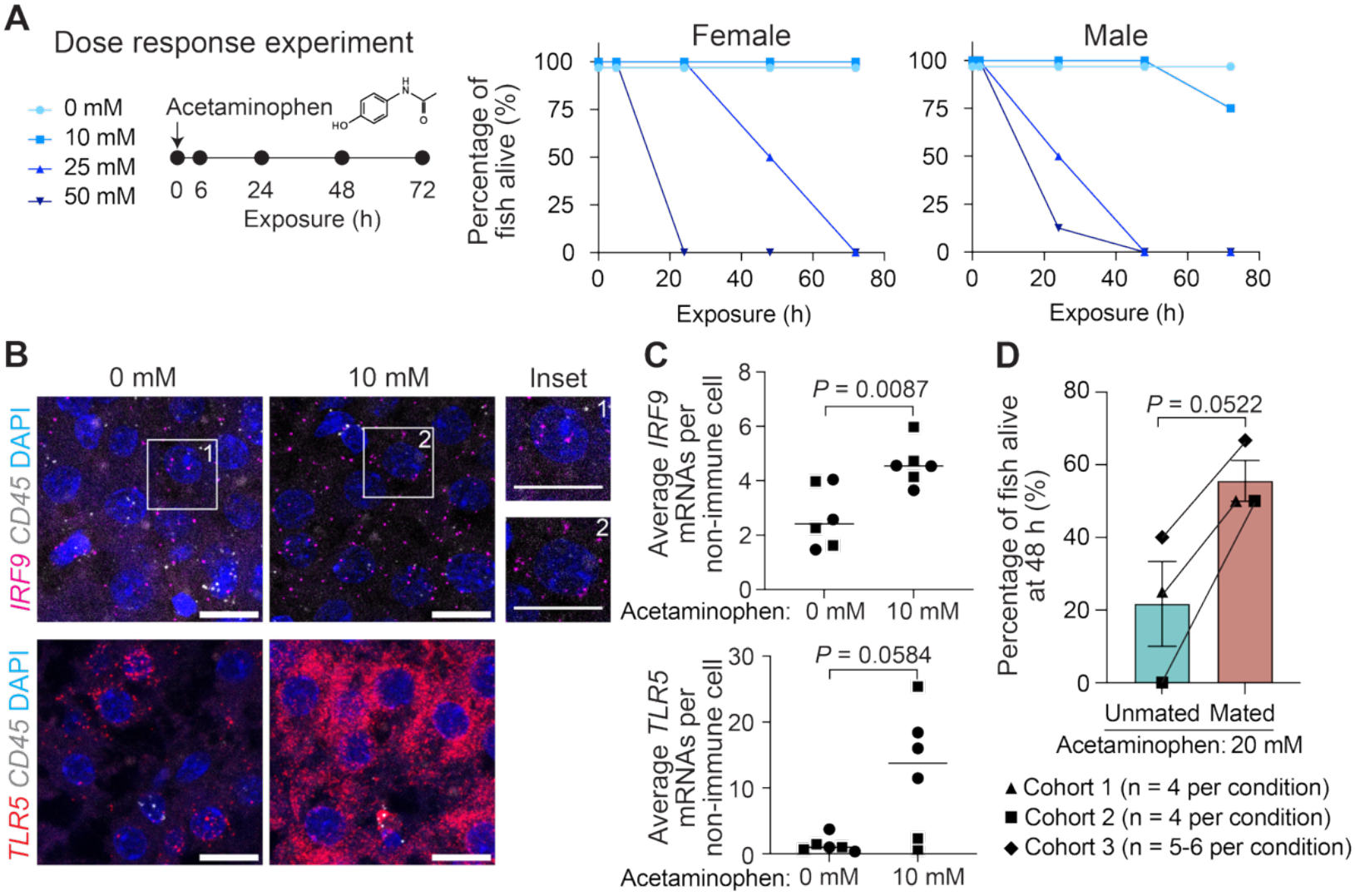
Mating enhances resilience against drug-induced lethality in old females. (A) Dose-response lethality curve for acetaminophen treatment in 3-month-old male and female unmated killifish. Killifish were exposed to various concentrations of acetaminophen dissolved in water for 72 hours. Percentage of fish alive at given time points were plotted for males (0 mM, n = 8 fish; 10 mM, n = 8 fish; 25 mM, n = 8 fish; 50 mM, n = 8 fish at the start of treatment) and females (0 mM, n = 10 fish; 10 mM, n = 6 fish; 25 mM, n = 6 fish; 50 mM, n = 7 fish at the start of treatment). Animals were grouped across 2 or 3 independent cohorts. (B) Example RNA *in situ* images of *IRF9* and *TLR5* expression in the liver cells of unmated 3-month-old females treated with 0 mM (n = 6 fish) or 10 mM (n = 6 fish) acetaminophen for 72 hours. Liver sections were stained for *IRF9* (magenta) or *TLR5* (red) mRNAs, *CD45* (gray) mRNA, and nuclei by DAPI (blue). Scale bar, 10 µm. (C) Quantification of panel B. Top, average *IRF9* mRNAs per *CD45*-negative (non-immune) cell (0 mM, n = 6 fish; 10 mM, n = 6 fish). Bottom, average *TLR5* mRNAs per non-immune cell (normalized to the median of the 0 mM condition of each cohort) (0 mM, n = 6 fish; 10 mM, n = 6 fish). The bar is the median expression of all the fish in each condition. Symbol shape, cohort number (animals were raised in 2 independent cohorts). The *P* values were determined using the Mann-Whitney *U* test for pairwise comparisons. (D) Proportion of unmated (n = 13 fish total at the start of treatment) and mated 4.5-month-old females (n = 14 fish total at the start of treatment) alive after a treatment of 20 mM acetaminophen for 48 hours. Each dot is the survival percentage for a cohort, and the bar height is the average across 3 independent cohorts. The error bars are the standard error of the mean. The *P* value was determined using a paired t-test comparing unmated and mated females treated with 20 mM acetaminophen, matching by cohort number.

To examine how mating affects the response of old female killifish to acetaminophen treatment, we treated mated or unmated old females (3 independent cohorts) with either vehicle control or 20 mM acetaminophen, an intermediary concentration that would allow us to examine potential changes in survival (**Fig. 4A**). At 48 hours after acetaminophen exposure, we observed that mated old females had a survival benefit that was directionally consistent across all three independent cohorts, despite not reaching the significance threshold of 0.05 (**Fig. 4D** and **fig. S7B**, *P* = 0.0522, t-test paired by cohort). These data provide evidence that lifelong mating may be beneficial for liver detoxification in old female killifish.

### Aspects of female fertility may contribute to the effect of mating on liver immune cell levels

Liver health is linked to reproduction in vertebrates, including humans, and some female fertility defects are associated with liver dysfunction (*81–89*). To test whether any aspect of female fertility may contribute to mating-dependent changes in liver immune cell levels, we generated and analyzed a killifish genetic mutant (*TCEB3*) that we serendipitously identified as producing infertile eggs. *TCEB3* (killifish name: *LOC107396610*) encodes a putative transcription elongation factor with ∼40% protein sequence identity and some structural similarity to the human TCEB3 (**fig. S8A**). While expression of the mammalian *TCEB3* gene is not tissue specific (*123, 124*), the killifish *TCEB3* gene is specifically expressed in female gonads, as observed in the Killifish Aging Atlas RNA-seq dataset (**fig. S8B**) (*60*). Males did not express detectable levels of *TCEB3* mRNA in the testis or any of the other 12 tissues assessed (**fig. S8B**). We generated a homozygous *TCEB3* mutant using a CRISPR-based method (*54*) to insert a transcriptional terminator at the beginning of the first exon (**Fig. 5A**; see Methods for details). Both heterozygous (*TCEB3^+/−^*) and homozygous *TCEB3^−/−^*males and females were viable. We verified that compared to their heterozygous siblings (*TCEB3^+/−^*), *TCEB3^−/−^* females had strongly reduced *TCEB3* mRNA expression in oocytes (∼9-fold reduction) (**Fig. 5B**), suggesting a loss-of-function mutant. We observed that *TCEB3^−/−^* females engaged in sexual behavior with males of different genotypes (wildtype and mutant males) and laid eggs in similar number per mating session as *TCEB3^+/−^* females (**Fig. 5C**). However, *TCEB3^−/−^* females laid infertile eggs when mated with either wildtype of mutant males (**Fig. 5C**) (those eggs were also ∼30% smaller than those produced by their heterozygous siblings) (**Fig. 5C** and **fig. S8C**). These results indicate that we have generated a genetic killifish mutant in which females are infertile, probably due to defects in egg maturation.

**Fig. 5.**
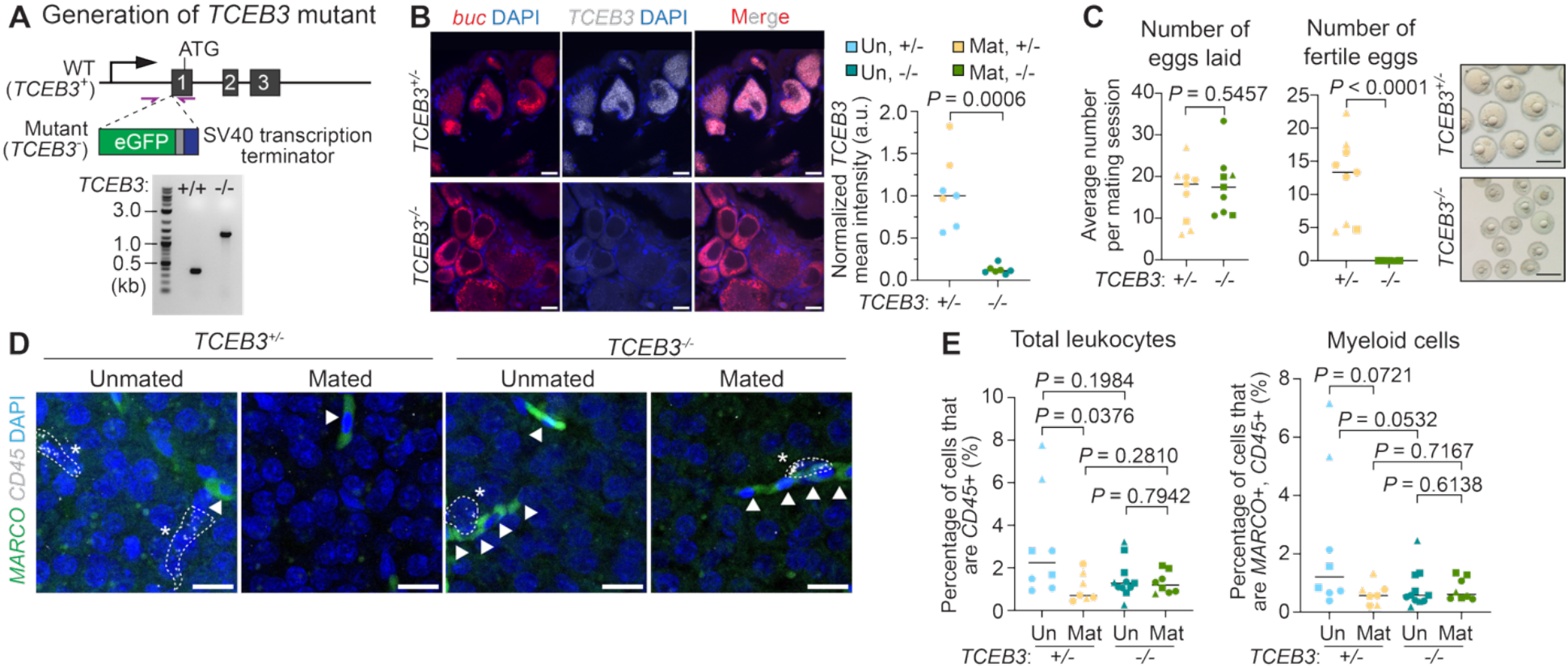
Generation of a mutant that produces infertile eggs to test whether female fertility is involved in the mating effect on young liver immune cells. (A) Schematic of *TCEB3* knockout (top) and DNA gel for genotyping *TCEB3^+/+^* versus *TCEB3^−/−^* fish (bottom). The *eGFP* open reading frame and an SV40 transcription terminator sequence are inserted immediately upstream of the start codon (ATG) of *TCEB3*. Purple arrows indicate the primers used for genotyping (wildtype: 375 bp; inserted: 1.4 kb), and the results were shown in the DNA gel. WT, wildtype. (B) Example RNA *in situ* images (left) and quantification of *TCEB3* mRNA levels in young (2-3 months) *TCEB3^+/−^* and *TCEB3^−/−^* females (right). Ovary sections were stained for an oocyte marker *buc* (red) and *TCEB3* (gray) mRNAs and nuclei by DAPI (blue). Scale bar, 10 µm. Each dot in the graph is a fish (unmated *TCEB3^+/−^*, n = 4 fish; mated *TCEB3^+/−^*, n = 3 fish; unmated *TCEB3^−/−^*, n = 4 fish; mated *TCEB3^−/−^*, n = 3 fish). The normalized *TCEB3* mean intensity is plotted, calculated by first averaging the mean cellular *TCEB3* mRNA staining intensity across all oocytes quantified for each animal, then normalizing to the median of the unmated fish. The bar is the median expression of all fish in a condition. Un, unmated; Mat, mated; a.u., arbitrary unit. The *P* values were determined using the Mann-Whitney *U* test for pairwise comparison. (C) Quantification of the average number of eggs laid per mating session (left), the number of fertile eggs (i.e., eggs that can develop into 1-2 cell stage embryos) per mating session (middle), and example images of the eggs laid 4 hours post mating (right) for young mated *TCEB3^+/−^* (n = 9 fish) and *TCEB3^−/−^* females (n = 9 fish). The bar is the median of all fish in a condition. Symbol shape, cohort number (animals were raised in 3 independent cohorts). Scale bar, 1 mm. The *P* values were determined using the Mann-Whitney *U* test for pairwise comparisons. (D) Example RNA *in situ* images for female liver sections (unmated *TCEB3^+/−^*, n = 8 fish; mated *TCEB3^+/−^*, n = 7 fish; unmated *TCEB3^−/−^*, n = 11 fish; mated *TCEB3^−/−^*, n = 8 fish), staining for *MARCO* (green) and *CD45* (gray) mRNAs and nuclei by DAPI (blue). Dashed lines outline cells. Star, an immune cell (*CD45*+); arrowhead, a red blood cell (autofluorescent). Scale bar, 10 *µm*. (E) Quantification of panel D, plotting the proportion of leukocytes (*CD45+*) and myeloid cells (*MARCO*+, *CD45*+) in the livers of young *TCEB3^+/−^* (unmated *TCEB3^+/−^*, n = 8 fish; mated *TCEB3^+/−^*, n = 7 fish) and *TCEB3^−/−^* (unmated *TCEB3^−/−^*, n = 11 fish; mated *TCEB3^−/−^*, n = 8 fish) females. The bar is the median of all fish in each condition. Symbol shape, cohort number (animals were raised in 3 independent cohorts). The *P* values were determined using the Mann-Whitney *U* test for pairwise comparisons. Note that two-way ANOVA was not performed due to non-normal sample distributions. One mated *TCEB3*^+/−^ fish and one mated *TCEB3*^−/−^ fish were detected as outliers and excluded.

We asked how mating affects liver immune cells in *TCEB3* mutants by comparing heterozygous (*TCEB3^+/−^*) and homozygous (*TCEB3^−/−^*) siblings that were mated with wildtype or mutant males at a young age (2.5-3 months). In the young *TCEB3^+/−^* control females, we found that mating led to significantly lower levels of leukocytes (**Fig. 5D** and **5E**, ‘Total leukocytes,’ *P* = 0.0376), consistent with what we observed for wildtype young females (see **Fig. 2B**), and a trending decrease in myeloid cells (**Fig. 5D** and **5E**, ‘Myeloid cells,’ *P* = 0.0721). In contrast, *TCEB3^−/−^* young females no longer exhibited a mating-dependent reduction in leukocytes (*P* = 0.7942) and myeloid cells (*P* = 0.6138) (**Fig. 5D** and **5E**). Expression of the inflammation marker *IRF9* in non-immune liver cells displayed a similar trend as the immune cells (trending decrease by mating in young *TCEB3^+/−^* control (*P* = 0.1049) but no decrease by mating in *TCEB3^−/−^* mutant (*P* = 0.5027)) (**fig. S8D**). These results suggest that at least in the context of the *TCEB3* mutant, certain aspects of female fertility, perhaps linked to egg maturation, may contribute to the mechanism through which mating reduces liver immune cell abundance at a young age.

### Mating remodels inflammation and metabolism transcriptomic signatures in multiple organs

Are the effects of mating specific to the liver or general across several other organs? We performed RNA-seq on three other organs (brain, gut, and head kidney, which is the killifish primary hematopoietic compartment), comparing mated and unmated killifish males and females at young (2-2.5 months old) and old (4.5 months old) ages (**Fig. 6A** and **fig. S9A**). PCA analysis revealed that, generally, age (often reflected by PC1 and PC2) had a stronger impact on the transcriptomes of these organs than mating status (**Fig. 6A**, **fig. S9B**, and **fig. S9C**). Among these organs, the brain had the greatest number of genes differentially expressed by mating status in both sexes at old age (**Fig. 6B** and **fig. S9D**). Consistently, variance partition analysis revealed a higher degree of mating-age interaction in the brain than in the gut and head kidney when both sexes were analyzed together (**fig. S9E**).

**Fig. 6.**
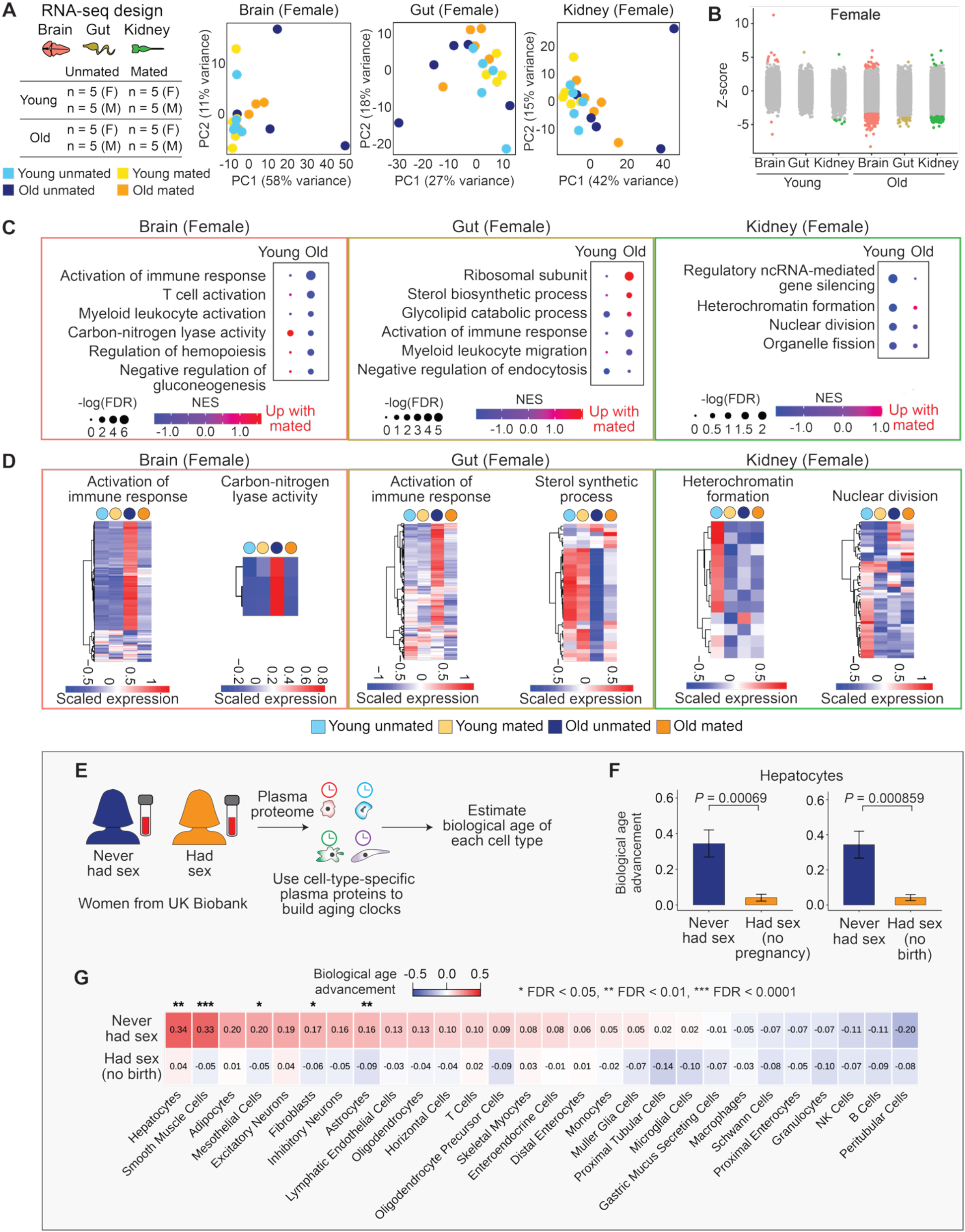
Impact of mating on several killifish organs and association between sexual experience and cell-type age predicted by plasma proteomic aging clocks in UK Biobank women. (A) Organ RNA-seq schematics and PCA for the female brain, gut, and head kidney. Five fish (n = 5, collected from 3 independent cohorts, same as those used in lipidomics) were used for each condition (combination of sex, age, and mating status). Hereinafter, in this figure, young, 2-2.5 months old; old, 4.5 months old. F, female; M, male. In the PCA plot, each dot is an animal (n = 5 fish per condition). (B) Dot plot of the differentially expressed genes by mating at each age in female organs, graphing z-scores. Colored, significantly upregulated or downregulated in mated animals (false discovery rate, FDR ≤ 0.05); gray, not significantly altered (FDR > 0.05). Statistics were performed using a Wald test followed by Benjamini-Hochberg multiple hypothesis correction. (C) Gene set enrichment analysis (GSEA) results for each female organ, identifying the pathways significantly enriched for the genes upregulated or downregulated by mating in young or old animals. NES, normalized enrichment score. Dot size, –log_10_ of the false discovery rate (FDR). (D) Heatmaps of the genes that drive each selected gene ontology (GO) term for each female organ. For each gene, its DESeq2-normalized expression is averaged from the 5 animals for each condition and z-scaled. (E) A framework for estimating cell-type-specific biological aging patterns with plasma proteomics in the UK Biobank. Women were separated into two groups (‘Never had sex’ and ‘Had sex’) based on their responses to the UK Biobank data field question of ‘Age at the first sexual intercourse’ (self-reporting). Cell-type biological age estimates were derived from previously established plasma proteomics aging clock models (*129*). The models quantify biological age based on the abundance of cell-type-specific plasma proteins measurable in blood. Using a population-based approach, age gaps are derived for each cell type: a positive age gap indicates that an individual’s cell type appears biologically older than the same-age peers, whereas a negative age gap indicates a biologically younger aging status. (F) Hepatocyte normalized cell-type biological age gaps (‘biological age advancement’) for women who ‘Never had sex’ (n = 165) versus ‘Had sex’ but had never been pregnant (n = 2,446, ‘no pregnancy’), or versus ‘Had sex’ but had never given birth (n = 3,211, ‘no birth’). Bar height corresponds to the mean of the age gaps within each group, and the error bar corresponds to ± standard error of the mean. Statistics were performed using the Mann-Whitney *U* test. (G) Heatmap of mean normalized cell-type biological age gaps (‘biological age advancement’) for women who ‘Never had sex’ (n = 165) versus ‘Had sex’ but had never given birth (n = 3,211). Rows represent sexual history groups, and columns represent selected cell types. Color indicates mean normalized biological age advancement (value listed in each box of the heatmap), with red indicating a positive biological age advancement and blue indicating a negative biological age advancement. Statistics were performed using the Mann-Whitney *U* test to compare age gap distributions between groups within each cell type, followed by Benjamini-Hochberg multiple hypothesis correction. FDR, false discovery rate.

GSEA revealed that in the brain and gut, immune-related pathways (e.g., ‘activation of immune response,’ ‘myeloid leukocyte migration,’ ‘T cell activation’) were downregulated in mated old females relative to the unmated counterparts (**Fig. 6C** and **6D**). Downregulation of immune-related pathways by mating (e.g., ‘adaptive immune response,’ ‘myeloid leukocyte activation,’ ‘cellular response to chemokine’) was also observed in the brain and gut of old males (**fig. S9F** and **S9G**). In addition, metabolism-related pathways were affected by mating in these two organs of old females (e.g., ‘Sterol biosynthetic process,’ ‘glycolipid catalytic process,’ ‘negative regulation of gluconeogenesis’) (**Fig. 6C** and **6D**), although the specific pathways were distinct from those altered by mating in old female hepatocytes (e.g., ‘olefinic compound metabolic process,’ **Fig. 1F**). Therefore, lifelong mating modulated the transcriptomic signatures of the brain and gut in both sexes, dampening inflammation and altering metabolism-related pathways, particularly at old age.

Not all organs were similarly impacted by mating. In the head kidney and particularly at a young age, mated females showed downregulation of pathways related to cell division (e.g., ‘nuclear division’ and ‘organelle fission’) and gene expression regulation (e.g., ‘gene silencing’ and ‘heterochromatin formation’) compared to the unmated counterparts (**Fig. 6C** and **6D**). In contrast, mated males exhibited downregulation of cytoskeletal-related pathways (e.g., ‘contractile muscle fiber,’ ‘cytoplasmic stress granules,’ ‘multicellular organismal movement’) relative to the unmated counterparts (**fig. S9F** and **S9G**). These observations suggested that mating may affect cell-type balance and architecture in the head kidney, although the functional implications remain unclear. Although these RNA-seq studies lack cell-type resolution, our results suggested that some mating effects could be organ specific.

Collectively, our data supported the idea that, for at least three organs (liver, brain, and gut), lifelong mating remodels organ inflammatory and metabolic signatures in killifish females during aging.

### *Plasma proteomics clocks suggest that sexual experience in women is associated with a* ‘younger biological age’ of several cell types

Our work in the killifish suggested that sexual history influences organ aging, particularly in females. Despite major differences between killifish and human reproductive biology (most notably pregnancy) (*64, 125, 126*), we wondered whether there was any indication that sexual history could associate with differences in biological age in women. Recent developments of plasma proteomic aging clocks provide a non-invasive method for evaluating the biological age of specific organs and cell types (*127–132*). These clocks first classify thousands of blood proteins from healthy individuals by the organs or cell types in which these proteins are most highly expressed and then use these proteins as features to train machine learning models across the human lifespan and predict the ‘biological age’ of new samples in ways that align with clinical outcomes (**Fig. 6E**). For example, individuals with ‘older’ brain cells (astrocytes), as estimated by plasma proteomic aging clocks, also have elevated Alzheimer’s disease risk (*129*).

Using previously established plasma proteomics aging clocks specific to cell types (*129*), we estimated the biological ages of hepatocytes among groups of women with different sexual histories. We performed a cross-sectional study and classified women from the UK Biobank into two groups: those who self-reported an age at first sexual intercourse (‘Had sex’) and those who did not (‘Never had sex’) (see Methods). To limit the influence of pregnancy on women’s biological age, we restricted our analysis in the ‘Had sex’ group to women who met either of the two criteria: had never been pregnant (no pregnancy) or had never given birth (no birth). Among these two categories of women, the ‘Had sex’ group also showed ‘younger’ hepatocytes than the ‘Never had sex’ group (**Fig. 6F**, ‘Hepatocytes,’ *P* = 0.00069 for no pregnancy; *P* = 0.000859 for no birth). Bootstrapping confirmed that the unequal sample sizes between the ‘Never had sex’ and ‘Had sex’ (no pregnancy) groups did not drive these observations (see statistical results in Supplemental Text). Although the directionality of the relationship between sexual experience and biological age cannot be inferred, these results could suggest an association between self-reported sexual experience and a ‘younger biological age’ for hepatocytes in women (in the absence of pregnancy or childbirth).

We examined whether potential confounders, including overall health, specific lifestyle factors, and social connectedness, could explain the difference in predicted ages between women with different sexual histories. We found that the ‘younger hepatocyte age’ observed in the ‘Had sex’ group (no birth) did not necessarily associate with better overall health in this group, as the two groups showed broadly similar diagnosis frequency of major diseases, despite modest differences in age, BMI, hypertension, and thyroid disease (**fig. S10A**). The ‘Had sex’ group also did not have lower smoking and alcohol use compared to the ‘Never had sex’ group (**fig. S10B** and **S10C**). A critical difference between the two groups was that the ‘Had sex’ group reported greater ability to confide in others, a proxy for lower loneliness and stronger social connection (**fig. S10D**). However, when we matched the groups for ‘loneliness’ level, the ‘Had sex’ group (no birth) still showed a younger predicted hepatocyte age than the ‘Never had sex’ group (**fig. S10E**). These findings suggested that the association between sexual experience and younger hepatocytes in women is unlikely to be solely explained by differences in overall health, smoking and alcohol use, and social connectedness, although other confounders, such as the use of contraceptives, medications, and socioeconomic factors, could contribute.

We further extended this analysis using plasma proteomics clocks related to 28 cell types that constitute the human brain, gastrointestinal organs, and kidney (corresponding to the organs we had tested in killifish). Several cell types were estimated to be significantly ‘younger’ by plasma proteomics clocks in women from the ‘Had sex’ group (no birth) compared to women from the ‘Never had sex’ group (**Fig. 6G**). These included hepatocytes, smooth muscle cells, mesothelial cells, fibroblasts, and astrocytes (**Fig. 6G**). Several other cell types (e.g., proximal tubular cells and gastric mucus-secreting cells) did not exhibit significantly younger or older estimated age in the ‘Had sex’ group (**Fig. 6G**).

Therefore, plasma proteomic aging clocks suggested a potential associated between self-reported sexual experience in women and a younger biological age specific to some, but not all, cell types, although the directionality of the relationship between sexual experience and biological age cannot be inferred.

## Discussion

In this study, we have shown that lifelong mating in African turquoise killifish affects organ aging in ways specific to biological sex and cell type. Mating ‘rejuvenates’ the transcriptomic profiles of specific liver cells and dampens inflammatory signatures in females (but not in males) at old age. Functionally, mating reduces hepatic accumulation of specific lipid classes and is associated with improved detoxification resilience in old females, suggesting a beneficial effect. More generally, mating induces transcriptomic programs characteristic of reduced inflammation and metabolic rewiring in other female organs, including the brain and gut. Finally, analysis of publicly available human plasma proteomics data provides evidence that, in women, based on self-reporting, sexual experience is associated with a ‘younger’ biological age in several cell types.

Our study suggests that lifelong mating can have a beneficial effect in females, at least under one drug treatment condition. This is surprising given that reproduction is often viewed as having a cost to health, particularly in females. The ‘disposable soma’ theory (*133*) suggests an evolutionary trade-off between reproduction and longevity. For example, in invertebrates, continuous mating reduces the lifespan of male and female *Drosophila melanogaster* (*12, 20, 21, 23–25, 134–136*) and *C. elegans* hermaphrodites (*9–11, 18, 19*). Mating also shortens the lifespan of female mice (independent of pregnancy) (*39*). Given that lifespan is an integrative systemic outcome, it is possible that mating has both beneficial and detrimental organ-specific effects. Future studies that systematically examine additional organs and tissues in the context of mating could provide a more complete assessment of the mating effects on health.

In humans, the relationship between sexual history and women’s health is complex. Early childbirth is associated with a higher likelihood of frailty and more advanced epigenetic age in women (*137*), and the number of children is negatively correlated with maternal longevity in several, though not all, populations (*138–142*). In contrast, a recent study reported that moderate sexual frequency was associated with lower all-cause mortality among young and middle-aged women (*44*). Our work with plasma proteomics aging clocks suggests that sexual experience in women (when not resulting in pregnancy or childbirth) was associated with a ‘younger’ biological age of several cell types, including hepatocytes. This feature was also observed in killifish females, despite major differences between killifish and human reproductive biology (*64, 125, 126*).

The effect of sexual reproduction on organ aging differed in males and females. For example, sexual reproduction dampened the age-dependent increases in immune cells and inflammatory signatures in the livers of females but not males. Nevertheless, sexual reproduction also affected several age-dependent transcriptomic signatures in the brain and the gut in males, as well as the liver lipidome. Sex-dimorphic responses have been observed for aging and longevity interventions in several species (*60, 64–67, 143–157*). Part of the sex-dimorphic effect of mating on liver health could be due to the tight connection between the liver and female fertility in vertebrates (*158–166*). In support of this idea, we observed that a killifish genetic mutant exhibiting female sterility (likely due to defects in egg maturation) reduced the effect of mating on liver immune cells in young females.

Lastly, sexual reproduction is a complex process that involves gamete production, social interaction, physical contact, gamete release, and fertilization. In *C. elegans* and *Drosophila*, pheromones, seminal fluid, and sexual reception contribute to mating-induced effects (*9–11, 18, 19, 134–136, 167*). In killifish, the use of the *TCEB3* mutant, which sexually interacts but exhibits female-specific sterility, provides a first indication that social interaction and physical contact are unlikely to be the main drivers of at least one mating-induced effect (the dampening of immune cell levels in the liver). Future iterations of our model could incorporate female-female interactions to isolate social stimuli from sexual interaction.

Our study provides evidence that sexual history matters for vertebrate aging, even though mating status is often overlooked in laboratory settings. Given how common sexual reproduction is in nature and society, considering ‘mating status’ as a biological variable would be important for studies of various biological questions, such as adult physiology, disease modeling, and therapeutic development.

## Acknowledgements

We thank all members of the Brunet Lab, particularly Felix Boos, Giulia Notarangelo, Jeeyoon Na, Angela Pogson, Rahul Nagvekar, Gina Wade, and Nimrod Rappoport for scientific discussion and feedback on the manuscript. We thank Felix Boos, Gina Wade, and Nimrod Rappoport for help with independent code validation. We also thank Natalie Schmahl, Sarah Boyle, Jacob Chung, and Rogelio Barajas for their assistance with killifish husbandry over the years.

## Funding

This work was supported by the Milky Way Research Foundation (A.B.), Glenn Foundation for Medical Research (A.B.), CZ Biohub Investigator Program (A.B.), NOMIS Foundation (A.B., T.W.-C.), Simons Foundation (A.B., T.W.-C.), Phil and Penny Knight Initiative for Brain Resilience (T.W.-C.), and Michael J. Fox Foundation (T.W.-C). D.Y. was supported by the National Institutes of Health (NIH)/NIA F99/K00AG083292. E.K.C. was supported by the NIH Training Grant (T32MH020016) awarded to the Leland Stanford Junior University and the Fondation Bertarelli Graduate Fellowship Fund. J.C. is a Jane Coffin Childs fellow, Stanford Jump Start Awardee, and Stanford Katharine McCormick fellow.

## Author contributions

J.C. and A.B. conceptualized the study. J.C. designed, performed, and analyzed all experiments and performed all computational analyses, except those indicated below. A.T. helped to harvest all tissues for single-cell RNA-seq and multi-organ RNA-seq, to perform single-cell dissociation and library preparation, and to perform histological sectioning, HCR staining, imaging, and analysis. E.K.C. helped to harvest tissues for multi-organ RNA-seq and to perform RNA extraction and prepare sequencing library, as well as performed read mapping of the multi-organ RNA-seq data under the supervision of T.W-C.. D.Y. optimized and ran the CellBiAge clocks under the supervision of A.E.W.. X.L. established and optimized the lipidomics pipeline, performed mass spectrometry and data acquisition, and preprocessed the data under the supervision of M.P.S.. K.L.C and V.A.B performed human association studies using UK Biobank data under the supervision of T.W.-C.. M.H. helped to optimize the lipid extraction pipeline, to perform lipid extraction, and to set up killifish cohorts. L.P. assisted with histological sectioning, HCR staining, imaging, and analysis. S.V.P. helped with acetaminophen sensitivity assays and setting up killifish cohorts. R.C.K. helped to optimize the controlled mating experiments and to set up killifish cohorts. J.C. and A.B. wrote the manuscript, and other authors provided comments.

## Competing interests

T.W-C. is a co-founder and scientific advisor of Teal Omics Inc., Alkahest Inc., and Qinotto Inc. and has received equity stakes in these companies. M.P.S. is a cofounder and scientific advisor of Crosshair Therapeutics, Exposomics, Filtricine, Fodsel, Iollo, InVu Health, January AI, Marble Therapeutics, Mirvie, Next Thought AI, Orange Street Ventures, Personalis, Protos Biologics, Qbio, RTHM, and SensOmics, and is a scientific advisor to Abbratech, Applied Cognition, Enovone, Jupiter Therapeutics, M3 Helium, Mitrix, Neuvivo, Onza, Sigil Biosciences, TranscribeGlass, WndrHLTH, and Yuvan Research. A.B. is a member of the scientific advisory board of Calico. All other authors have certified that they have no competing interests to declare.

## Data, code, and material availability

For publication, all the RNA-seq and scRNA-seq data (raw FASTQ files) will be deposited to the Sequence Read Archive (SRA). The normalized expression data matrix of RNA-seq will be deposited to NCBI-GEO and the scRNA-seq Seurat objects will be deposited to Zenodo. The lipidomics data will be deposited to the NIH Common Fund’s National Metabolomics Data Repository (NMDR) website and the Metabolomics Workbench. All genetic mutants and reagents generated in this paper will be made available without restrictions. All original code will be made available in a public GitHub repository upon publication.

## Materials and Methods

### African turquoise killifish husbandry

All experiments used the GRZ strain of the African turquoise killifish species *N. furzeri*. Fish were housed in a 26°C circulating water system, with a conductivity ranging from 1200 to 4500 μS/cm, a pH of 6.8 to 7.2, and a daily exchange of 10% with water treated by reverse osmosis. A 12-hour day/night cycle was used. The fish were housed in the Stanford Research Animal Facility under protocols approved by the Stanford Administrative Panel on Laboratory Animal Care (IACUC protocol # 13645).

All fish were raised from embryos collected from group breeding tanks (one male paired with at least three females in a 9.8 L tank) or individual breeding tanks (one male paired with one female in a 2.8 L tank). Breeders were bred with sand trays in the tanks for embryo collection. After ∼4 hours, the sand trays were collected, and embryos were separated from the sand by gently sieving (*168*). To reduce contamination, the embryos were rinsed twice in ∼4 mL embryo solution (Ringer’s solution [Sigma-Aldrich, 96724] supplemented with 0.01% methylene blue [Kordon, #37344]). Washed embryos were placed in fresh embryo solution in 30 mm x 15 mm Petri dishes (Greiner Bio-One, 627102) at a density of 10 to 30 embryos per plate and incubated at 28°C for ∼2 weeks. Next, the embryos were placed on moist coconut fiber at 27°C. After ∼2 weeks on coco fiber, fish were hatched in 10 mL chilled (4°C) 1 g/L humic acid solution (Sigma-Aldrich, 53680) and incubated at room temperature overnight. The hatched fish were transferred to the animal facility and placed in 0.8 L tanks at a density of 10-20 fish per tank on the circulating water system with gentle water flow (∼1-2 drips per second) for one week. Fish were split to a density of 4 fish per 0.8 L tank in the second week after hatching, two fish per 0.8 L tank in the third week, and one fish per 0.8 L tank in the fourth week. Juvenile fish were fed live brine shrimp daily from hatching until sexual maturity. Five weeks after hatching, each fish was photographed (top-down, with fish placed in ∼2 cm deep water), transferred to a 2.8 L tank, and sexed by the tail fin color (males and not females exhibit vivid colors). Fish with severe gill defects, curved spines, and an inability to float (‘belly sliders’) were excluded. Once the fish were housed in 2.8 tanks, they were fed using a custom-made feeder twice a day, 15 mg per feeding, for 30 mg of Otohime fish pellet (Reed Mariculture, Otohime C1) daily.

### Mating cohorts

Fish were randomly assigned to the unmated or mated groups using the ‘rand()’ function in Excel. Each mating pair included one male and one female hatched in the same cohort. The same mating pair was maintained throughout life until one fish died, and then the dead partner was replaced by another fish of similar age (hatch dates were within 1 month of each other). All tank positions were randomly assigned to reduce experimental confounds, such as the tank location on the water system. Male and female tanks, as well as the ‘mated’ and ‘unmated’ tanks, alternated (tank order: mated female, mated male, unmated female, unmated male, etc.).

The mating experiments began at sexual maturity, which corresponded to 5-6 weeks post-hatching (unless otherwise specified). Twice a week (e.g., Mondays and Thursdays), each fish was fed 15 mg of Otohime fish pellet at 8:30 - 9 AM. After the fish finished all the food pellets (∼10-20 min), a white weigh boat containing ∼25 mL autoclaved fine-grain sand (Cemex 30 Mesh Sand, 200000278) (‘sand tray’) was placed in the female tank. The male partner was transferred into the female tank with a soft fish net. To control for fish handling and environmental change, we also placed a sand tray in the tank of each unmated fish. Unmated males also received a ‘mock cross,’ in which we used a soft fish net to pick up the males and place them back in their own tanks to mimic fish transfer. After 4 hours of mating or ‘mock cross,’ the sand trays were removed from the tanks, and the male partners were transferred back to their own tanks (‘uncross’). The unmated males also received a ‘mock cross’ to mimic the ‘uncross’ experience of the mated males. Lastly, each fish was fed 15 mg of fish pellet. We note that most mating events occurred in the first 2 hours of each mating session. Prolonged co-housing between male and female partners in a 2.8 L tank (over 8 hours) can sometimes lead to ‘harassment’ of the female, causing her to hide at the bottom of the tank away from the male. Because competition for food could occur between males and females, we separated mated males and females after each mating session to better control for these behavioral and dietary confounds.

After each 4-hour mating session, the sand trays were collected, and embryos were separated from the sand by gentle sieving (*168*) and rinsed twice with ∼4 mL of embryo solution. The number of eggs laid by unmated females was recorded by manual counting during collection. For mated females, all the embryos/eggs were collected and imaged using an Olympus stereomicroscope with an eyepiece mount-on AmScope AU1000 camera (AmScope, AU1000), using default settings. Unfertilized eggs were removed, and the remaining embryos were cleaned with embryo solution and incubated in ∼4 mL embryo at 28°C. Dead embryos were removed daily.

### Egg quantification

All egg images for both wildtype and *TCEB3* mutants were analyzed in FIJI (*169*) and manually counted. The total number of eggs laid by mated and unmated females was measured on the collection day. The mean number of eggs laid per mating session was averaged from 3-6 mating sessions at 6-8 weeks after hatching for young females and at 16-18 weeks after hatching for old females. Fertile eggs corresponded to eggs with at least 1-2 cells present at >3 h post collection. The number of eggs released per mating session (regardless of whether they were fertile) was plotted.

For egg size quantification, we drew a straight line along the longest axis of each egg (‘diameter in pixel’), as well as a line marking a 1-mm distance on the ruler (‘ruler pixel per mm’). We calculated the diameter in millimeters as the ratio of ‘diameter in pixels’ over ‘ruler pixel per mm.’ For each animal, we quantified at least 30 eggs at 7-9 weeks of age, across at least 2 mating sessions. All results are included in Extended Data Table 1.

### *TCEB3* mutant generation and genotyping

We became interested in the *TCEB3* gene while generating the Killifish Aging Atlas dataset (*60*) because of its unique expression pattern in female gonads in killifish. To understand the function of *TCEB3*, we generated a knockout mutant and observed that its females laid infertile eggs. Thus, we used this mutant in our study to disrupt female fertility. The *TCEB3* transcription-null allele was generated using a CRISPR-based knock-in system (*54*). The gene-specific guide RNA was designed using the CHOPCHOP program (https://chopchop.cbu.uib.no/) and the ‘Nfu_20140520/Jena’ *Nothobranchius furzeri* genome. This guide RNA (5’-G<u>ATG</u>GCCAGAAAATCCGATG-3’), located at the *TCEB3/LOC107396610* gene translation start site (ATG was underlined), was synthesized as a custom 2-nmol Alt-R^TM^ CRISPR RNA (crRNA) sequence by IDT (Newark, NJ, USA). The DNA repair template was acquired as a gBlock from IDT (Newark, NJ, USA). It included four features in the following 5’-to-3’ order: 1) ∼100 bp homolog sequences immediately upstream of the start codon of the *TCEB3/LOC107396610* open reading frame, 2) the *eGFP* open reading frame, 3) the *SV40* transcription terminator sequence, and 4) ∼100 bp homolog sequences immediately downstream of the *TCEB3* start codon. The PAM sequence was mutated from TGG to TTG to prevent re-cutting of the target site. Extended Data Table 2 includes the sequences of the guide RNA, repair templates, and genotyping primers.

For microinjection, we first reconstituted the reagents to 100 µM crRNA, 100 µM tracrRNA (Alt-R^TM^ CRISPR-Cas9 tracrRNA, IDT, 1072533), and repair template (∼150 ng/µL) using 20 µL, 200 µL, and 7 µL nuclease-free duplex buffer (IDT, 11-01-03-01), respectively. Next, we incubated the gRNA mix (1 µL 100 µM tracrRNA, 1 µL 100 µM crRNA, and 31.3 µL nuclease-free duplex buffer) at 95°C for 5 min and then cooled to room temperature. A RNP complex mix (10 µL of the gRNA mix, 0.5 µL Alt-R^TM^ S.p. Cas9 nuclease V3 protein [10 µg/µL, IDT, 1081058], 5.5 µL PBS [Corning, 21-040-CV]) was assembled by incubating at 37°C for 10 min and then placed on ice. Lastly, we generated the injection solution by mixing 1 µL of the repair template, 1 µL of the Alt-R^TM^ HDR enhancer protein (IDT, 10029790), 8 µL of the RNP complex mix, and 1 µL diluted phenol red in PBS (1:3 dilution of commercial phenol red can prevent needle clogging).

Embryos were collected from group breeding tanks (see the ‘African turquoise killifish husbandry’ section), thoroughly rinsed with embryo solution, and injected at 1-cell and 2-cell stages on an injection pad made from 1% agarose (*56*). Embryos were incubated, monitored, hatched, and raised as described in the ‘African turquoise killifish husbandry’ section. To extract genomic DNA for genotyping, a small piece of tail tissue from hatched juvenile fish (3-5 weeks old) was obtained and incubated in 50 µL DNA extraction buffer (50 µL DirectPCR Lysis Reagent, tail [Fisher Scientific, NC9724951] and 1 µL proteinase K [Invitrogen, 25530049]) at 55°C for 2 hours followed by enzyme heat inactivation at 100°C for 10 min. PCR reaction was set up using GoTaq Green 2x Master Mix (Promega, M7123), the crude DNA lysate, and appropriate primers (see Extended Data Table 2 for primer information). PCR products were confirmed by gel electrophoresis for size and by sequence identity using column purification followed by Sanger sequencing (service provided by MCLab, South San Francisco, CA, USA).

To generate *TCEB3^−/−^* mutants, mosaic F0 fish were backcrossed to wildtype fish, and F1 fish from a single F0 fish were used for further backcrossing to wildtype fish for 5 generations. Both *TCEB3^+/−^*and *TCEB3^−/−^* males and females were viable and had no overt differences in physical appearance, size, or sexual maturation rate compared to wildtype animals. For experiments, *TCEB3^+/−^* females in group breeding tanks were crossed to *TCEB3^−/−^* males (these males are fertile), generating ∼50% *TCEB3^+/−^* (heterozygous) and ∼50% *TCEB3^−/−^*(homozygous) offspring with no sex bias. For each mating cohort, female offspring siblings were randomly paired with male siblings of either genotype because the male genotype did not appear to affect mating performance. The mating experiments associated with the *TCEB3* mutants were performed as described in the ‘Mating cohorts’ section.

### Sample collection for liver single-cell RNA-seq

All procedures were performed in an RNase-free environment, and all tools and procedural surfaces were cleaned using RNaseZap (Thermo Fisher, AM9784). All reagents were nuclease-free. Fish were randomly assigned a harvest order using the ‘rand()’ function in Excel. If a fish with the highest harvest number died, the second-highest harvest number would be used. Fish were harvested at 2 months and 4-4.5 months of age. Mating pairs that failed to produce at least an average number of 2 fertilized eggs per mating session over the lifetime were excluded from analysis because these mating pairs might not have sexually interacted. Males who severely injured or killed their female partners due to excessive aggression were also excluded from the analysis. Lastly, we excluded fish that showed overt signs of illness on the day of sample collection, including not eating, sinking to the bottom of the tank, and being unresponsive to tapping on the tank wall.

Sample collection occurred 24-27 hours after the last mating. On that day, each fish was fed 15 mg of Otohime fish pellet from 8:30 - 8:45 AM and typically sacrificed from 11:30 AM to 1:30 PM (3-5 hours after the last meal). The dissection orders were balanced by mating status (for example, if mated fish were dissected before unmated fish on harvest day 1, then the dissection order would be reversed on harvest day 2).

At ∼11:30 AM on a harvest day, we transferred the fish (typically 4 fish) from the animal facility to the lab space. To limit batch effect, two operators (J.C., A.T.) dissected the mated and unmated fish of the same sex simultaneously, and each operator dissected half of the animals of a single condition. See Extended Data Table 3 for harvest details. Male fish were dissected before female fish. Fish were sacrificed by incubating them in system water supplemented with ice (0°C) for 2-3 min for rapid cooling. The fish were dried and placed on an ice-cold Sylgard-coated Petri dish filled with wet ice and covered in plastic wrap. An incision was made from the vent (anus), along the midline of the abdominal region, to just before the heart. The abdominal muscles along the left flank were removed to expose the liver, which was carefully removed from the body cavity using tweezers. After removing the gallbladder and residual fat tissue on the livers, the livers were stored in ice-cold PBS (Corning 21-040-CV) until digestion. Dissecting all four fish typically took ∼30 minutes (∼15 minutes per dissection batch). The livers were cut in half along the anterior-posterior axis while positioned concave up. The left side of the liver (still concave up) was used for single-cell dissociation, and the right side of the liver was fixed in 4% paraformaldehyde (PFA) in PBS (Santa Cruz, CAS 20525-89-4) at 4°C for ∼20-24 hours. We note that our single-cell dissociation protocol would yield sufficient live cells (3×10^5^ – 2×10^6^) from half of a liver. The fixed half of the liver was used for validation staining. Because each half-liver came from the same animal as a given scRNA-seq sample, this setup should allow us to detect any dissociation-induced artifact if the expression of a gene from scRNA-seq did not match the *in situ* staining. Alternatively, there could be spatial differences between the left and the right sides of the liver. We thus stained additional cohorts during validation to test whether our main findings hold for the whole liver.

We used a cold-active digestive enzyme from *Bacillus licheniformis* (Sigma-Aldrich, P5380) to generate single-cell suspensions from the liver. This enzyme is used for digesting sensitive tissues (e.g., kidney in mice (*170, 171*), lung and nasal epithelium in humans (*172, 173*) and provides the advantage of performing digestion on ice, greatly reducing transcriptional changes and RNA degradation during digestion (*170, 171*). For female livers, we fully optimized the digestion enzyme concentration, digestion duration, centrifugation speed and timing, and the reagents (e.g., specific brands of tubes) for the dissociation protocol described below. We note that male liver dissociation led to some cell loss due to clumping. For each liver, we placed a 1.5 mL Protein LoBind tube (Eppendorf, 022431081) containing 250 µL PBS on a metal block on ice. Next, we opened the tube so that the cap was in physical contact with the ice-cold metal block to keep it cold. The half liver was transferred to the cap, which served as a tray for tissue cutting. We quickly chopped the half-liver 100 times with small scissors to produce a fine tissue paste. The tube was capped and inverted several times to dislodge the tissue paste into PBS. Next, the tissue was vigorously mixed 30-50 times using a p1000 pipette set to 150 µL, pipetting up and down ∼1-2 times per second. We verified that no visible clumps were present after the mixing. Next, we added 250 µL 2x digestion mixture (25 µL 10 mg/mL *Bacillus licheniformis* enzyme dissolved in PBS, 1 µL 0.5 M EDTA pH8.0 [Ambion, AM9261], and 224 µL PBS) and mixed ∼20 times using a p1000 pipette set at 250 µL, gently pipetting up and down approximately once every 2 seconds. If visible clumps were present, they were transferred to the tube cap and mixed vigorously. The digestion tube was incubated on ice for ∼10 min and gently mixed 10 times every 2 min. After the 10 min incubation, digestion was monitored under a brightfield microscope and stopped once ∼70% of cells were individualized, or at most ∼50% of cells showed mild plasma membrane deformation (which could be an early sign of ‘blebbing’). Typically, 2-month-old liver samples took ∼12-18 min to complete digestion, and 4-month-old liver samples took ∼15-27 min. To preserve sample quality, the following steps were performed in a 4°C room. Each sample was filtered through a pre-wet 40 µm strainer (Corning, 431750) on ice, and the flow-through was collected in a 50-mL polypropylene conical tube (Falcon, 352070). The strainer was gently rinsed with 10.5 mL of cold 0.1% BSA/PBS (diluted from 7.5% Gibco BSA Fraction V [Gibco, 15260037]). All the flow-through was gently stirred 2-3 times to mix and then decanted into a 15-mL polypropylene conical tube (Fisher Scientific, 352196). The deep V-shape of the 15-mL conical tube helps pellet the cells during centrifugation. The samples were centrifuged in a swinging-bucket rotor at 200 x g at 4°C for 3 min. The supernatant was removed until ∼50 µL liquid remained, then 75 µL 2.5% BSA/PBS (diluted from 7.5% BSA) was added, and the cell pellet was resuspended by gently pipetting using a p200 pipette ∼30 times and transferred to a 1.5 mL low-adhesion tube (USA Scientific, 14152600). The single-cell suspension was stored on ice during cell counting.

To determine the cell density of the single-cell suspension, we mixed 2 µL of each single-cell suspension with 18 µL vital dye (1 µg/mL propidium iodide [BioLegend, 421301], 5 µM Hoechst [Cayman Chemical Company, 16756], 2.66 µg/mL acridine orange [Abcam, ab270791]) at room temperature, loaded the diluted cells onto a disposable INCYTO C-Chip for cell counting (Fisher Scientific, DHC-F015), and imaged on a fluorescence microscope using a 20x objective and a ZEN blue software (3.0). The number of live cells (propidium iodine-negative and acridine orange-positive) was counted in three small squares of each C-chip. All liver samples contained >88% live cells and had at least 3×10^5^ cells (see Extended Data Table 3 for metadata).

We used a single-cell transcriptome pipeline from Parse Biosciences, which enabled us to fix and process all 32 samples (collected on different days) in parallel. Before fixation, we first removed red blood cell to reduce their number in our dataset. We found that commercial red blood cell lysis protocols could increase cell death. Therefore, we took advantage of the differential buoyancy of cell types and performed sequential centrifugation as follows. Each single-cell suspension was centrifuged at 200 x g, 4°C, for 3 min at a tabletop centrifuge, 75 µL of the supernatant was removed without disturbing the cell pellet (reducing the supernatant volume helps ‘collapse’ the red blood cells onto the main cell pellet), and centrifuged again at 200 x g, 4°C, for 3 min. After the second centrifugation step, the red blood cells (which are red) would be located at the top edges of the cell pellet, leaving a red outline. We carefully removed the red outline using a p10 pipette (typically < 3-5 µL) while limiting the removal of other non-red cell types. Next, we used the Parse Evercode Cell Fixation kit v2 (Parse, ECF2101) and followed the protocol V2.1.2, except that half of the volumes were used for all the reactions. Briefly, the cell pellet was resuspended in 375 µL Cell Prefixation buffer supplemented with 0.469% BSA (100 µL 7.5% Gibco BSA Fraction V [Gibco, 15260037] in 1500 µL Cell Prefixation buffer) on ice and then filtered through a 40 µm strainer into a 5 mL DNA LoBind tube (Eppendorf, 0030122208; ideally should be a 5 mL Protein LoBind tube Eppendorf, 0030108302). Next, the cell mixture was incubated on ice in the following solution provided by the Parse kit: 125 µL Cell Fixation Solution for 10 min, 40 µL Cell Permeabilization Solution for 3 min, and 500 µL Cell Neutralization Buffer. The cell mixture was centrifuged in a swinging-bucket rotor at 200 x g at 4°C for 10 min. The supernatant was removed, and the cell pellet was resuspended in 100 µL cold Cell Buffer and filtered through a 40 µm strainer into a 1.5 mL Protein LoBind tube. Three rounds of ∼1.67 µL DMSO (adjusted based on the average final volume of the four samples in each processing batch) were added to the cell mixture, with a 1-minute incubation between additions, for a final DMSO concentration of 5% in the cell mixture. A 10 µL aliquot was prepared for each sample (for later cell counting), and the aliquot and the sample tubes were placed in a Mr. Frosty (Thermo Scientific, 5100-0001) filled with isopropanol and stored at −80°C (controlled cooling preserves cells better).

### Liver single-cell RNA-seq library preparation

#### cDNA library generation and sequencing

We used a multiplex single-cell RNA-seq kit (Parse Biosciences, Evercode WT v3) to generate cDNA libraries. In this single-cell RNA-seq pipeline, each cell acts as a ‘reaction compartment,’ where the RNA molecules in a given cell are reverse transcribed and uniquely barcoded through rounds of split-pool combinational barcoding. This pipeline can profile up to 100,000 cells across 48 biological samples. We followed the Evercode WT v3 protocol precisely, which is briefly described below. We first determined the cell density of each fixed sample using the 10 µL aliquots for cell counting. The 10 µL aliquots for these samples were thawed at a 37°C water bath (took ∼30 sec), mixed with 10 µL vital dye, and counted using an INCYTO C-Chip as described above in ‘Sample collection for liver single-cell RNA-seq.’ We input the cell counts into the ‘Sample Loading Table’ from Parse Biosciences to calculate the dilution volume of each sample, targeting to load 200,000 cells per sample. Next, we prepared the cell dilution plate. For each sample, the appropriate volume of dilution buffer was loaded into a V-bottom 96-well plate (Falcon, 353263), termed the ‘cell dilution plate,’ one day before sample loading (preloading the buffer accelerated sample processing) and then frozen at −20°C overnight. We used a polypropylene V-bottom 96-well plate (and not a flat-bottom plate) to reduce cell loss. The next day, the samples were thawed in a 37°C water bath (∼45 sec for ∼100 µL cells) in a batch of 6-8 samples and then loaded onto the cell dilution plate on ice. All samples were diluted within 40 min after thawing the first sample.

Next, we performed *in situ* cell barcoding. After gentle mixing, the cells were transferred to a barcoding Round 1 plate and incubated on a deep-well thermocycler (Bio-Rad, 1851197) at 50°C for 10 min, three cycles of serial temperature gradient (8°C for 12 sec, 15°C for 45 sec, 20°C for 45 sec, 30°C for 30 sec, 42°C for 2 min, and 50°C for 3 min), and 50°C for 5 min. All samples were pooled into a 5 mL Protein LoBind tube (Eppendorf, 0030108302), 9.6 µL Spin Additive was added, and the tube was centrifuged in a swinging-bucket rotor at 200 x g at 4°C for 10 min. The supernatant was removed, and the cell pellet was resuspended in 1 mL Resuspension Buffer, followed by another 1 mL Resuspension Buffer for a total of 2 mL Buffer. Next, the 2 mL sample in the Resuspension Buffer was gently mixed with 1.95 mL Round 2 Ligation Buffer and 20 µL Round 2 Ligation enzyme. The cell mixture was poured into a basin (Corning, 4870), and 40 µL of the cell mixture was added to each well of the Round 2 Ligation Plate (96-well) to incubate at 16°C for 15 min. Ligation was stopped by adding 10 µL Round 2 Stop Buffer and incubating at 16°C for 5 min. The cells from each well were pooled, filtered through a 40 µm strainer into a basin (Corning, 4870), gently mixed with 20 µL Round 3 Ligation Enzyme, and transferred into the Round 3 Plate at 50 µL per well. The Round 3 Plate was incubated at 16°C for 15 min. The ligation reaction was stopped by adding 20 µL Round 3 Stop Buffer. The cells were pooled, filtered through a 40 µm strainer into a 15-mL polypropylene conical tube (Fisher Scientific, 352196), 70 µL Spin Additive was added, and the mixture was centrifuged in a swinging-bucket rotor at 200 x g at 4°C for 10 min. After the supernatant was removed, the cell pellet was gently resuspended in 1 mL of Pre-Lysis Wash Buffer, and then an additional 3 mL of Pre-Lysis Wash Buffer was added. The cells were centrifuged as before, the supernatant was removed, and the cells were resuspended in 60 µL Pre-Lysis Dilution Buffer. Cell density (∼1500 cells/µL) was determined using an INCYTO C-Chip described above after mixing 5 µL of cells and 15 µL of vital dye. A total of 125,000 cells were used, split across 8 sub-libraries. Using the ‘Sublibrary Loading’ guideline, we mixed 8.93 µL cells, 16.07 µL Pre-Lysis Dilution buffer, and 30 µL Lysis Master Mix. The cells were lysed at 65°C for 15 min and then frozen at −80°C.

To perform cDNA capture and amplification, the cDNAs were incubated with 2.5 µL Capture Enhancer at room temperature for 10 min and then captured by incubating 50 µL Streptavidin Beads (pre-equilibrated in Binding Buffer) on a vortexer (∼800-1000 RPM) at room temperature for 30 min. The beads were washed twice with 125 µL Wash Buffer 1, once with 125 µL Wash Buffer 2, and once with 125 µL Wash Buffer 3. On-bead template switching was performed by incubating the beads with Template Switch Master Mix at room temperature for 30 min, then at 42°C for 1 hour. The beads were washed once with Wash Buffer 2, once with Wash Buffer 3, and then incubated with 100 µL Amplification Master Mix to amplify the cDNAs for 5 cycles at 65°C (annealing temperature) and then 8 cycles at 67°C (annealing temperature). The amplified cDNA was purified using 0.8x AMPure beads (80 µL per 100 µL reaction; Beckman Coulter, A63881) with two 180 µL 85% ethanol washes and eluted in 25 µL nuclease-free water. The concentration and quality of each cDNA sub-library were measured using a Quant-iT dsDNA HS Kit (Thermo Fisher, 33120), as well as the Agilent 2100 Bioanalyzer and Agilent’s High Sensitivity DNA Kit (Agilent, #5067-4626) by the Stanford Protein and Nucleic Acid Facility.

To prepare the sequencing library, we first fragmented the cDNA sub-libraries in a 50 µL reaction by incubating 200 ng of each cDNA sub-library with 15 µL Fragmentation and End Prep Master Mix at 32°C for 10 min, then 65°C for 30 min. The fragmented cDNAs were size-selected and purified using 30 µL AMPure beads per library, then 10 µL AMPure beads per library, washed twice with 180 µL 85% ethanol, and eluted in 50 µL nuclease-free water. Adaptors (with Illumina TruSeq Read 1 sequence) were ligated to the 5’ end of each fragmented cDNA library via incubation with 50 µL Adapter Ligation Master Mix (in a 100 µL reaction) at 20°C for 15 min. Adapter-ligated DNA was purified using 0.8x AMPure beads as described above and eluted in 21 µL nuclease-free water. The purified adapter-ligated DNA (21 µL per sub-library) was amplified for 10 cycles with 4 µL of the unique indexing primer pairs (Illumina TruSeq Read 1 and R2 primers provided in the Parse UDI Plate) and 25 µL Library Amp Mix. The sequencing libraries were size-selected and purified using 30 µL AMPure beads per library, then 10 µL AMPure beads per library, washed twice with 180 µL 85% ethanol, and eluted in 20 µL nuclease-free water. The concentration and quality of each sequencing library were measured using a Quant-iT dsDNA HS Kit (Thermo Fisher, 33120), as well as the Agilent 2100 Bioanalyzer and Agilent’s High Sensitivity DNA Kit (Agilent, #5067-4626) by the Stanford Protein and Nucleic Acid Facility.

The 8 sub-libraries were pooled at equal final concentration (0.25 ng/µL per library) and sequenced on an Illumina NovaSeq X Plus (25 B output; 2×150 bp paired-end) flow cell with 5% PhiX spike-in by Novogene (Beijing, China), at a sequencing depth of >20k pair-end reads per cell, and 100,000 cells total. Raw FASTQ reads will be deposited at SRA and become accessible upon publication.

### Liver single-cell RNA-seq analysis

#### Read mapping, quality control, and normalization

FASTQ files were processed using the Parse Biosciences Pipeline (v.1.5.0, Parse Biosciences). This pipeline first generated a genome reference using the NCBI African turquoise killifish reference genome (Nfu_20140520, GCF_001465895.1) and its genome annotation (a GTF file manually edited to include ‘MT’ in the descriptions of genes located on the mitochondrial genome). Next, the raw sequencing data (FASTQ files) were mapped using the split.pipe function with the default parameters. The output of this function includes a count matrix and metadata for each cell.

Next, we used the Seurat package (v4.0) (*174*) and R (v4.0.3) to perform quality control and analysis, following the tutorial article provided by Parse Biosciences (*175*). A total of 120,435 cells (across 32 biological samples) were used as the input to create a Seurat object. We retained 100,338 cells with ‘nFeature_RNA’ ranging between 600 and 5000, ‘nCount_RNA’ less than 20000, and the percentage of mitochondria less than 5%. We log-normalized the filtered Seurat object using ‘NormalizeData’, selected features using ‘FindVariableFeatures’ (selection.method=‘vst’ and nfeatures= 2000), scaled the data using ‘ScaleData’, and performed Principal Component Analysis (‘RunPCA’). The ‘ElbowPlot’ function was used to identify the number of minimal dimensions needed for cell clustering (‘elbow’ around 10), and 20 dimensions were conservatively selected to perform cell clustering (FindNeighbors with dims=1:20 and FindClusters with resolution=0.5).

#### Doublet removal

To remove doublets, we used the doubletFinder package (v2.0.4) (*176*) and R (v4.0.3) to identify putative doublet cells. We stringently estimated the doublet rate to be ∼5.5% by considering the empirical doublet rate reported by Parse Biosciences (∼3%) and the estimated dissociation-related doublet rate in another study (*61*). We ran the doubletFinder function using these parameters: PCs = 1:20, pN = 0.25, pK = 0.3, nExp = round(0.055*nrow(liver@meta.data)), reuse.pANN = F, and sct = F. The cells classified as ‘doublets’ were removed from the Seurat object, resulting in 94,819 cells after doublet removal.

#### Cell type assignment

To assign each cell cluster to a cell type, we used Seurat (v5.1.0) and R (v4.1.0) to plot the cell-type marker genes identified in a recently published killifish liver single-cell RNA-seq dataset (*61*). At least 2-3 cell-type marker genes per cell type were plotted to confirm cell-type identity. We note that the relative cell-type proportion differed between our liver dataset and (*61*), for example, the cell cluster corresponding to ‘efferocytosing hepatocytes.’ This cluster shows gene expression markers from both hepatocytes and red blood cells and has been suggested to form as a result of hepatocytes engulfing lysed red blood cells (*61*). The ‘efferocytosing hepatocytes’ cluster was much smaller in our dataset than in (*61*). This difference could be due to the different liver dissociation and red blood cell removal methods used. In addition, there were three small clusters with little cluster-specific gene expression, possibly corresponding to damaged cells. We denoted these clusters as ‘unknown.’ Lastly, we visualized the cell clusters using Uniform Manifold Approximation and Projection (UMAP) and colored them by each biological variable (sex, age, and mating status).

#### Pseudobulk, Principal Component Analysis, and differential gene expression analysis

While male and female samples did not separate into distinct clusters in UMAP for most cell types, we first separated the data by biological sex to assess the effects of mating and aging more directly. For hepatocytes, we did not separate the data by sex because hepatocytes naturally separated into two distinct clusters for males and females. For pseudobulk analysis, we used the ‘AggregateExpression’ function, grouped by cell types and samples. Next, the biological conditions (sex, age, and mating status) were assigned to each biological sample.

For each cell type, the counts per gene resulting from the pseudobulk analysis were used for Principal Component Analysis (PCA). We applied the variance stabilization transformation (‘vst’ function) to the counts and then used the ‘plotPCA’ function to generate PCA plots for various PC combinations.

For differential gene expression analysis, we used the ‘FindMarkers’ function and DESeq2 analysis (test.use=’DEseq2’) to identify the genes differentially expressed between two conditions (e.g., unmated old females vs. mated old females). The genes’ average Log2 fold change (ave_log2FC), *P* values, and adjusted *P* values (Wald test corrected for multiple testing using Benjamini-Hochberg) for each comparison are available in Extended Data Table 4.

#### Variance partition analysis

We used variancePartition (v1.36.3) to identify the percentage of variance explained by age, mating status, or the age-mating interaction for each gene in a cell type. The expression count of each gene after pseudobulking was used as input, which was applied to the following formula:

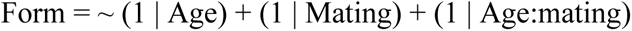

The percent variance for each variable (age, mating status, age-mating interaction) was plotted for all genes as a collective dot plot, and the average percent variance across all genes was also reported. All data related to this analysis are available in Extended Data Table 5.

#### Gene set enrichment analysis (GSEA)

We performed GSEA separately for male and female cell types. Using the output from DESeq2, we first calculated a rank score for each gene by multiplying the gene’s -log_10_(*P* value) with Log_2_FC and then sorted all the genes in descending rank scores. Genes without *P* values were removed. After identifying the human ortholog name for each killifish gene via protein BLAST (best-hit protein with BLASTp E-value > 1e-3) (*67*), we ran the enrichment analysis using clusterProfiler (v4.14.6) (*177*) and the Bioconductor annotation data package (org.Hs.eg.db v3.20.0). If multiple killifish paralogs were blasted to the same human gene, the average of the ranked scores was calculated. If no human ortholog was found, this killifish gene was removed. Multiple hypothesis testing was performed using the Benjamini–Hochberg method (p.adjust). The top gene ontology (GO) terms were graphed as a dot plot, with dot size denoting the p.adjust values and color denoting the normalized enrichment scores. All GSEA results are available in Extended Data Table 6.

For selected GO terms, we used pheatmap (v1.0.12) to generate separate heatmaps for male and female cell types. We first identified the killifish genes corresponding to the human ortholog genes that drove each GO term. If one human gene name corresponded to multiple killifish genes, we included all the killifish genes in the heatmaps. Next, for each gene, each sample’s pseudobulk-DESeq2-normalized count was converted to a z-score. The average z-score for each condition (i.e., young unmated, young mated, old unmated, and old mated) was calculated by averaging the four biological replicates. We used the average z-score for each condition as input to generate the heatmap.

### CellBiAge classifiers for liver single-cell age prediction

To evaluate the effect of mating on the age of individual liver cells, we implemented CellBiAge, a supervised machine learning pipeline (*94*). This approach uses binarized gene expression data to create a high-dimensional biomarker for cellular age.

#### Preprocessing and feature selection

We analyzed the fish liver dataset of 120,435 cells and 20,968 genes that passed quality control. To prevent information leakage, all training and test sets were preprocessed independently, and all models were evaluated on animals not seen during training. Models were built and tested separately for each sex. To derive a robust and consistent set of features, we performed the following steps. First, batch integration (*178*) was performed on the training set animals. Second, to identify Highly Variable Genes (HVGs), we identified the top 2,000 HVGs (*178*) in three separate sets: (i) the training set alone, (ii) the training set combined with the control test set, and (iii) the training set combined with the mated test set. Third, the intersection of these three gene lists was used as the final feature set for all subsequent modeling to ensure consistency. The gene expression matrix that was ultimately binarized was derived from this ‘integrated’ data after it was log-normalized and scaled.

#### All-cell models vs. cell-type models

We developed two distinct types of models to analyze the data. All-cell models were trained using all liver cells simultaneously, regardless of their specific cell-type annotation. This approach is more straightforward and robust to variations in cell type annotation. In contrast, cell-type-specific models were trained and tested exclusively on cells from a single, pre-annotated cell type (e.g., only hepatocytes). This second approach allows for a more focused biological interpretation and prevents the transcriptional signals of less abundant cell types from being obscured by more numerous cells during model training. We note that the T-cell cluster had too low number of cells to build robust models, and thus, we did not include this analysis.

#### Rotating the control training and test sets to examine the robustness of the analysis

To ensure our results were robust and not dependent on a specific train/test split, we used a rotational validation strategy. For each sex, the eight control animals were divided into four sets, with each set containing one young and one old animal dissected by two different people. We performed a 4-fold cross-validation in which, in each fold, one set was held out for testing, and the other three sets were used for training. This process was repeated until every set had been used for testing once.

Within each training fold, group-based cross-validation was used to select the optimal hyperparameters for an ElasticNet model (*179*), evaluated using the Area Under the Precision-Recall Curve (AUPRC) score to account for class imbalance (*180*). The final trained model was then applied to the previously unseen mated animals to evaluate the effect of mating on the predicted probability of a cell being ‘old.’

During this process, we noted poor performance in two specific rotations, likely due to severe class imbalance between the young and old cells in the held-out test set. For females, the model for which set 3 was held out performed poorly (test set imbalance: 2,256 vs. 4,767 cells). To correct for this, we mimicked this imbalance in the corresponding training data by excluding animal #F17_1_Vir5F during training for that specific fold. For males, the model for which set 2 was held out performed poorly (test set imbalance: 1,206 vs. 2,578 cells). Similarly, we addressed this by excluding animal #F17_2_Vir15M from the corresponding training data.

#### Statistical tests

The performance of the age prediction model was validated on the held-out control animals in each test set. A Welch’s t-test was used to confirm that the model could effectively distinguish between the age groups. Specifically, the test assessed whether the mean predicted ‘old’ probability for cells from young control animals differed significantly from that for cells from old control animals. To determine whether mating had a significant effect on predicted cellular age, we used a linear mixed-effects model. This model accounts for the non-independence of cells from the same animal by including the individual animal ID as a random effect. This approach properly models both within-animal and between-animal variation to accurately assess the impact of mating (*181*). See Supplemental Text and Extended Data Table 7 for additional results and discussion of the data and statistics for each model.

### Sample collection for RNA-seq and lipidomics cohorts

For the collection days, we prepared the harvest and fed the animals the same way as for the liver scRNA-seq experiment (see the ‘Sample collection for liver single-cell RNA-seq’ section). Fish were enrolled in three batches, 2-4 weeks apart (See Extended Data Table 8 for enrollment details and experimental metadata). Typically, four fish (two males and two females) were dissected on each harvest day (∼25 min to dissect each fish). We perfused each fish before collecting the RNA-seq and lipidomics samples to reduce the number of circulating immune cells and blood cells in the tissue. Our recent publication (*60*) described the detailed perfusion procedures. Briefly, the fish were anesthetized with 100 mg/L tricaine (pH 7.0) for 2-month-old fish or 200 mg/L tricaine (pH 7.0) for 4-5-month-old fish. A custom needle (Hamilton Syringe, custom needle, 22030-01; 30 gauge, Hubless Needle, No Hub, 30 gauge, 1.5 inches length, point style 4 (12°)) was inserted ∼1 mm into the apex of the fish heart ventricle. A syringe pump (KD Scientific, Legato 200 Series, 788200) was used for hands-free depression of the plunger of a 20-mL syringe (Fisherbrand, 14955460), which was depressed at a rate of 3.5 mL/minute to release 0.25 M EDTA in PBS (diluted from 0.5 M EDTA, pH 8.0 [Ambion, AM9261]). The needle was steadily held in place to perfuse the blood until the gill and liver were visibly pale (typically took ∼10 mL 0.25 M EDTA). After perfusion, organs were dissected on top of ice-cold Sylgard-coated Petri dishes, which were filled with wet ice and covered in plastic wrap. Each of the three operators (JC, EC, AT) dissected the same organs for most of the fish (See Extended Data Table 8 for details). Tissues were placed in 1.5 mL tubes (Fisherbrand, 02-681-320), snap-frozen in liquid nitrogen, and stored at −80°C until RNA and lipid extraction. The brain, gut, liver, and head kidneys were collected.

### Organ RNA-seq

#### RNA extraction

All samples of the same tissue were processed on the same day to reduce within-tissue batch effects. RNA extraction was performed in 2 batches for each tissue, with the sample processing order randomized and roughly equal numbers of samples from different ages and sexes in each batch. The processing order of each sample within a tissue type was randomized using the ‘Randomize’ option in Excel. After randomization, tissue samples were assigned unique numerical ‘IDs’ and split into batches of 20 samples to process simultaneously.

The RNA isolation protocol was performed as instructed by the RNeasy Mini RNA extraction protocol from Qiagen. The following protocol was used for the brain, gut, and kidney. Briefly, the sample tubes were removed from −80°C storage and placed in a box submerged in liquid nitrogen to arrange them in their respective processing order. Next, the arranged samples were transferred to a metal block on dry ice, and 100 µL cold (pre-chilled on dry ice) Zirconium beads (0.5 mm diameter, BioSpec Products, 11079105z) were added to each tube. Sample tubes were quickly transferred to wet ice, and 700 µL ice-cold QIAZOL lysis reagent (QIAGEN, #79306) was added to each tube. The sample tubes were placed between the cold (pre-chilled on dry ice) TissueLyser adaptors and homogenized on a TissueLyser II machine (QIAGEN, #85300) at 25 Hz and room temperature for 5 min. After the first round of breakage, the left and right adaptors were swapped, and the second round of breakage (25 Hz, 5 min) was initiated. Swapping the adaptors ensures that all samples experience the same degree of breakage, as the TissueLyser II machine can exhibit uneven breakage strength on each side. For the gut samples, a third round of breakage (25 Hz, 5 min) was performed to ensure full tissue lysis. After breakage, the sample tubes were placed at room temperature for 3-5 min (this step helps dissolve lipid and membrane into the organic phase). Next, all 700 µL of the lysate was transferred to 200 µL chloroform in 1.5 mL DNA LoBind tubes (Eppendorf, 0030108051), vortexed for 15 sec, and incubated at room temperature for 2-3 min.

Next, samples were centrifuged at 12,000 x g at 4°C for 15 min. For each tube, 350 µL total of aqueous phase (use a p200 pipette to transfer 175 µL twice to ensure no white interface debris was transferred) was transferred to 350 µL 70% ethanol (200 Proof, Gold Shield Distributors, #412804, diluted in nuclease-free water) in another 1.5 mL DNA LoBind tube. The tubes were inverted ∼10 times to mix and briefly centrifuged to collect all liquid. For each sample, the ethanol-lysate mixture was transferred to an RNAeasy Mini spin column (reagent from QIAGEN, #74536) and centrifuged at 10,000 x g at room temperature for 30 seconds (all subsequent wash steps used this centrifugation condition). The column was washed with 350 µL RW1 (reagent from QIAGEN, #74536) and treated with DNase I by incubating it in 80 µL DNase I solution (prepared as instructed by the manufacturer) at room temperature for 15 min. Next, 350 µL RW1 was added directly to the column and centrifuged. The columns were washed twice with 500 µL RPE buffer (reagent from QIAGEN, #74536), with a 2-min centrifugation step for the final RPE wash. RNA was eluted in 50 µL nuclease-free water (Invitrogen, #10977023) in 1.5 mL DNA LoBind tubes, aliquoted (5 µL each), and stored at −80°C. For RNA samples with A260/230 ratios lower than 1.8 (measured by Nanodrop), the RNA was ethanol-precipitated as follows: 25 µL nuclease-free water, 135 µL nuclease-free 100% ethanol, and 4.5 µl RNase-free 3M sodium acetate, pH 5.5 (Invitrogen, AM9740) were added to the RNA. After inverting 10 times to mix, the samples were incubated at −20°C overnight. Next, the samples were centrifuged at 21,000 x g at 4°C for 20 min, washed with 500 µL 70% ethanol, centrifuged again at 21,000 x g at 4°C for 10 min, and fully air dried at room temperature (∼15 min). Lastly, the dried RNA pellets were resuspended in 15-25 µL nuclease-free water by incubating at room temperature for ∼30 min with gentle vortexing. RNA concentration was measured using a Quant-iT RNA BR kit (Thermo Fisher, Q10213) on a Varioskan LUX Multimode microplate reader (Thermo Fisher, VL0000D0). Two to four RNA samples from each tissue were randomly selected to check RNA quality using the Agilent 2100 Bioanalyzer and Agilent’s RNA Nano kit (Agilent, 50671511) by the Stanford Protein and Nucleic Acid Facility.

#### cDNA library generation and sequencing

Sequencing libraries were prepared as described in (*60*). To achieve high pipetting accuracy, the Dragonfly (SPT Labtech) or Mosquito HV (SPT Labtech) robotic liquid handlers were used to perform most pipetting steps. Briefly, RNA sample concentrations were normalized to 25 ng/µL. Next, 2 µL of each sample was used in the cDNA first-strand synthesis reaction, followed by 9 cycles of cDNA library amplification. A portion of the full cDNA library volume (6 µL) was cleaned using AMPure XP beads (Beckman Coulter, A63881) at a 0.7X ratio, following the manufacturer’s guidelines, including two washes of 10.7 µL 80% ethanol (200 Proof, Gold Shield Distributors, #412804; diluted in nuclease-free water) and elution in 4.5 µL nuclease-free water (Invitrogen, #10977023). The concentrations of the amplified cDNA libraries were measured using a Quant-iT dsDNA HS Kit (Thermo Fisher, 33120), and a subset of libraries was also measured using the Agilent 2100 Bioanalyzer and Agilent’s DNA high-sensitivity kit (Agilent, 50674626) by the Stanford Protein and Nucleic Acid Facility.

Next, we used the Nextera XT DNA Library Preparation Kit (Illumina, FC-131-1096) and the IDT for Illumina DNA/RNA UD Index Sets A-D (Illumina, 2002713, 20027214, 20042666, 20042667) to prepare sequencing libraries, following the manufacturer’s instructions except by reducing all reactions by half. The Illumina Index Sets A-D were converted into a 384-well format. For tagmentation, 0.5 ng of the cDNA (2.5 µL total) was mixed with 5 µL TD buffer and then 2.5 µL ATM buffer, incubated at 55°C for 5 min, and cooled to 10°C. A total of 2.5 µL NT buffer was added to stop the reaction, followed by a 5-min incubation at room temperature. The cDNA library was indexed and amplified for 12 cycles in a PCR reaction containing 10 µL tagmented DNA, 5 µL dual indices, and 7.5 µL NPM buffer. Two 12.5 µL aliquots were made from the indexed cDNA library (25 µL total), and each aliquot was purified using 22.5 µL AMPure XP beads as described above. A total of 11 µL Buffer EB (QIAGEN, 19086) was sequentially incubated with the AMPure beads of each aliquot, such that both aliquots were re-pooled (the final elution volume was 10 µL). The concentration of each indexed library was measured using the Quant-iT dsDNA HS Kit. A subset of libraries from each tissue was analyzed on a Bioanalyzer to obtain their average size distributions. The indexed libraries were pooled equimolarly, purified using AMPure beads at a 1.8x ratio, and sequenced by Novogene (Novogene, China) on 2 lanes of an Illumina NovaSeq X 25B flow cell (2×150 bp paired-end) with 10% PhiX spike-in for each lane, at a target sequencing depth of >20 million pair-end reads per sample. Novogene performed base calling, demultiplexing, and FASTQ file generation.

### Organ RNA-seq data analysis

#### Read mapping and quality control

Raw FASTQ data were processed using Trim-galore v0.5.0 and aligned to the African turquoise killifish reference genome (Nfu_20140520, GCF_001465895.1) using STAR v2.7.10b (*182*) with the default parameters. We used samtools v1.16.1 (*183*), with the parameters of MAPQ < 255 (‘samtools view -q255 -b’), to remove the reads mapped to multiple genomic regions and input the uniquely mapped reads into the ‘featureCounts’ program (with the default parameters) from subread v2.0.6 (*184*) to generate the read counts for each gene.

We used ‘gene expression connectivity’ to detect outliers from the WGCNA package v1.73 (*185*). This method computes correlations among all samples, derives network connectivity for each sample, standardizes the connectivity scores, and identifies samples with Z-scores below −2 as outliers. We identified one brain sample as an outlier and removed it from subsequent analyses.

#### Principal Component Analysis (PCA)

All analyses were performed in R v4.3.3. To perform PCA, we first created a *DESeqDataSet* object of all samples using DESeq2 v1.42.1 (*186*). After filtering out genes for which the sum of the raw counts across all samples was 0 (15 genes), we applied variance stabilizing transformation (‘vst’) on the raw counts of the *DESeqDataSet* object and visualized using the ‘plotPCA’ function. To generate the PCA plot for a given tissue, we subsetted to that tissue and performed variance stabilization of the raw counts before running PCA, as described above.

#### Variance partition analysis

Using the package variancePartition v1.36.3 (*187*), we calculated the proportion of variance explained by the covariates sex, age, mating status, and interactions of age:sex, age:mating status, and sex:mating status on a per-tissue basis. For brain, gut, and head kidney, the count matrix of each tissue was subsetted to include only genes with a row sum of 40 (i.e., assuming at least 1 count per sample). The formula ∼ (1 | sex) + (1 | age) + (1 | mating status) + (1 | sex:age) + (1 | sex:mating status) + (1 | age:mating status) was used to explore the respective contributions of these variables to variance. The results of the variancePartition analyses for each tissue can be found in Extended Data Table 8 and plotted as a dot plot.

#### Differential gene expression analysis, pathway analysis

We first subsetted the data into male and female samples and then performed differential expression (DE) analysis using DESeq2 for each tissue. We used a compound variable called ‘Condition’ that specified both age and mating status for each sample. To compare between the young and old samples for a given mating status (unmated or mated), the young samples were used as a baseline of comparison (i.e., a positive Log_2_ fold change means a gene upregulated with age). To compare the mated and unmated samples for a given age (young or old), the unmated samples were used as the baseline (i.e., a positive Log_2_ fold change means a gene upregulated in the mated samples). The significantly changed genes were defined as those with p.adj < 0.05 (Wald test, followed by Benjamini-Hochberg multiple hypothesis correction). Z-scores corresponded to the values of the Wald statistics (i.e., the ‘stat’ term of the DESeq2 output), which is defined as Log_2_ fold change divided by the standard errors of the Log_2_ fold change (i.e., the ‘lfcSE’ term of the DESeq2 output). GSEA and generation of heatmaps were performed as described in the scRNA-seq section. The DE analysis and GSEA results of each tissue are available in Extended Data Tables 9 and 10.

### Liver lipidomics

#### Sample preparation and lipid extraction

Tissue collection is described in the ‘Sample collection for RNA-seq and lipidomics’ section. Because killifish livers are relatively small (less than 0.5 cm in diameter for young fish), consistent dissection into sub-regions was not feasible. Therefore, each liver was homogenized in its entirety to generate a tissue lysate, which was then aliquoted. To lyse the livers, the sample tubes were arranged based on the processing order in liquid nitrogen, transferred to dry ice, and 100 µL cold (pre-chilled on dry ice) Zirconium beads (0.5 mm diameter BioSpec Products, 11079105z) were added. Next, while on wet ice, the sample tubes received 200 µL ice-cold lysis buffer, prepared by dissolving 1 tablet of cOmplete mini EDTA-free protease inhibitor cocktail (Roche, 11836170001) and 100 µL protease/phosphatase inhibitor cocktail (Cell Signaling Technology, CS5872) in 2 mL of nuclease-free PBS. This lysis buffer was used to obtain lysate for multi-modal analysis (RNA, proteins, and lipids). Next, the sample tubes were placed between the cold (pre-chilled on dry ice) TissueLyser adaptors and homogenized on a TissueLyser II machine (QIAGEN, #85300) at 25 Hz and room temperature for 1 min. The samples were quickly transferred to a metal block on ice in a room at 4°C. Each sample was then aliquoted three times, with 55 µL of the lysate transferred to each of the following tubes: one DNA LoBind tube (Eppendorf, 022431021) and two Protein LoBind tubes (Eppendorf, 022431081), which were immediately frozen in liquid nitrogen and stored at −80°C for lipidomics. Lipid extraction was performed in two batches using a biphasic methanol/tert-butyl methyl ether (MTBE)-based method adapted from a published protocol (*65*). To minimize inter-batch technical variation, we employed the following strategies: (1) a balanced design in which one animal from each condition (sex, age, and mating status) was assigned to each extraction group (8 samples per group; 5 groups total); (2) randomization of extraction order within each group; and (3) parallel processing by two researchers (J.C. and M.H.), each handling an equal number of samples side by side to minimize subtle differences in execution (e.g., time of day, vortex speed). Three commercial internal standard mixtures were added to the extraction methanol (Sigma-Aldrich, 34860, HPLC grade) for quality control and extraction efficiency assessment: EquiSPLASH (Avanti, 330731; a mixture of 13 deuterated lipid standards at 100 µg/mL each; diluted 1:150 in extraction methanol), Cer/SphII (Avanti, LM6005; a mixture of 9 ceramide and sphingoid standards at 25 µM each; diluted 1:150 in extraction methanol), and cholesterol-d7 (Avanti, LM4100; 513.59 µg/mL; diluted ∼1:770 in extraction methanol). Liver lysate aliquots were thawed on ice, and 40 µL per sample was transferred to a 2-mL tube (Axygen, MC-200-C-S). Then, 300 µL the methanol/internal standards solution was added, mixed by pipetting five times, and vortexed vigorously at room temperature for 20 seconds. Next, 1 mL of ice-cold MTBE (Sigma-Aldrich, 650560, HPLC Plus grade) was added, vortexed vigorously for 20 seconds, and the tubes were incubated on ice for 30 min with periodic vortexing (10 seconds every 5 min). After incubation, 250 µL ice-cold LC/MS-grade water (Fisher Scientific, W6-1) was added, and the samples were vortexed for 1 min and centrifuged at 14,000 × g at room temperature for 10 min. The upper (organic) layer (800 µL) was transferred to an amber glass vial (Agilent, 5183-2073) and dried under a stream of N₂ gas using a TurboVap LV (Biotage, 415000) at 0.5 L/min for 15 min, followed by 1.0 L/min for 15 min, or until no visible liquid remained. Dried extracts were stored at −80°C. Protein concentration of each sample was determined using a Pierce^TM^ BCA Protein Assay Kit (Thermo Scientific, 23227) after diluting 2 µL of the lysate in 98 µL RIPA buffer (Thermo Scientific, 89900) to fall within the linear range of the assay. Dried extracts were reconstituted in 300 µL methanol:toluene (9:1, v/v; Fisher Scientific) and centrifuged at 18,407 × g for 5 min. Then, 180 µL of the resulting supernatant was transferred to an amber LC vial equipped with a 300-µL insert. Five pooled quality control (QC) samples were prepared by combining equal-volume aliquots from all study samples and were injected periodically after every 10 study samples. Two solvent blank samples were also prepared and analyzed—one at the beginning and one at the end of the analytical run—to monitor background signals and potential contamination or carryover. The lipidomics metadata were included in Extended Data Table 11.

#### LC-MS/MS-based lipidomics

Untargeted lipidomic analysis was performed on a Thermo Fisher Scientific Ultimate 3000 UHPLC system coupled to a Q-Exactive Orbitrap mass spectrometer (Thermo Fisher Scientific, Rockford, IL, USA). A 5 µL aliquot of each reconstituted sample was analyzed in both positive (ESI+) and negative (ESI−) electrospray ionization modes, using a method adapted and optimized from previously published studies (*188–190*). The UHPLC column compartment and autosampler temperature were maintained at 40°C and 10°C, respectively. Prior to analyzing study samples, the Q-Exactive mass spectrometer underwent thorough maintenance, including cleaning of the ion source and ion transfer tube, followed by instrument evaluation and calibration in both ionization modes. System suitability was confirmed by analyzing multiple blank and QC samples before initiating the analytical run. Full-scan MS1 data were acquired for all samples. Data-dependent acquisition MS/MS (ddMS2) was performed on a subset of QC samples for lipid identification and confirmation. An iterative ddMS2 approach was additionally applied to enhance lipidome coverage (*191*). MS parameters were as follows: full-scan MS1 resolution, 70,000 FWHM (at m/z 200, 3.7 Hz); ddMS2 resolution, 17,500 FWHM (at m/z 200); scan range, m/z 200–1200. Mass accuracy was maintained at <1 ppm RMS error using internal calibration and <3 ppm RMS error using external calibration. Automatic gain control (AGC) targets and maximum injection times were 1 × 10⁶ and 100 ms for full-scan MS1, and 1 × 10⁵ and 50 ms for ddMS2, respectively. The top 10 most abundant precursor ions were selected for fragmentation (TopN = 10). Normalized collision energies (NCE) were 25 and 35 in positive mode, and 20, 40, and 60 in negative mode. A heated electrospray ionization (HESI) source was used, with spray voltages of +3.5 kV (positive mode) and −3.0 kV (negative mode). Capillary and auxiliary gas heater temperatures were 320°C and 300°C, respectively. Sheath gas and auxiliary gas flow rates were 40 and 10 arbitrary units, respectively, and the S-lens RF level was set to 55.

#### Untargeted Lipidomics data preprocessing

Our lipidomics analysis pipeline followed recommendations from Lipidomics Standard Initiative (https://lipidomicstandards.org/), recent publications, and the Gitbook ‘Omics Data Visualization in R and Python (https://laboratory-of-lipid-metabolism-a.gitbook.io/omics-data-visualization-in-r-and-python) (*192*). Raw LC-MS/MS data were converted to .mzML format using ProteoWizard (*193*). Data from each ionization mode were processed separately as single batches using MassCube v1.2.8 for peak extraction, alignment, quantification, and annotation (*194*). Processing parameters were optimized for the instrument configuration and analytical method; detailed parameters are provided in the Extended Data Table 11. Lipid identification was performed by matching acquired MS/MS spectra against the MassBank of North America, the Global Natural Product Social Molecular Networking (GNPS), and the MS-DIAL metabolomics MSP spectral kit (*195–197*). The positive-mode dataset contained 71,308 features, and the negative-mode dataset contained 43,392 features. Preprocessed data for both ionization modes are available in Extended Data Tables 12 and 13.

#### Quality control, data filtering, batch effect assessment

Positive-mode and negative-mode data were analyzed separately. To focus on high-quality detections, we filtered features using three key quality-control parameters. These parameters include: 1) in-source fragments, which are a major source of false feature annotations; 2) Gaussian similarity, which measures how similar a given chromatographic peak resembles a perfect Gaussian function; and 3) noise score, which quantifies how smooth the mass spectrometry signal is over time. We removed features that 1) had more than 50% missing values (zeros) across QC and study samples, 2) were in-source fragments (‘is_in_source_fragment’ was true), and 3) had a Gaussian similarity ≤ 0.7 and a noise score ≥ 0.4. Lastly, we retained only detected lipids for which the average intensity in the QC samples was at least 3-fold higher than the maximum intensity of the two blank samples (*198*). After this filtering step, 4,268 and 1,459 detected lipids remained in the positive-mode and negative-mode datasets, respectively. We also identified ‘uniquely annotated’ lipids by keeping the detection with the highest Gaussian similarity for any lipid detected more than once (‘duplicates’). We provided the filtered results in Extended Data Table 14.

For batch effect assessment, the raw intensity of detected features (preprocessed data) was Log_2_-transformed. We performed Principal Component Analysis (PCA) to assess the effects of technical batches, including lipid extraction batch and extraction group, extraction operators, and animal cohorts. We did not observe batch-associated clustering in PC1 (20.6% variance explained) and PC2 (14.46% variance explained), although there was some extraction batch effect in PC3 (6.73% variance explained). We provided the batch-effect assessment PCA results in Extended Data Table 15.

#### Lipid species quantification

In this study, we performed a ‘Level 2’ quantification for annotated lipids. As defined by Lipidomics Standards Initiative, a ‘Level 2’ quantification accounts for sample processing variation using the lipid class internal standard co-ionized with the analyte. We first normalized the filtered ion counts by the total protein input of a given sample. The total protein input was calculated by multiplying the protein concentration of each sample (measured by BCA assay in µg/ml) by the total lysate volume used for extraction (0.04 ml). For the QC samples, we used the average protein input across all 40 study samples as their protein input, since the QC samples were pooled from equal volumes of each study sample. Next, for uniquely annotated lipids, we performed a ‘class normalization,’ which attempted to account for sample processing variation by normalizing the ion counts to internal standards that eluted at similar retention times and were therefore subjected to the same degree of competition for ionization in the source. We noted that because there was only one internal standard per lipid class, the quantification accuracy was affected by matrix effects and may not fully reflect the distinct MS responses of lipids with different fatty acyl chains. Most annotated lipids were named after their lipid class, followed by information of the fatty acyl chains (e.g., PC(16:3/16:0)). For the lipids that lacked such standard nomenclature, we manually annotated them by cross-referencing to public databases (e.g., LIPID MAPS, PubChem). We used the following lipid ions for quantification: [M+H]+ or [M+H-H2O]+ for ceramides (Cer); [M+H]+ or [M+NH4]+ for diglyceride (DAG) and triglyceride (TAG/TG); [M+H]+ or [M+Na]+ for phosphatidylcholine (PC) and sphingomyelin (SM); [M+H]+ for cholesterol, LysoPC/LPC and LysoPE/LPE; [M+H]+, [M+NH4]+, or [M+Na]+ for phosphatidylethanomine (PE); and [M-H]- for phosphatidylglycerol (PG), phosphatidylinositol (PI), and phosphatidylserine (PS). To estimate the concentration of each annotated lipid in each sample, we divided the protein-normalized count of each lipid by its associated class standard (‘iSTD’ count). For example, for PC(16:3/16:0), we divided its count by the count of its class standard, PC(15:0/18:1(d7)). We then multiplied this class-normalized count by the internal standard concentration in the reconstituted samples (the final unit for each class-normalized lipid is nmol per mg protein input). We provided the class-normalized lipid concentrations (for a total of 313 lipids) in Extended Data Table 14.

#### PCA, differential expression analysis, heatmaps

Principal Component Analysis (PCA) was performed using the ‘class-normalized’ concentrations for the 313 uniquely annotated lipids. To perform differential expression analysis for each uniquely annotated lipid, we used the ‘class-normalized’ concentrations as the input. We applied a Welch’s t-test, which does not assume equal variances and has been used in prior lipidomics studies (*65, 199*), followed by Benjamini-Hochberg multiple hypothesis correction. This analysis generated all pairwise comparisons among the 8 conditions across two biological sexes (males vs. females), two age groups (young vs. old), and two types of mating status (mated vs. unmated). For quantification at the lipid-class level, we first summed the abundance of all the uniquely annotated lipids in that class. Then, for each lipid class, we performed the Mann-Whitney *U* test to compare across conditions. To generate the heatmaps, we first averaged the class-normalized lipid concentrations for each lipid across the five animals (biological replicates) per condition, performed a Log_2_ transformation, z-scaled each lipid (row-wise) across all conditions, and then hierarchically clustered the lipids. We provided the differential expression data in Extended Data Table 16.

### Protein sequence alignment and AlphaFold analysis

For protein sequence similarity, we aligned the protein sequences encoded by human (*IRF9* or *TCEB3*) and killifish (*irf9* or *LOC107396610*) genes using Clustal Omega (https://www.ebi.ac.uk/jdispatcher/msa/clustalo) (*200*). For structural prediction, we performed the ‘Search’ function using the killifish protein sequence in the AlphaFold Protein Structure Database (https://alphafold.ebi.ac.uk/). The orthologous human sequence was identified using the default MMseqs2 clustering algorithm in the AlphaFold database. We imported the PDB files for both the killifish and human proteins into the ChimeraX software (v1.9rc202412100036) (*201–203*), adjusted the display color of the protein sequences and orientation, and performed superimposition between the killifish and human sequences using the ‘Matchmaker’ function (parameter: ‘best pairing of chains’, reference sequence is the human sequence). The protein images were exported and saved as ‘png’ files (resolution: 2000 x 1920, supersample: none). The protein sequences and alignment files are included in Extended Data Table 17.

### Acetaminophen toxicity test

The acetaminophen treatment protocol was adapted from a zebrafish study (*119*). Approximately 900 mL of acetaminophen solution at defined concentrations (see below for treatment details) was distributed into a 1.8 L tank before fish placement, and the tanks were maintained in non-circulating conditions at room temperature in the animal facility. Fresh acetaminophen solution was prepared daily, and a full water exchange was performed in each tank every 24 hours. After the daily water change, each fish would receive 15 mg of dry food pellet (‘reduced feeding regimen’), which was half of the daily feeding quantity for other experimental fish to minimize waste in non-circulating tanks. The fish were monitored for physiological responses and death for ∼72 hours.

To assess the sensitivity of adult killifish to acetaminophen, we treated unmated male and female killifish (3 to 4 months old) with varying concentrations of acetaminophen (Sigma-Aldrich, A7085) dissolved in system water: 0 mM (system water only), 10 mM, 25 mM, and 50 mM acetaminophen. The unmated females were maintained in 9.8 L group tanks at a density of 6-10 fish from sexual maturity to the time of drug treatment, whereas unmated males were singly housed in 1.8 L or 2.8 L tanks. All animals were randomly assigned to each treatment group. Fish that survived for more than 72 hours were harvested for molecular analysis.

To compare the acetaminophen sensitivity of unmated and mated females, we enrolled killifish females in mating cohorts as described in the ‘African turquoise killifish husbandry’ section. At 17 weeks of age (∼4.5 months), females (after lifelong mating) underwent one more mating session. The next day (∼24 hours post mating), females were randomly assigned to the ‘no drug’ control group (0 mM; only system water) or the treatment group (20 mM acetaminophen in system water). We chose 20 mM acetaminophen for the treatment group because this concentration represented a sensitized condition for lethality in most unmated old females between 24 hours and 48 hours. Each tank received a daily water exchange and a reduced feeding regimen as described above. The lethality data were included in Extended Data Table 18.

### Tissue preparation for staining

For experiments related to quantifying immune cell abundance in killifish livers, fish were enrolled in two independent cohorts, ∼9 months apart (See Data S8 for enrollment details and experimental metadata). The fish were sacrificed by incubating them in system water supplemented with ice (0°C) for 2-3 min for rapid cooling. The fish were dried and placed on an ice-cold Sylgard-coated Petri dish filled with wet ice and covered in plastic wrap. The skin covering the gills was excised using scissors. An incision was made from the Basibranchial cartilage, along the midline of the abdominal region, to the anus. The abdominal muscles on both sides of the fish were removed to expose the internal organs. Typically, it took 6-10 min to dissect each fish. The dissected fish were incubated in 6-10 mL 4% PFA in PBS (Santa Cruz, CAS 20525-89-4) on a rocking platform at 4°C for ∼20-24 hours. The PFA-fixed fish were washed 3 times by incubating in ∼12 mL PBS on ice for 1 hour during each wash (residual PFA was quickly rinsed out using ∼12 mL cold PBS before the first PBS wash) and placed in ∼12 mL fresh PBS to store at 4°C until dissection (up to 4 days). The fish were dissected in ice-cold PBS to separate the organs. To store the organs, they were placed in methanol by incubating in the following series of solutions for ∼10 min each on ice: 1 mL 66% methanol:33% PBS, 1 mL methanol, and 1 mL methanol. The dissected organs were stored in 1 mL methanol at −20°C until sample sectioning.

To prepare tissue cryosections, the tissues stored in methanol were progressively rehydrated by incubating (10 min each) in 1 mL of 75% methanol:25% PBS, 1 mL of 50% methanol:50% PBS, 1 mL of 25% methanol:75% PBS, and twice in 1 mL of PBS at room temperature. Next, the tissues were incubated in 1 mL of nuclease-free 30% sucrose solution (30 g sucrose in 100 mL PBS) at 4°C for approximately 20-24 hours. The tissues should fully sink to the bottom of the sucrose solution. For embedding, each organ was dried using Kimwipe (Fisher Scientific, 06-666A), placed in a Peel-A-Way mold (Fisher Scientific, 2219) or a Tissue-Tek cryomold (15 mm x 15 mm x 5 mm, Sakura Finetek, 4566) filled with optimal cutting temperature (OCT) compound (Fisher Scientific, 23-730-571). Each liver was oriented along the anterior-posterior axis with the concave side down and frozen in ethanol-dry ice bath for ∼15 min before storing at −80°C. The liver samples were sectioned to a thickness of 10-25 µm (ovary samples, 50 µm) using a M35 Ultra Low-Profile Microtome Blade (Mercedes Scientific, FIS 3053835) in a cryostat (Leica, CM3050 S) at −18 to −20°C objective and chamber temperatures. All the tissue sections were captured on the Superflost Plus slides (Fisher Scientific, 12-550-15), dried at room temperature for ∼15 min, and stored at −20°C.

### Hybridization chain reaction (HCR) for RNA *in situ* hybridization

#### HCR probe design

All HCR probe sequences and the associated killifish gene names are listed in Extended Data Table 2. We designed HCR probes for each gene using a custom-made Python script Probe_Maker.py, which takes the longest annotated mRNA sequence from the NCBI killifish genome (Nfu_20140520, GCF_001465895.1) and breaks it into 20-basepair (bp) segments with a 2-bp gap between each segment, starting from the 5’-end of the transcript. Two adjacent 20-basepair segments are assigned as a pair (e.g., Pair 1: segment 1 and 2; Pair 2: segment 3 and 4; etc), and a hairpin sequence (e.g., B1h1 or B1h2 for using the hairpin set B1) is added to the first and second segments in each pair, respectively. These two new sequences form an initiator sequence pair. In total, each transcript has 20–24 initiator sequence pairs (a total of 44–48 initiator sequences), which were purchased from IDT as oPool Oligo pools (50 pmol, then resuspended to 0.5 pmol/μL in RNase-free TE 8.0 [Ambion, AM9858]). The hairpins were purchased from Molecular Instruments (HCR amplifiers), with the B1 hairpins conjugated to Alexa Fluor 647, the B3 hairpins to Alexa Fluor 546, and the B5 probes to Alexa Fluor 488.

#### Staining procedure

To stain similar anatomical regions of each liver, we performed HCR on liver cryosections from the same slide quantiles across animals. To determine the slide quantile of a given slide, we divided the cumulative tissue thickness up to this slide by the total thickness of the whole liver. For example, for animal 1, we used slide 45, which corresponded to the 50^th^ percentile of its liver. But for animal 2, which had a larger liver, we used slide 55, corresponding to the 50^th^ percentile of this liver. To sample a wider range of liver regions for the acetaminophen experiments and for ovaries, we put tissue sections of three or four animals (biological replicates) on the same slide rather than using the same slide quantile across animals.

All HCR-related procedures were conducted in RNase-free conditions. Thawed tissue sections were incubated in 1 mL PBS at room temperature for ∼10 min to dissolve OCT, then OCT was gently washed off the slides with 1 mL PBS. Next, each slide was washed 4 times with ∼5-min incubation of 500 μL PBST (PBS supplemented with 0.1% Tween 20 [Bio-Rad, 170-6531] diluted in nuclease-free water) and then pre-hybridized using 500 μL hybridization buffer (Molecular Instruments, HCR tissue probe hybridization buffer) at 37°C in a humidified chamber for 30 min–1 hour. The different probe sets were mixed with the hybridization buffer, with 2 μL of each 0.5 pmol/μL probe added to 100 μL hybridization buffer. Approximately 65–100 μL of the mixture was added to each slide and incubated at 37°C in a humidified chamber for 17–19 hours (a coverslip was added to each slide to reduce evaporation). The slides were washed with probe Wash Buffer (Molecular Instruments, HCR tissue probe wash buffer), 15 min each, in the following series of solutions: 500 μL 100% Wash Buffer, 500 μL 75% Wash Buffer:25% 5x SSCT (0.1% tween, made using nuclease-free water and 20x SSC Invitrogen, AM9763), 500 μL 50% Wash Buffer:50% 5x SSCT, 500 μL 25% Wash Buffer:75% 5x SSCT, and 500 μL 5x SSCT (optional). The slides were pre-amplified with ∼200 µL Amplification Buffer (Molecular Instruments, HCR amplification buffer) at room temperature for 30 min–4 hours. Hairpin solutions were prepared per the manufacturer’s instructions (each hairpin was incubated at 95°C for 90 sec, then cooled down at room temperature in the dark for ∼25 min). Amplification solution was prepared by mixing 2 μL of each harpin (3 µM) to 100 μL Amplification Buffer, and 70-100 µL of the amplification solution was added to each slide, followed by incubation at room temperature for 18–22 hours. After incubation, the amplification solution was removed and washed with two 30-min washes of 500 μL DAPI (0.01 mg/mL) diluted in 5x SSCT and one 5-min wash of 500 μL 5x SSCT. Antifade reagent (Thermo Fisher Scientific, Prolong Gold Antifade Mountant, P36934) was added to each slide, which was then covered by a 22×50 mm, No. 1.5 coverslip (Fisher Scientific, 12-544-DP) and sealed with nail polish. The slides were curated in the dark for at least 16 hours before imaging.

#### Confocal microscopy

Each slide was imaged as 16-bit images on an inverted confocal microscope (Zeiss LSM 900), operated with a ZEN blue software (3.0) using the following conditions: 1) 40x objective (Zeiss plan/Apocromat/40X/1.40, oil) and Zeiss Immersol oil 518F (Zeiss, 4Y00-R0DY-1007-3VF3); 2) z-stacks: 9-15 slices with a 0.75-μm step size; 3) pin hole: 32 μm; 4) channels in this order: Alexa 546 (577/603, 1.5% laser intensity, master gain: 775V, detector offset: 256, detector digital gain: 1.0); Alexa 488 (493/517, 2.5% laser intensity, master gain: 650V, detector offset: 256, detector digital gain: 1.0); DAPI (353/465, 0.5% laser intensity, master gain: 650V, detector offset: 256, detector digital gain: 1.0); and Alexa 647 (653/668, 8.0% laser intensity, master gain: 650V, detector offset: 512, detector digital gain: 1.0). Each z-slice was imaged in the four channels before moving to the next z-slice. Anatomically, unlike mammalian livers, fish livers are not known to exhibit zonation (*204*). Therefore, we typically imaged 4 evenly spaced fields of view on each liver section, with the coordinates marked by dividing the section’s x- and y-dimensions by 4. For the ovary, we selected 3-4 fields of view with oocytes (*buc+* signal) to image.

### Image analysis and quantification

#### Quantification of immune cells

Macrophages and neutrophils can exhibit asymmetric cell shapes and membrane extensions projecting toward other cells (*100*). Therefore, canonical cell segmentation based on DAPI masks did not work well. Maximum projection in the z-direction could also lead to mis-assignment of *MARCO* and *mpx* mRNAs to non-immune cells. Given these limitations, all macrophage and neutrophil cell counts were performed manually by sequentially checking each z-plane. For a cell to be counted as ‘*mpx+*’ or ‘*MARCO+*’, this cell needed to have at least 4 *mpx* mRNAs or 4 *MARCO* mRNAs, respectively, within ∼3 µm of the nucleus in the x, y, and z directions. Typically, a neutrophil expressed many more than 4 *mpx* transcripts and a few *MARCO* transcripts. A macrophage expressed many more than 4 *MARCO* transcripts and did not express *mpx* transcripts. In the first and last 2-3 z-planes, the full volume of some immune cells was not captured, but *mpx* and *MARCO* mRNAs were present. To determine how to quantify these ‘edge cases’ accurately, we acquired thicker z-stacks for some images (21 z-slices instead of 9 z-slices). We found that a threshold of 4 or more mRNA spots typically identifies these ‘edge’ immune cells correctly (including *CD45+* leukocytes).

We classified an ‘*mpx+*’ cell as a neutrophil. The number of macrophages was calculated by subtracting the number of neutrophils from the total number of ‘*MARCO+*’ cells (since both macrophages and neutrophils expressed *MARCO* mRNAs). We summed across four fields of view to find the total number of neutrophils and macrophages for each animal. To calculate the proportion of neutrophils and macrophages, we first estimated the total cell number using QuPath (v0.5.1, https://qupath.github.io/). We projected each image by maximum intensity in the z-direction using a custom-made FIJI script. Max-projected images were then loaded into QuPath and segmented using a nuclear mask based on the DAPI signal (DAPI threshold: 2000), followed by a 3-µm expansion of the DAPI mask to define cell boundaries. For each animal, the total cell counts were summed across four fields of view. The proportion of neutrophils or macrophages was calculated by dividing the total number of each cell type by the total cell count for that animal. We note that we were unable to distinguish intravascular immune cells from infiltrated immune cells in these experiments because the animals were not perfused to match the scRNA-seq harvest conditions. Data were plotted in GraphPad Prism (v10). The immune cell quantification data were included in Extended Data Table 19.

#### Quantification of IRF9, TLR5, IL1β, and TNFα mRNAs

All images were batch-processed in FIJI using a custom-made script. First, the full z-stack of each image was projected by maximum intensity in the z-direction (note: for *TLR5,* only three z-slices were used for projection due to the high density of mRNAs). Projected images were loaded into QuPath. Cells were segmented using a nuclear mask generated from the DAPI signal (DAPI threshold: 2000), and a 3-µm expansion of the DAPI mask was used to define cell boundaries. Red blood cells, which exhibit strong autofluorescence across all channels, were manually removed to avoid false-positive subcellular spot detection.

Next, we used the QuPath subcellular detection function to detect each type of mRNA using specific parameters (see Data S9 for details). Every cell was visually inspected to verify that the detected spots matched manual counting. A small number of false-positive spots were manually removed (these spots typically occur in regions containing red blood cells). The number of a specific type of mRNA per cell and the total cell count for each image were recorded. A non-immune cell was identified as one with fewer than 4 *CD45* mRNAs. The average number of *IRF9* or *TLR5* mRNAs was calculated by dividing the total mRNA counts of all the non-immune cells by the total number of non-immune cells. ‘*TNFα+*’ cells were defined to have at least 5 *TNFα* mRNAs per cell; ‘*IL1β*+’ cells, at least 4 *IL1β* mRNAs per cell. The quantification data were included in Extended Data Table 20.

#### Pro-inflammatory marker selection criteria

To define pro-inflammatory markers in the killifish liver, we looked for genes with possible pro-inflammatory roles using two main criteria: 1) evolutionary conservation, where the killifish gene has mammalian orthologs with pro-inflammatory activity, and 2) expression upregulation in aging and liver injury (i.e., insults on the liver induce this gene). The killifish gene *irf9* fits these criteria. It had ∼33% sequence identity and protein domain similarity (**fig. S4A**) to the orthologous human *IRF9* gene (hereafter referred to by the human gene name), which has known pro-inflammatory roles in mammals. The Killifish Aging Atlas (*60*) revealed that *IRF9* showed a trending upregulation with age in unmated female livers (**fig. S4B**). *IRF9* was upregulated in non-immune liver cells upon treatment with a non-lethal dose of acetaminophen, a liver toxin (see **Fig. 4C**), suggesting that liver injury induced *IRF9* expression. Given that *IRF9* upregulation was associated with liver aging and damage, we measured *IRF9* expression in non-immune liver cells.

#### Quantification of TCEB3 mRNAs in the tissue sections of the ovary

We first performed z-projection by maximum intensity using the full z-stack of each image in FIJI (2.9.0). Due to a weak DAPI signal in the oocytes, cell segmentation in QuPath was not robust. Instead, we performed manual segmentation by drawing boundaries around oocytes, which are regions with positive signal for *buc* mRNAs (threshold: >5000). Next, we measured the mean intensity of *TCEB3* signal in each oocyte (i.e., *buc+* region) and calculated the average for all oocytes imaged across all (typically 4) fields of view (‘average oocyte *TCEB3* signal’). In addition, we quantified the background signal for each image by measuring the mean *TCEB3* signal intensity in a 500 x 500-pixel region in tissue areas without oocytes (these areas often had DAPI+ nuclei and background fluorescence in the GFP channel but lacked *buc* signal). To perform background subtraction, we averaged the background signal for all fields of view imaged for an animal and subtracted this average background signal from the average oocyte *TCEB3* signal. The background-adjusted average oocyte *TCEB3* signal for each animal was normalized to the median of all *TCEB3^+/−^* animals and plotted. The quantification data were included in Extended Data Table 20.

### Cellular age estimation with human plasma proteomics data in the UK Biobank

#### Cohort description and plasma proteomics data source

Analyses of cellular aging in reproductive and sex-specific study populations were performed using UK Biobank human plasma proteomics data as a cross-sectional study. The UK Biobank (UKB) is a large prospective observational cohort that enrolled approximately 500,000 participants aged 40–69 years between 2006 and 2010. Participants underwent baseline assessment, including physical measurements, questionnaires, and biological sample collection, with longitudinal follow-up for disease outcomes and mortality extending over 15 years. Written informed consent was obtained from all participants enrolled in the cohort. In our study, UKB analyses were conducted using the UK Biobank Resource under Application Number 184058.

Baseline plasma proteomic profiling was performed in ∼54,000 UK Biobank participants as part of the UK Biobank Pharma Proteomics Project (UKB-PPP) using the Olink Explore 3072 platform. After UKB-PPP quality control, 2,923 proteins were available across ∼53,000 samples. Additional filtering excluded samples with more than 1,000 missing protein measurements (n = 8,182), samples with discordant reported and genetic sex (n = 373), and proteins missing in more than 20% of samples (n = 7). The final dataset included 44,458 participants and 2,916 proteins. Data were split by assessment center into separate training (n = 21,983) and test (n = 22,475) sets. Protein values were scaled to have a mean of zero and a standard deviation of 1 using the distribution of protein abundance values in the training cohort. Missing values (comprising 2.7% of the dataset) were imputed using k-nearest neighbors imputation (k = 148; *scikit-learn* KNNImputer v1.3.1). Imputation performance was evaluated by masked-value reconstruction, yielding a mean absolute error of 0.57.

#### Reproductive and baseline clinical phenotypes

The following UK Biobank data fields were used to define the study populations: ‘Sex’ (data field 31); ‘age first had sexual intercourse’ (data field 2139); ‘lifetime number of sexual partners’ (data field 2149); ‘ever had stillbirth, spontaneous miscarriage, or termination’ (data field 2774); ‘number of live births’ (data field 2734); and ‘number of previous pregnancies’ (data field 41267). We classified women into two groups based on their self-reported sexual histories using field 2139. We identified ‘Never had sex’ women as those who submitted a survey response ‘Never had sex’ in data field 2139. Individuals with ambiguous responses in field 2139 (i.e., ‘Prefer not to answer’ or ‘Do not know’) were excluded from the analysis. Additionally, individuals with conflicting answers (e.g., reported ‘Never had sex’ but also reported ‘Yes’ to data field 2774 or a positive integer value to data field 2734) were excluded from the analysis. To control for the effect of pregnancy and parenting, those who reported one or more live births (data field 2734) were excluded. Lastly, only individuals included in the UK Biobank Pharma Proteomics Project by random selection was included. Individual selected non-randomly by the UK Biobank Pharma Proteomics Project for inclusion due to disease status (data field 30903 ‘Yes) were excluded. Using these definitions, we identified 3,211 women in the ‘Had sex’ category and 165 women in the ‘Never had sex’ category.

Cohort-level demographics and participant disease characteristics were additionally extracted. Disease history was defined using UKB hospital inpatient ICD-10 diagnosis codes from data field p41270 and the corresponding first-recorded diagnosis dates from fields p41280_a0 through p41280_a258. ICD-10 codes were counted only if their first-recorded date was on or before the participant’s baseline assessment date. Body mass index (BMI; data field 21001) and ethnic background (data field 21000) were included as baseline demographic characteristics.

#### Cell type aging model development with plasma proteomics

Human cell-type biological age estimates were derived using previously developed Olink plasma proteomic cellular aging clock models (*129*). In these models, imputed plasma proteomic abundance values were applied to pretrained model weights, with ‘Sex’ included as a covariate. Further details regarding model development, cell-type plasma protein enrichment classification, and validation were reported by Ding *et al*., (*129*). The hepatocyte aging model consisted of 82 uniquely mapped cell-type-enriched plasma proteins that were measurable in the Olink 3072 assay (e.g., alpha-1-microglobulin/bikunin precursor, *AMBP*; serpin family C member 1, *SERPINC1*). The hepatocyte model was among the best-performing cell-type clocks in the original study, with training-set and testing-set Pearson correlations between predicted and actual age of 0.5524 and 0.5598, respectively, and corresponding mean absolute errors (MAEs) of 5.57 years and 5.62 years, respectively.

The cell-type-specific aging models were developed using ElasticNet regression implemented with the *glmnet* R package (v4.1.7). In brief, for each cell type, a distinct set of cell-type-enriched plasma proteins was used to predict chronological age, with ‘Sex’ included as an additional covariate, encoded as Female = 1 and Male = 0. Protein abundance values were log-transformed and z-scored before model fitting. During training, ElasticNet regression learned a weighted linear combination of cell-type-enriched protein features and the ‘Sex’ covariate to predict chronological age, while applying regularization to reduce overfitting. Models were fitted with an ElasticNet mixing parameter of α = 0.5, and the regularization parameter λ was selected by 10-fold cross-validation using *cv.glmnet* in the *glmnet* R package (v4.1.7). Predicted biological age was calculated by applying the trained model coefficients at *lambda.min* to each individual’s ‘Sex’ covariate and the corresponding z-scored protein abundance values. Models were trained in the UK Biobank training set (n = 21,983) and applied without refitting to the held-out UK Biobank test set (n = 22,475).

#### Cellular age estimation and age gap calculation

Consistent with the original publication (*129*), we quantified cell-type biological aging patterns among participants by computing age gaps. Age gaps represent the difference between an individual’s predicted biological age and the population-average predicted age expected for their chronological age. A positive age gap indicates that an individual’s cell type appears biologically older than that of the same-age peers. For each cell-type aging model, age gaps were calculated by fitting a Locally Weighted Scatterplot Smoothing (LOWESS) regression between predicted biological age and chronological age using the *lowess* function from the Python *statsmodels* package (v0.14.4; fraction = 2/3). Individual age gaps were defined as the residual between predicted biological age and the LOWESS-estimated expected age at the corresponding chronological age. Age gaps were then z-scored within each aging model across all 44,458 individuals in the training and test cohorts to enable comparison across cell types. These normalized age gaps were referred to as ‘biological age advancement.’

For each selected cell type, z-scored biological age gaps were compared between sexual history groups using two-sided Mann-Whitney *U* tests implemented with scipy.stats.mannwhitneyu in SciPy v1.13.0. Multiple hypothesis correction was performed using the Benjamini-Hochberg method and applied across the 28 cell type comparisons to control the false discovery rate using statsmodels.stats.multitest.multipletests with method=“fdr_bh” in statsmodels v0.14.6. Heatmap values represent group-mean, z-scored biological age gaps, while statistical significance reflects Mann-Whitney *U* tests comparing the distributions of age gaps between groups within each cell type. The prediction results were included in Extended Data Table 21.

### Statistical analysis

Statistics were performed in R or using GraphPad Prism (v.10) for all analyses. For microscopy, molecular assay analyses, biological age advancement comparisons, and demographic distributions, we used a two-sided non-parametric Mann-Whitney *U* test to compare two conditions, with each animal or individual as a data point. Two-way ANOVA was used to compare across multiple factors (e.g., Age, Mating status, Age x mating interaction) if the datasets and residuals were normally distributed (tested using the Shapiro-Wilk or Kolmogorov-Smirnov tests, *P* < 0.01). For analyses related to liver scRNA-seq and organ RNA-seq, individual genes were tested for significance between two conditions using a two-sided Wald test, followed by Benjamini-Hochberg (BH) multiple hypothesis correction. For lipidomics, individual lipid species were tested using a two-sided Welch’s t-test, followed by BH correction. For CellBiAge clocks, the *P* values were determined using a linear mixed-effects model for all pairwise comparisons, except a two-sided Welch’s t-test was used for the comparison between unmated young vs. unmated old. For acetaminophen treatment, a two-sided paired t-test (paired by cohort) was used to compare the proportion of unmated and mated females that survived past 48 hours. For microscopy analysis, outlier detection was performed in GraphPad using the Robust Regression and Outlier Removal (ROUT) method, with the most stringent setting (Q = 0.1%). Unless specified, the general criterion for statistical significance was set at *P* < 0.05.

---

**Fig. S1.**
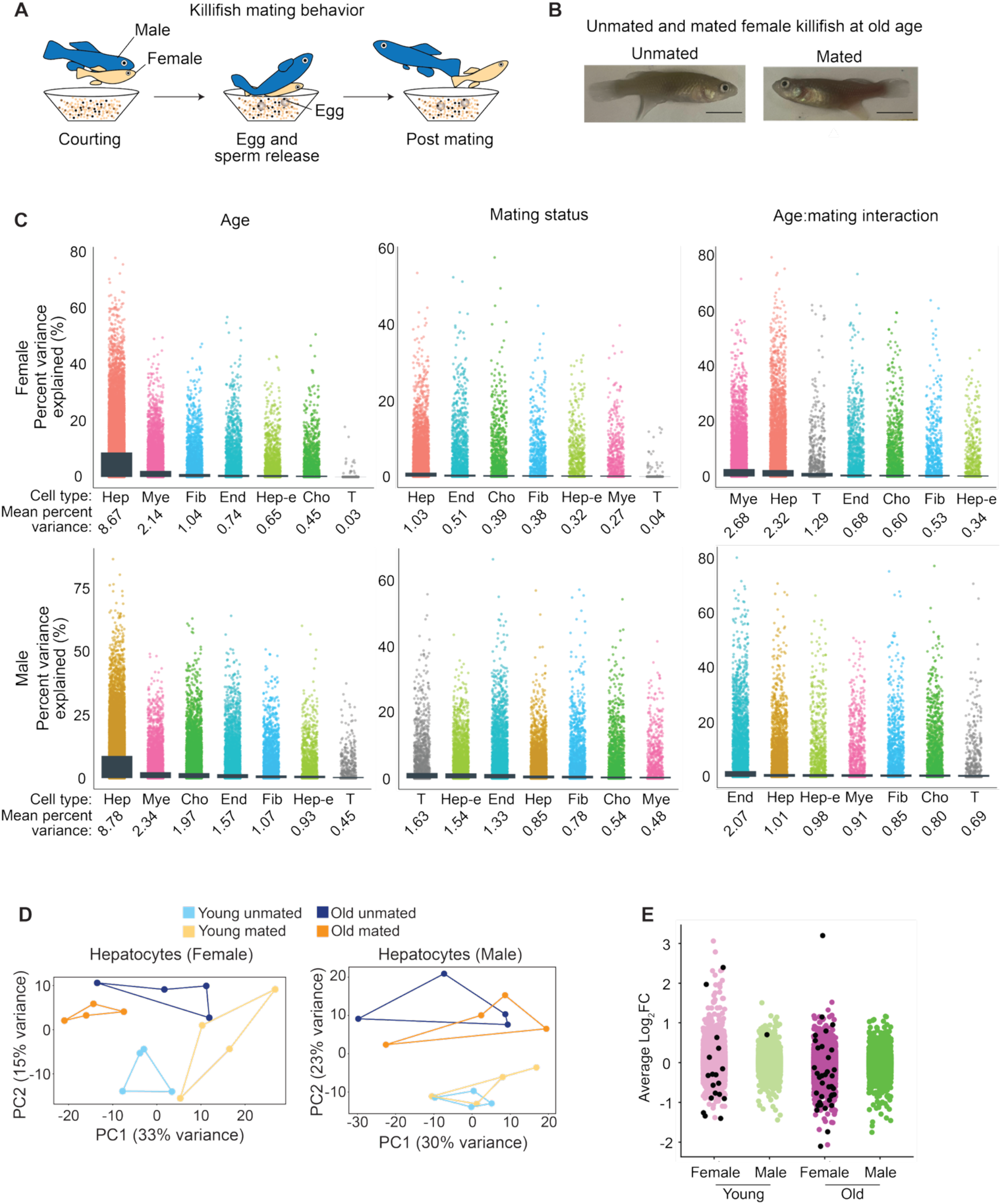
Aging and mating effects on cell type transcriptomes. (A) Schematic of killifish mating behavior. A male killifish courts a female, presses against her belly to stimulate the release of eggs onto a sandy surface, and then releases sperm to fertilize the eggs externally. The fertilized eggs are buried in the sand to develop externally. (B) Example images of unmated and mated 4.5-month-old killifish females. Scale bar, 1 cm. (C) Percent variance explained for age (left), mating status (middle), and age-mating interaction (right). Variance partition analysis was performed using the pseudobulked expression values of each gene in a cell type. Each dot represents the percent variance of a gene explained by each variable. The bar and the number under each cell type represent the mean percent variance across all genes in that cell type. Hep, hepatocytes; Mye, macrophages/neutrophils; Fib, fibroblasts; End, endothelial cells; Hep-e, hepatocytes-efferocytosing; Cho, cholangiocytes; T, T cells. (D) Principal Component Analysis (PCA) performed using the pseudobulked transcriptomes of female hepatocytes or male hepatocytes. Each dot is an animal. (E) Differentially expressed genes between unmated and mated animals at young or old age. Each dot represents a gene, plotting the pseudobulked Log_2_ fold change (Log_2_FC) comparing unmated and mated animals at each age and sex. The pseudobulked Log_2_FC is the average Log_2_FC across all hepatocytes for each age and sex. A positive Log_2_FC value means higher expression in the mated condition. Black dots, genes significantly changed by mating (false discovery rate, FDR ≤ 0.05). Statistics were performed using a Wald test, followed by Benjamini-Hochberg multiple hypothesis correction.

**Fig. S2.**
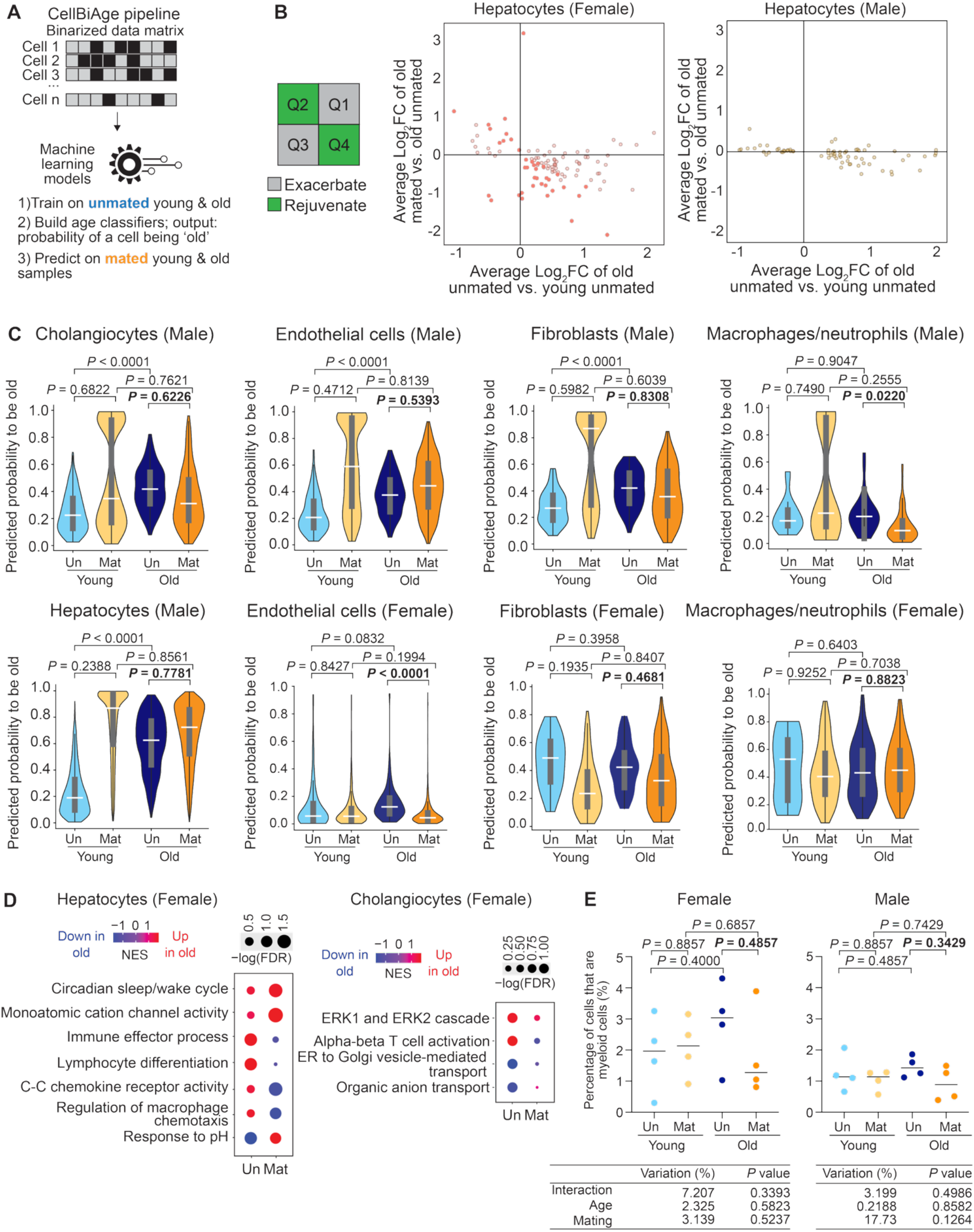
Effect of mating on age prediction on different liver cell types in females and males. (A) Schematic of the CellBiAge pipeline. (B) Correlation analysis comparing the age-related changes versus mating-related changes. Each dot is a gene. The pseudobulked expression of each gene (average gene expression in each cell type), for either female or male hepatocytes, was used to calculate the Log_2_ fold change (Log_2_FC) between old mated and old unmated (old unmated as the reference, ‘mating axis’) and between old unmated and young unmated (young unmated as the reference, ‘age axis’). Filled color, significantly changed for the mating axis. Q1, quadrant 1; Q2, quadrant 2; Q3, quadrant 3; Q4, quadrant 4. A gene located in Q1 or Q3 represents one for which mating ‘exacerbates’ the changes observed in aging. In contrast, for a gene located in Q2 or Q4, mating ‘reverses’ or ‘rejuvenates’ the changes observed in aging. (C) Distribution of each cell’s CellBiAge predicted probability of being ‘old,’ plotted for male and female liver cell types. Bar represents the median of all cells (grouping all fish of each condition), and the quantiles 1 (Q1) and 3 (Q3) are the 25^th^ and 75^th^ percentiles, respectively. The *P* values were determined using a linear mixed-effects model for all pairwise comparisons, except the comparison between unmated young vs. unmated old, where a Welch’s t-test was used. Un, unmated; Mat, mated. (D) Gene set enrichment analysis (GSEA) results for female hepatocytes (left) and cholangiocytes (right), identifying the pathways significantly enriched for the genes upregulated or downregulated with age in mated or unmated females. NES, normalized enrichment score. Dot size, –log_10_ of the false discovery rate (FDR). (E) Proportion of myeloid cells (macrophages/neutrophils) detected in scRNA-seq for female and male livers. Each dot is an animal (young unmated, n = 4 fish; young mated, n = 4 fish; old unmated, n = 4 fish; old mated, n = 4 fish), plotting the percentage of total cells that belong to the myeloid cell cluster. Each bar is the median of the four fish per condition. The *P* values were determined using the Mann-Whitney *U* test for pairwise comparisons and a two-way ANOVA to assess the effects of age, mating status, and the age-mating interaction.

**Fig. S3.**
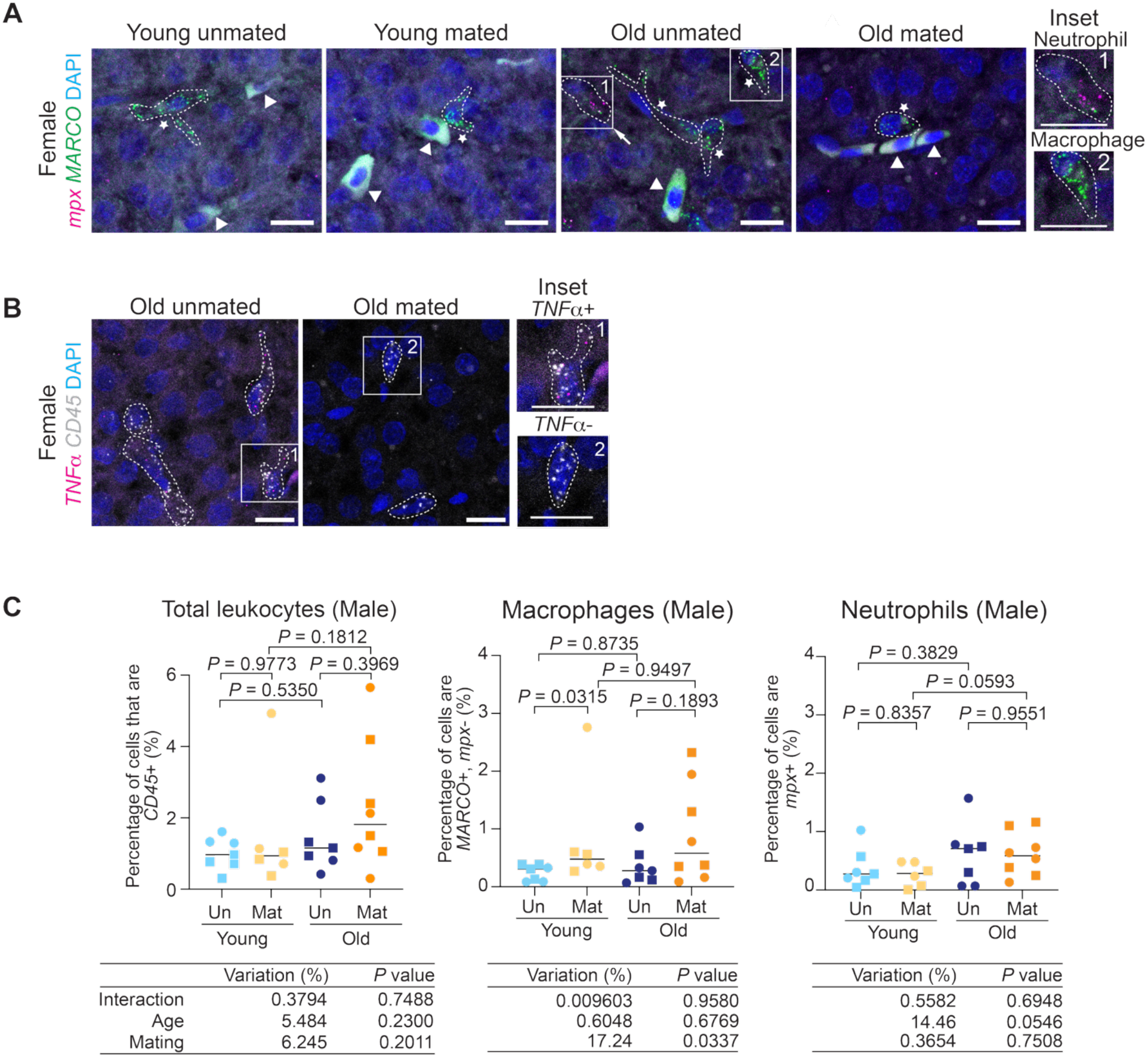
Mating effects on liver immune cell levels in young and old killifish. (A) Example RNA *in situ* images for female liver sections (young unmated, n = 7 fish; young mated, n = 6 fish; old unmated, n = 7 fish; old mated, n = 7 fish), staining for *mpx* (magenta) and *MARCO* (green) mRNAs and nuclei by DAPI (blue). Animals were raised in 2 independent cohorts, separate from the scRNA-seq cohorts. Hereinafter, in this figure, young, 2-2.5 months old; old, 4.5 months old. Dashed lines outline cells. Star, a macrophage (*MARCO*+, *mpx*-); arrow, a neutrophil (*mpx+*); arrowhead, a red blood cell, which is autofluorescent. Scale bar, 10 *µm*. All images were captured and displayed under the same settings. Inset images are also shown in Fig. 2A. (B) Example RNA *in situ* images for female liver sections (young unmated, n = 11 fish; young mated, n = 10 fish; old unmated, n = 10 fish; old mated, n = 11 fish), staining for *TNFα* (magenta) and *CD45* (gray) mRNAs and nuclei by DAPI (blue). Animals were pooled from the scRNA-seq cohorts and 2 additional cohorts independent of the scRNA-seq cohorts. Dashed lines outline cells. Scale bar, 10 *µm*. All images were captured and displayed under the same settings. Inset images are also shown in Fig. 2C. (C) Proportion of different immune cells in male livers, including total leukocytes (*CD45*+), macrophages (*MARCO*+, *mpx*-), and neutrophils (*mpx*+). Each dot represents an individual fish (young unmated, n = 7 fish; young mated, n = 6 fish; old unmated, n = 7 fish; old mated, n = 8 fish), and the bar is the median of all fish in each condition. Symbol shape, cohort number (animals were raised in 2 cohorts independent of the scRNA-seq cohorts). The *P* values were determined using the Mann-Whitney *U* test for pairwise comparisons and a two-way ANOVA to assess the effects of age, mating status, and age-mating interaction. Un, unmated; Mat, mated.

**Fig. S4.**
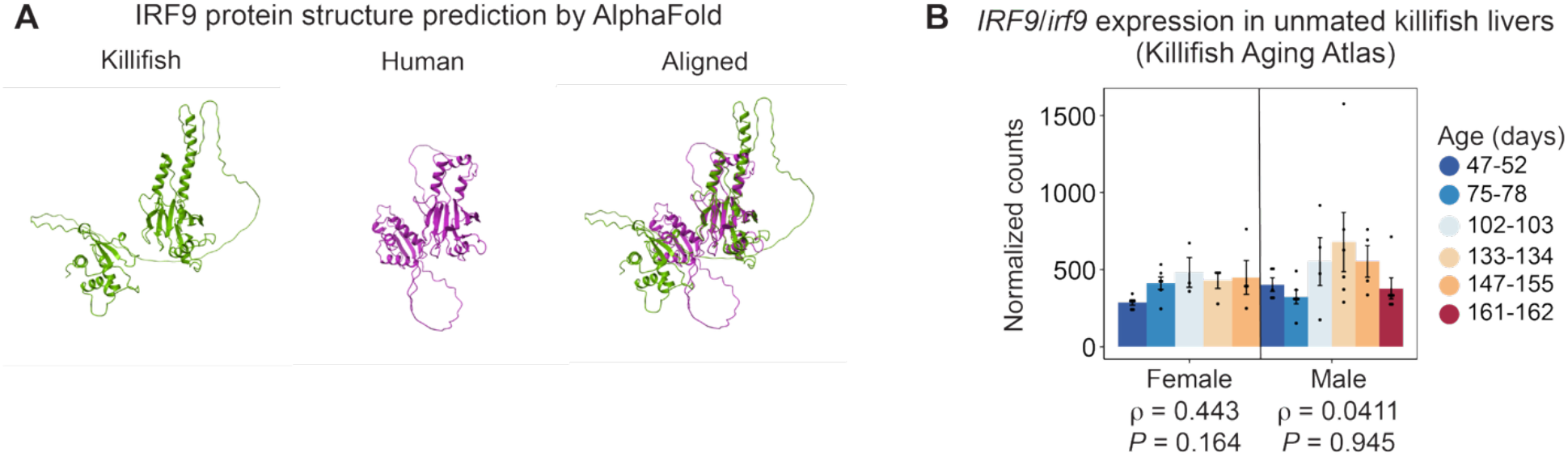
IRF9 protein structure and expression during aging. (A) IRF9 protein structure prediction using AlphaFold for the killifish (left) and human (middle) protein sequences. The killifish IRF9 protein had ∼33% sequence identity to the human protein by alignment and shared structural overlap with the human protein (right). (B) DESeq2-normalized RNA-seq expression of the *IRF9* (killifish name: *irf9*) gene in the liver of unmated fish. Data were derived from the Killifish Aging Atlas (*60*). Each dot is the expression value from an animal. Bar height represents the mean expression for each sex and age, and the error bars are the standard errors of the mean calculated across the biological replicates within each group. The Spearman’s rank correlation (*ρ*) across all age groups and the adjusted *P* values (after Benjamin-Hochberg multiple hypothesis correction) were calculated using female-only or male-only transcriptomes. A positive *ρ* indicates upregulation with age, and a negative *ρ* indicates downregulation with age.

**Fig. S5.**
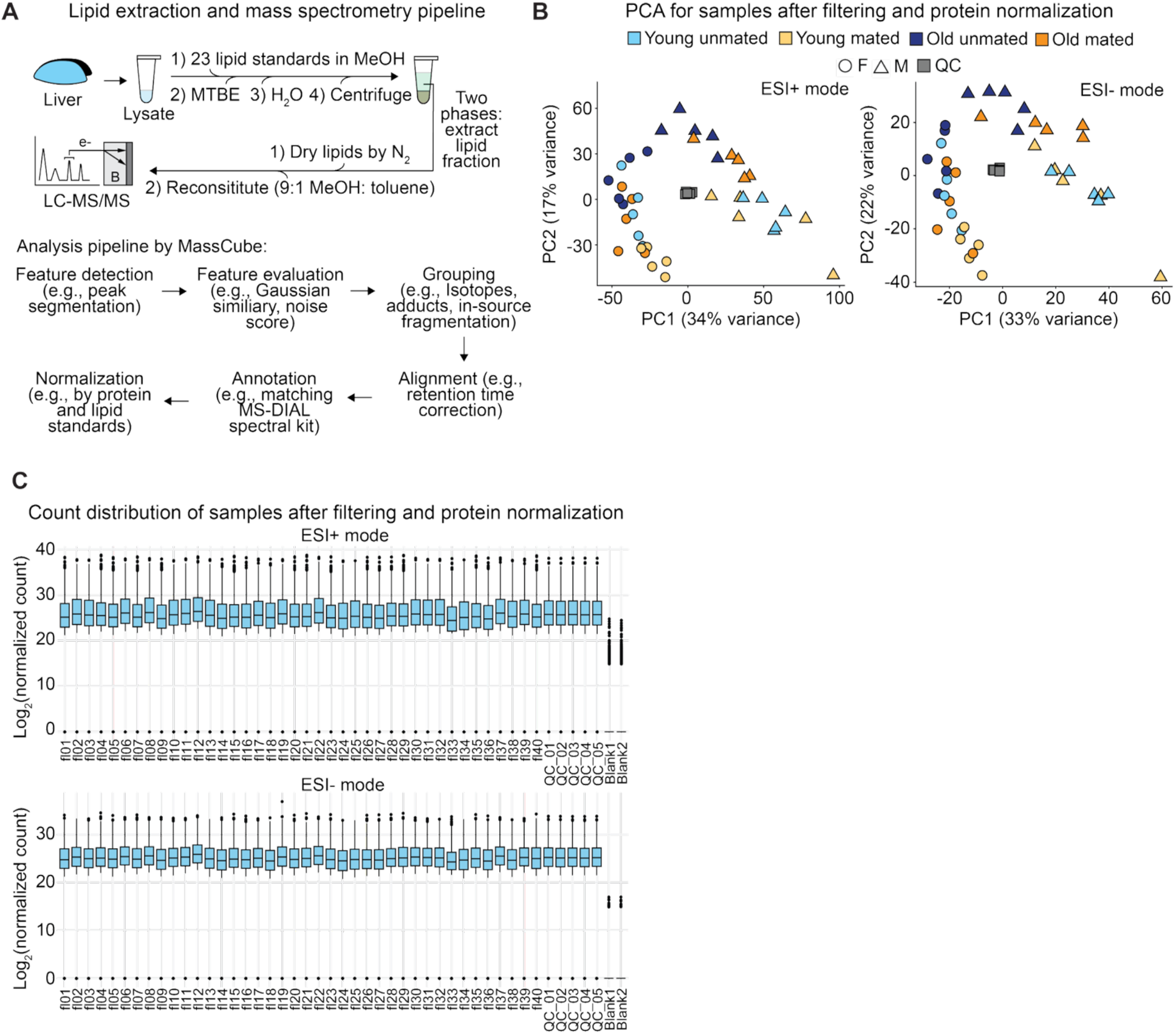
Quality control of the liver untargeted lipidomics dataset. (A) Schematic of the lipidomics pipeline. After lysis, lipids from each liver sample were extracted with methanol (MeOH) and MTBE, with 23 lipid standards spiked in before extraction for quantification. Liquid chromatography-tandem mass spectrometry (LC-MS/MS) was performed for each liver sample and quality-control samples, followed by analysis using the MassCube pipeline (see Methods for details). (B) PCA for all samples after quality filtering and protein normalization (both annotated and unannotated lipids are included). Left, data from the positive ionization mode (ESI+, 4,262 lipids plotted); right, data from the negative ionization mode (ESI-, 1,459 lipids plotted). Each dot is a fish (n = 5 fish per condition). Circle, female (F); triangle, male (M); square, quality control (QC) samples. (C) Log_2_ of the lipid abundance of all samples after quality filtering and protein normalization (both annotated and unannotated lipids are included). Top, data from the positive ionization mode (ESI+, 4,262 lipids plotted); bottom, data from the negative ionization mode (ESI-, 1,459 lipids plotted.

**Fig. S6.**
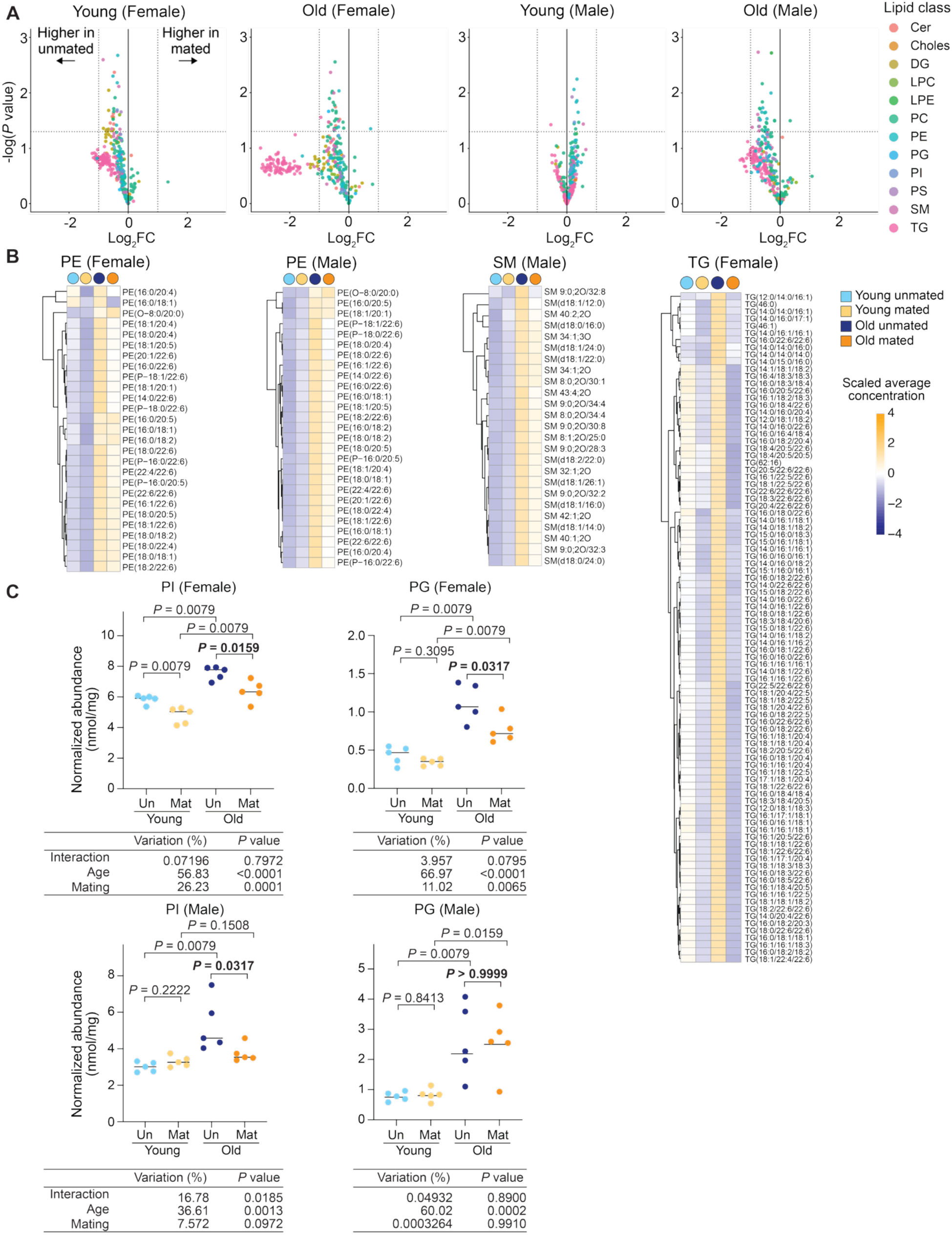
Mating affects liver lipid abundance in both males and females. (A) Volcano plot graphing the -log_10_ of the raw *P* value and the Log_2_ fold change (Log_2_FC) of mated vs. unmated samples at each age and sex. Each dot represents a uniquely annotated lipid species, color-coded by its lipid class. Vertical dotted lines correspond to Log_2_FC values of −1 and 1, respectively. Horizontal dotted line corresponds to a raw *P* value of 0.05. Note that the *P* values plotted were raw *P* values, and almost no individual lipid species were significantly affected by mating at the false discovery rate (FDR) of 0.05. Cer, ceramide; Choles, cholesterol; DG, diglyceride; LPC, lyso-phosphatidylcholine; LPE, lyso-phosphatidylethanolamine; PC, phosphatidylcholine; PE, phosphatidylethanolamine; PG, phosphatidylglycerol; PI, phosphatidylinositol; PS, phosphatidylserine; SM, sphingomyelin; TG, triglyceride. (B) Heatmaps of selected lipid classes in male or female livers. The concentration of each detected lipid species within a given lipid class was averaged across the five animals per condition and then z-scaled across conditions. PE, phosphatidylethanolamine; SM, sphingomyelin; TG, triglyceride. (C) The abundance of the phosphatidylinositol (PI) and phosphatidylglycerol (PG) lipid classes in male and female livers. Each dot is a fish (young unmated, n = 5; young mated, n = 5; old unmated, n = 5; old mated, n = 5) (raised from 3 independent cohorts), plotting the normalized abundance summed across all detected lipid species belonging to a given lipid class. The bar is the median of all fish in each condition. Young, 2-2.5 months old; old, 4.5 months old. Un, unmated; Mat, mated. The *P* values were determined using the Mann-Whitney *U* test for pairwise comparisons and a two-way ANOVA to assess the effects of age, mating status, and the age-mating interaction.

**Fig. S7.**
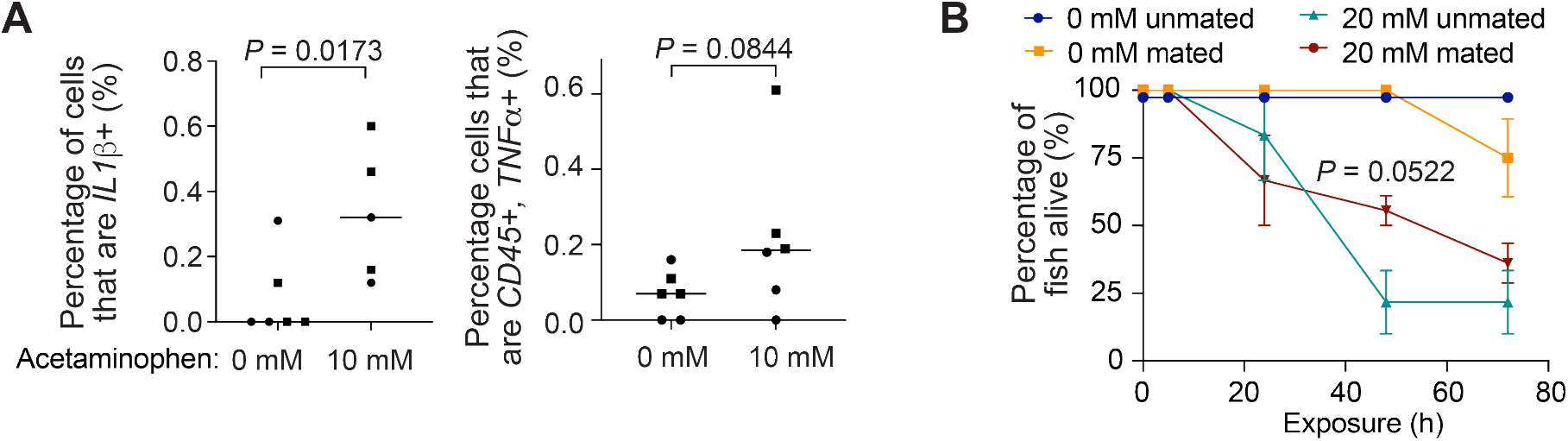
Acetaminophen induces inflammation and lethality in females. (A) Percentage of cells expressing pro-inflammatory marker *IL1β* (left) and percentage of *TNFα*+ leukocytes (*CD45*+, *TNFα*+) (right) in the livers of unmated 3-month-old females treated with 0 mM or 10 mM acetaminophen for 72 hours. Each dot represents an individual fish (0 mM, n = 6 fish; 10 mM, n = 6 fish), and the bar is the median of all fish per treatment group. Symbol shape, cohort number (animals were raised in 2 independent cohorts). The *P* values were determined using the Mann-Whitney *U* test for pairwise comparisons. For *IL1β* staining (left), an outlier was detected for the 10 mM condition and excluded. (B) Lethality curve for unmated old females (0 mM, n = 9 fish; 20 mM, n = 13 fish at the start of treatment; pooled from three independent cohorts) and mated old females (0 mM, n = 10 fish; 20 mM, n = 14 fish at the start of treatment; pooled from three independent cohorts) treated with 0 mM or 20 mM acetaminophen for 72 hours. Each dot represents the average survival percentage across 3 independent cohorts, and the error bar represents the standard error of the mean. The *P* value was determined using a paired t-test comparing unmated and mated females treated with 20 mM acetaminophen, matching by cohort number.

**Fig. S8.**
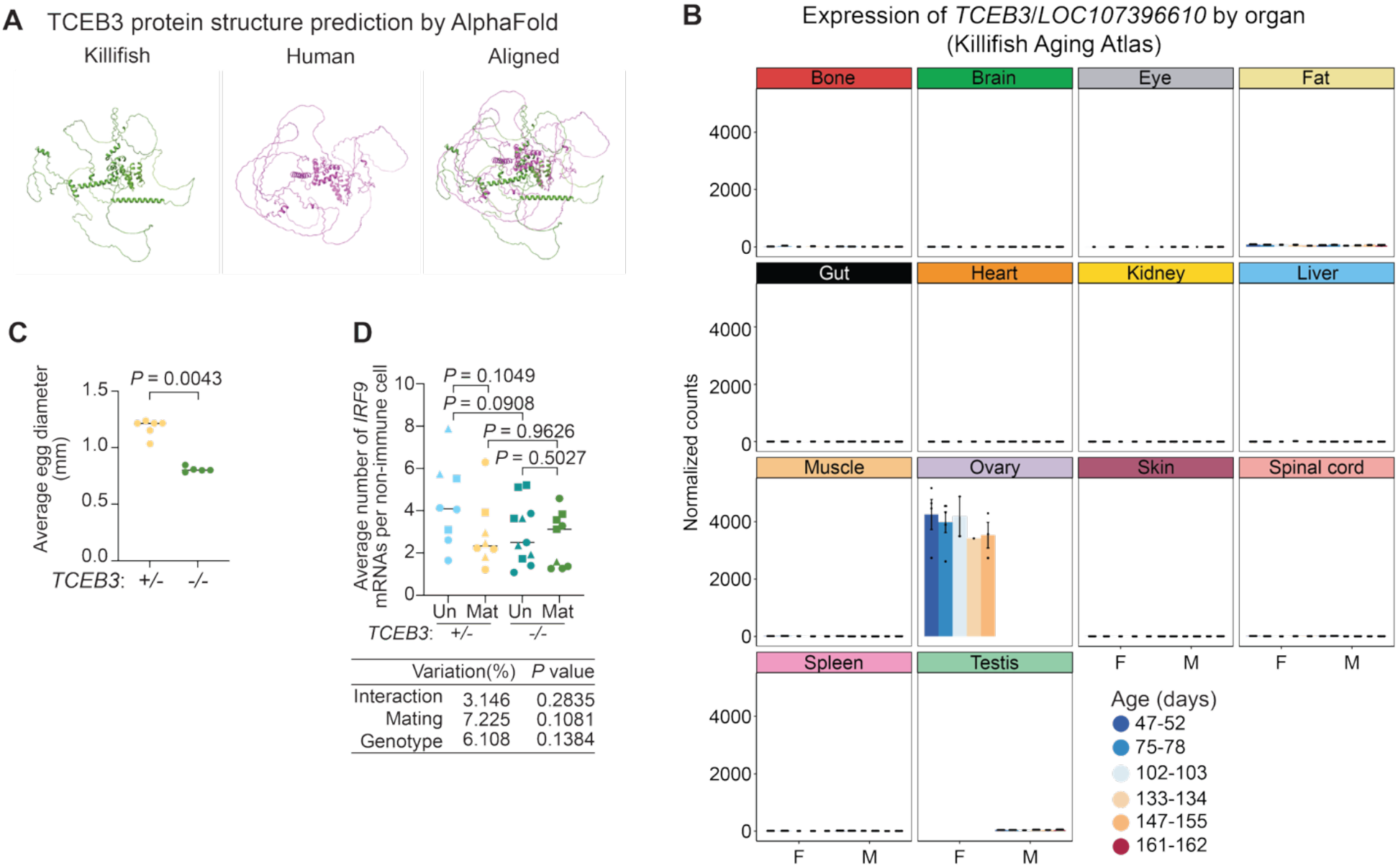
*TCEB3*, an ovarian-specific gene, is necessary for oocyte maturation. (A) TCEB3 protein structure prediction using AlphaFold for the killifish (left) and human (middle). The killifish TCEB3 protein had ∼40% protein sequence identity to the human protein by alignment and shared structural overlap with the human protein (right). (B) DESeq2-normalized RNA-seq expression of the *TCEB3* (killifish name: *LOC107396610*) gene in different organs of unmated fish. Data were derived from the Killifish Aging Atlas (*60*). Each dot is the expression value of an animal. Bar height represents the mean expression for each sex and age, and the error bars are the standard errors of the mean calculated across the biological replicates within each group. F, female; M, male. (C) Egg size of the *TCEB3* heterozygous (*TCEB3^+/−^*) and homozygous (*TCEB3^+/+^*) mutants, plotting the average diameter. The *P* values were determined using the Mann-Whitney *U* test for pairwise comparison. (D) Average expression of *IRF9* in the non-immune (*CD45*-negative) cells of *TCEB3* mutant female livers. Each dot represents an individual fish (unmated *TCEB3^+/−^*, n = 8 fish; mated *TCEB3^+/−^*, n = 8 fish; unmated *TCEB3^−/−^*, n = 11 fish; mated *TCEB3^−/−^*, n = 9 fish), and the bar is the median of all fish per condition. Symbol shape, cohort number (animals were raised in 3 independent cohorts). The *P* values were determined using the Mann-Whitney *U* test for pairwise comparisons and a two-way ANOVA to assess the effects of age, mating status, and the age-mating interaction.

**Fig. S9.**
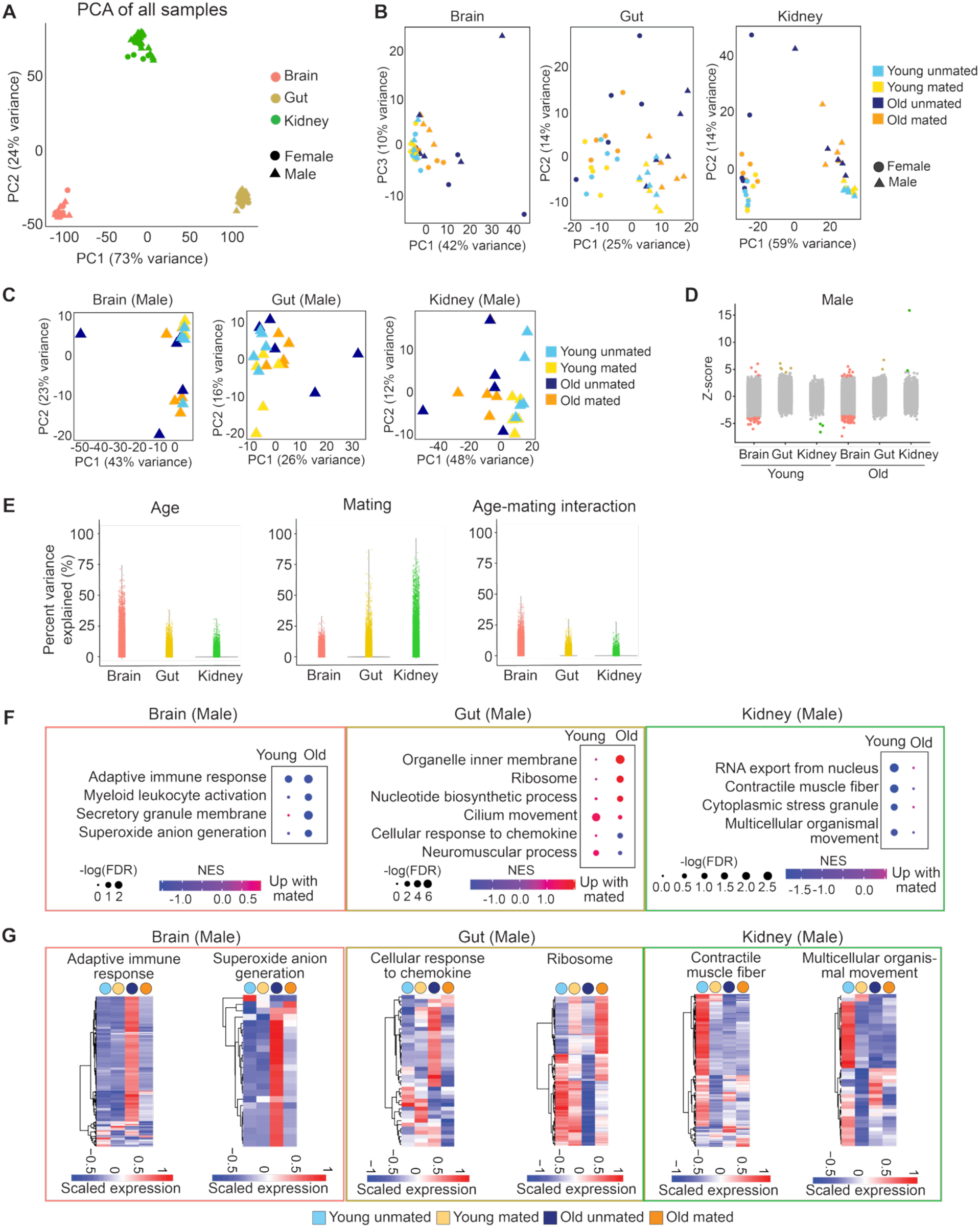
Mating impacts the aging transcriptomes of other organs. (A) Principal Component Analysis (PCA) of the transcriptomes of all organ samples, with samples coded by sex and organ identity. Raw counts were transformed using variance stabilization and used as PCA input. Each dot represents the full transcriptome of one individual fish organ. (B) Organ-specific PCA for the brain, gut, and head kidney. Five fish (n = 5 fish per condition; animals were raised across 3 independent cohorts) were used for each condition (combination of sex, age, and mating status). Hereinafter, in this figure, young, 2-2.5 months old; old, 4.5 months old. (C) PCA for the male brain, gut, and head kidney. Five fish (n = 5 fish per condition; animals were raised across 3 independent cohorts) were used for each condition (combination of age and mating status). (D) Dot plot of the differentially expressed genes by mating at each age in male organs, graphing z-scores. Colored, significantly upregulated or downregulated in mated animals (false discovery rate, FDR ≤ 0.05); gray, not significantly altered (FDR > 0.05). Statistics were performed using a Wald test, followed by Benjamini-Hochberg multiple hypothesis correction. (E) Percent variance explained for age (left), mating status (middle), and age-mating interaction (right). Variance partition analysis was performed using the raw count of each gene in each organ, with both sexes analyzed together. Each dot represents the percent variance of a gene explained by a given variable. (F) Gene set enrichment analysis (GSEA) results for each male organ, identifying the pathways significantly enriched for the genes upregulated or downregulated by mating in young or old animals. NES, normalized enrichment score. Dot size, –log_10_ of the false discovery rate (FDR). (G) Heatmaps of the genes that drive each selected gene ontology (GO) term for each male organ. For each gene, its DESeq2-normalized expression is averaged from the 5 fish for each condition and z-scaled.

**Fig. S10.**
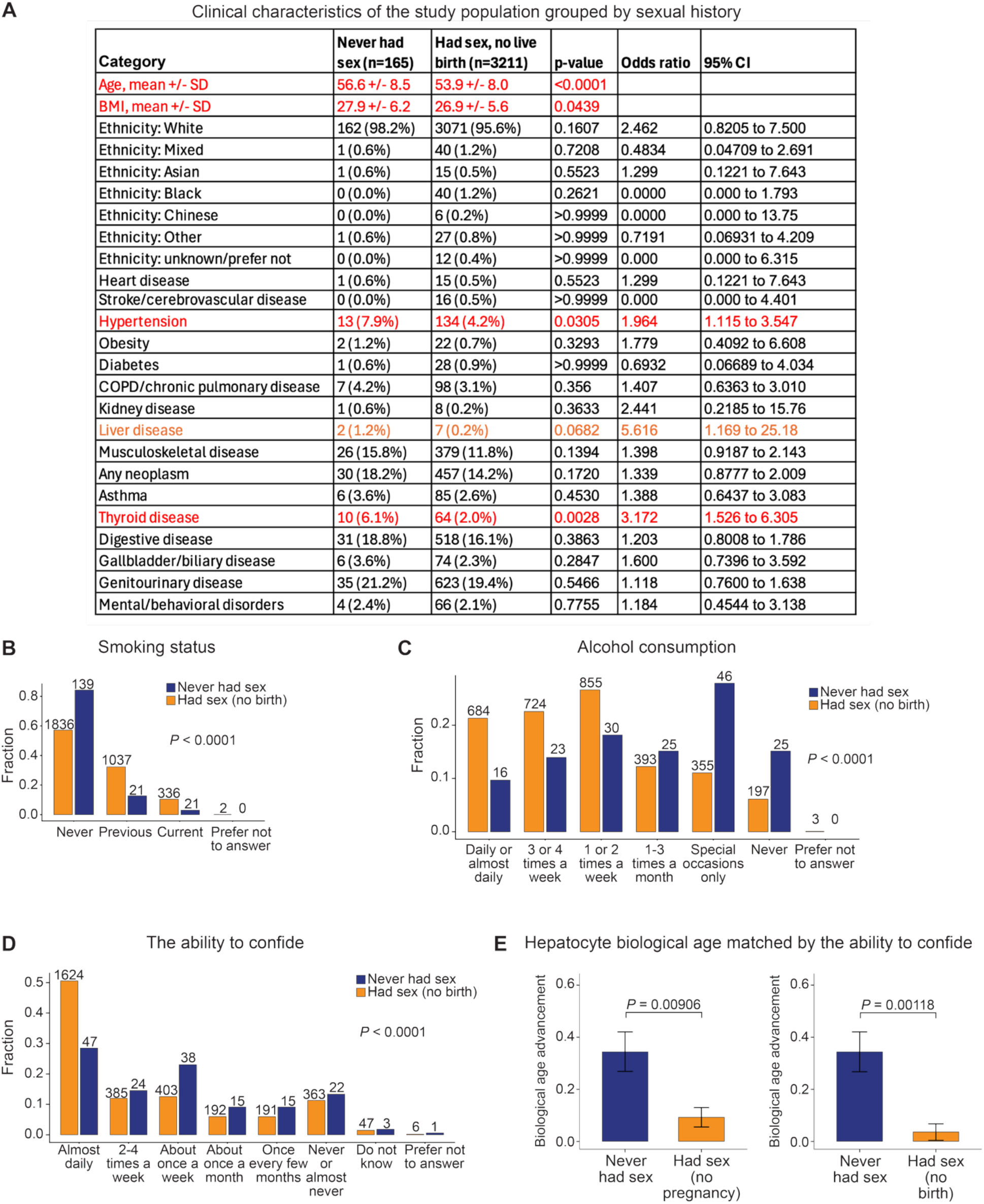
Characteristics of UK Biobank women stratified by sexual history. (A) Clinical characteristics of the study population. Women were classified into two groups: those who reported an age at first sexual intercourse (‘Had sex’) and those who did not (‘Never had sex,’ n = 165). We further limited our analysis to women who had never given birth (n = 3,211). Disease categories were derived from UK Biobank hospital inpatient ICD-10 diagnosis codes and grouped into prespecified clinically related categories (Data S10). First-recorded diagnosis dates were used to determine baseline disease status; codes were counted only if recorded on or before the participant’s baseline assessment visit. Data are presented as counts, with age reported in years and BMI in kg/m^2^, respectively. For age and BMI, the mean and standard deviation are listed, and a Welch t-test was used to compare the two groups of women. For all other categories, the number of individuals and the corresponding proportion (indicated in parentheses) in each category is listed, and Fisher’s exact test was used to compare the two groups of women. CI, confidence interval. In the red categories, the two groups of women were statistically different at the *P* < 0.05 threshold; in the orange category, at a *P* value slightly higher than 0.05. (B, C) Distributions of smoking status (panel B) and alcohol consumption (panel C) for women who ‘Never had sex’ (n = 165) or ‘Had sex’ but never given birth (n = 3,211). The number above each bar indicates the number of individuals in a given category. Bar height corresponds to the fraction of each category. Statistics were performed using the Mann-Whitney *U* test, excluding the ‘Do not know’ and ‘Preferred not to answer’ group. (D) Distribution of the ability to confide (an approximate metric of loneliness) for women who ‘Never had sex’ (n = 165) or ‘Had sex’ but had never given birth (n = 3,211). Graphics and statistics were generated as in panel B, excluding the ‘Do not know’ and ‘Preferred not to answer’ groups. (E) Hepatocyte biological age advancement calculated for women who ‘Never had sex’ (n = 165) or ‘Had sex’ but had never been pregnant (‘no pregnancy,’ n = 660) or have never given birth (‘no birth,’ n = 990), after matching the distributions based on the ability to confide in someone. Matching was performed by selecting, for each participant in the “Never had sex” group, 6 random individuals (for ‘no birth’) or 4 random individuals (for ‘no pregnancy’) from the ‘Had sex’ group who belong to the same category in the distribution in panel D (the categories include: almost daily, 2-4 times a week, about once a week, about once a month, once every few months, never or almost never, do not know, or prefer not to answer). The ratio was chosen to match the minimum ratio between the ‘Never had sex’ and the ‘Had sex’ populations stratified by the categories in panel D. Bar height corresponds to the mean of the age gaps within each group, and the error bar corresponds to ± standard error of the mean. Statistics were performed using the Mann-Whitney *U* test.

## Supplementary Text

### Additional results for the CellBiAge classifiers for liver single-cell age prediction

We have included all statistical test results in Extended Data Table 3. For female livers, **Fig. 1F** and **fig. S2C** shows the results for combination 4, where three cell types (female hepatocytes, cholangiocytes, and endothelial cells) reached statistical significance (*P* < 0.05) in the mixed-effects model when comparing mated and unmated females at old age. The coefficients for the effect of mating on the predicted probability, and their corresponding confidence intervals, were negative for both cell types, suggesting a ‘transcriptomic rejuvenation’ effect of mating. Female cholangiocytes had the highest effect size (coefficient = −0.360; 95% confidence interval or CI [−0.459, −0.261]; *P* < 0.0001), followed by female hepatocytes (coefficient = −0.136; 95% CI [−0.239, −0.0327]; *P* = 0.0098) and female endothelial cells (coefficient = −0.0801; 95% CI [−0.0986, −0.0616]; *P* < 0.0001). In the other combinations (excluding combination 3 due to poor model performance), these cell types generally showed the same trend but did not reach statistical significance, likely due to the stringent statistical method used to account for within-animal dependency of single cells. For example, for female hepatocytes, the coefficients were negative for both combination 1 (coefficient = −0.186; 95% CI [−0.923, 0.550], *P* = 0.620) and combination 2 (coefficient = −0.352; 95% CI [−0.844, 0.140], *P* = 0.551). Female fibroblasts showed a negative coefficient, though not reaching a statistical significance of *P* < 0.05 (combinations 1, 2, and 4). Macrophages/neutrophils showed inconsistent signs of the mean confidence intervals across combinations 1, 2, and 4, probably due to the limited number of cells for model training; thus, we could not conclude on whether mating ‘rejuvenated’ or ‘exacerbated’ the aged transcriptomes.

For male livers, **fig. S2C** reported the results from combination 4. For male hepatocytes, this combination shared the same trend as combination 1 (coefficient = 0.049; 95% CI [−0.291, 0.389]; *P* = 0.628), suggesting that mating led to ‘older’ transcriptomic signatures for the hepatocytes of old males. However, combination 2 had a negative coefficient for the mating effect, −0.324 (95% CI [−0.527, −0.121]; *P* < 0.0001), which was inconsistent with combinations 1 and 4. The inconsistent signs of the coefficients among all the combinations made it difficult to conclude whether mating ‘rejuvenated’ or ‘exacerbated’ the aged transcriptomes. Such inconsistency was also observed for male efferocytosing hepatocytes, cholangiocytes, endothelial cells, and macrophages/neutrophils. Male fibroblasts showed a negative coefficient, though not reaching a statistical significance of *P* < 0.05, for all four combinations.

At a young age, except for one cell type in each sex, all other cell types (in all combinations) did not significantly differ between the mated and unmated fish, for both males and females. Female fibroblasts had a significant positive coefficient in combination 1 (coefficient = 0.166; 95% CI [0.0386, 0.293], *P* = 0.0106) but a negative coefficient in combination 4 (coefficient = −0.143; 95% CI [−0.359, 0.073], *P* = 0.1934) that did not reach a statistical significance of *P* < 0.05. Male endothelial cells had a positive coefficient in three of the four combinations (except combination 3: coefficient = −0.098; 95% CI [−0.278, 0.082]; *P* = 0.285). Combination 2 reached a statistical significance of *P* < 0.05 (coefficient = 0.232; 95% CI [0.080, 0.384]; *P* = 0.0028), but combination 1 (coefficient = 0.056; 95% CI [−0.189, 0.300]; *P* = 0.656) and combination 4 (coefficient = 0.269; 95% CI [−0.462, 0.100]; *P* = 0.471) did not. Given the inconsistent signs of the coefficients, we were unable to draw clear conclusions about the mating effects on any cell type at a young age. We provided all the probability distribution plots (cell-type models, all-cell models, and plots split by each animal) in Extended Data Table 7.

### Bootstrapping results

To test whether the unequal sample sizes between the ‘Never had sex’ and ‘Had sex’ (no pregnancy) groups drive the differences in hepatocyte age estimation, we performed bootstrapping analysis. Specifically, we held the ‘Never had sex’ group fixed, n = 165. In each of 10,000 iterations, controls were sampled randomly in a 3:1 ratio within the same ‘ability to confide’ strata as each ‘Never had sex’ individual, so that 495 individuals were sampled without replacement from the ‘Had sex’ (no pregnancy) pool. For each iteration, the difference in group means and the *P* values by Mann-Whitney *U* test were computed (**fig. S11**). Bootstrapping results suggested that the differences in hepatocyte age estimation between the ‘Never had sex’ and ‘Had sex’ (no pregnancy) groups were stable when we reduced the sample size difference between the two groups.

**Fig. S11.**
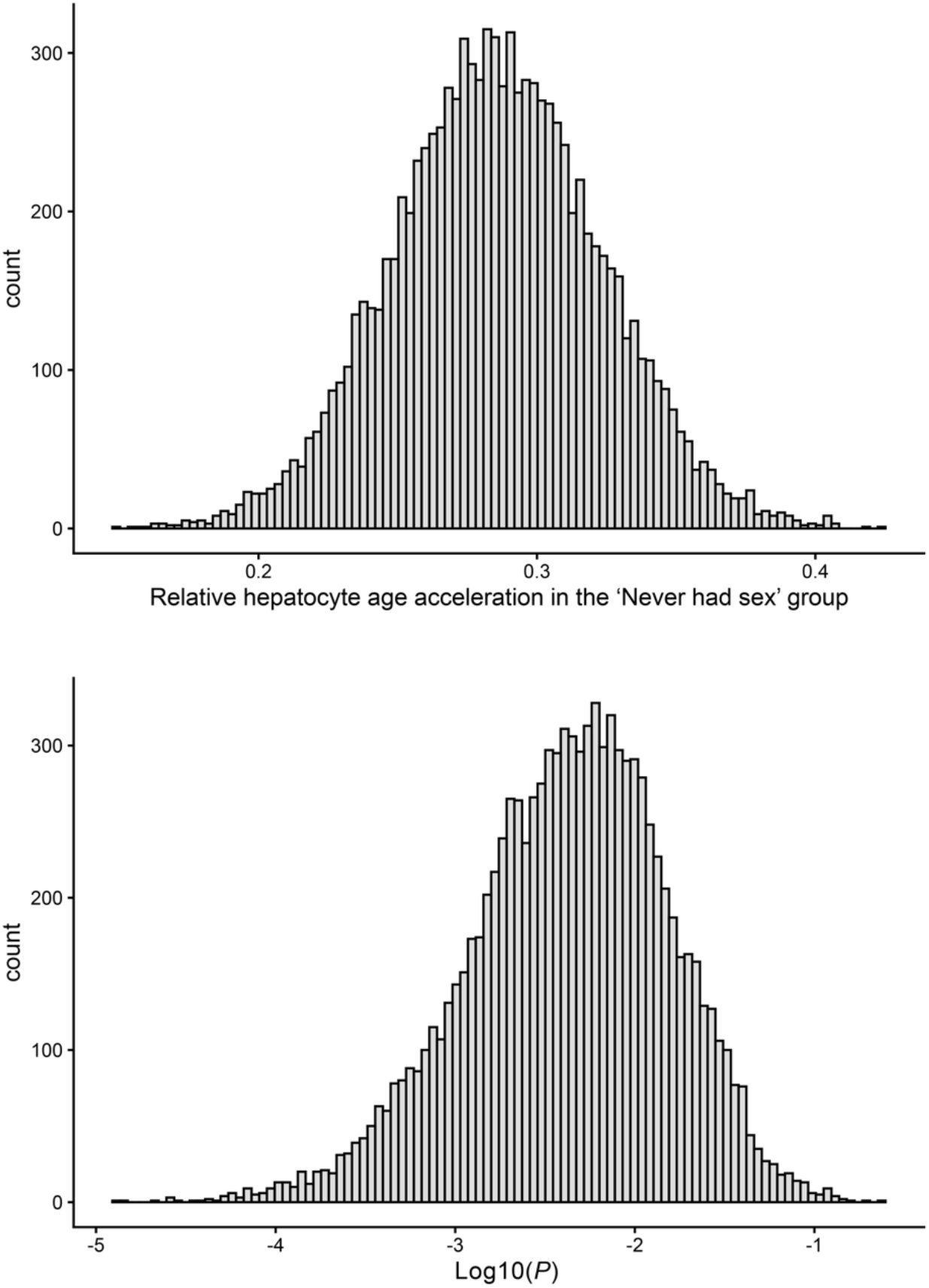
Bootstrapping results. The empirical distributions of the mean differences (top) and *P* values (bottom) across all iterations were plotted as histograms.

## Notes

### Competing Interest Statement

The authors have declared no competing interest.

